# Endocytosis at the mouse blood brain barrier is elevated during sleep

**DOI:** 10.64898/2026.04.02.716170

**Authors:** Fu Li, Dechun Chen, Amita Sehgal

## Abstract

Sleep is thought to be important for the clearance of brain waste, but exactly how it does so is still debated. Here, we demonstrate that endocytosis in brain endothelial cells (BECs) of the blood–brain barrier (BBB) is enhanced during sleep, facilitating the removal of brain-derived waste, including amyloid-β, into the circulation. Using proteomics, in vivo tracer imaging, and endothelial-specific genetic perturbations in mice, we demonstrate that sleep enhances endocytic vesicle formation and cargo transcytosis in brain endothelial cells (BECs). Conversely, blocking endocytosis through endothelial Dnm2 knockout suppresses BEC-mediated transport and elevates sleep need, revealing a causal feedback loop between sleep and vascular endocytosis. These findings identify BBB endocytosis as a key sleep-dependent clearance pathway with implications for neurodegenerative disease.

## Main

Sleep is associated with improved cognition, synaptic plasticity and overall fitness. On a cellular/molecular level, work in Drosophila indicates that sleep maintains the integrity of neuronal mitochondria, at least in part by promoting the transfer of oxidative damage from neurons to glia^1^. Other clearance functions attributed to Drosophila sleep, such as autophagy and mobilization of lipids through the BBB, may also serve this goal^2^ ^3^. Indeed, efficient clearance of metabolic byproducts and misfolded proteins from the brain is critical for neuronal health across species. Impairment of waste clearance in mammals contributes to neurodegenerative diseases such as Alzheimer’s disease, in which amyloid-β (Aβ) and tau accumulate in the parenchyma^4^. Glymphatic flow and meningeal lymphatic drainage have been implicated in clearance during sleep^5^, but participation of the blood–brain barrier (BBB) remains unclear even though this clearly offers another mode of transport from the brain. Other than circadian and wake modulation of efflux transporters^6,7^, regulation of the mammalian BBB has not been linked to sleep: wake states.

Hypothesizing that the content and function of the BBB endothelium are modulated by sleep, we first sought to determine whether the protein profile of the endothelium is affected by sleep. We subjected one group of mice to 6 hours of sleep deprivation (SD), starting at Zeitgeber Time (ZT0) or lights on, followed by 2 hours of recovery sleep, whereas a second group mice underwent 8 hours of SD (Fig.1a). Thus, they were collected at the same circadian time, and brain endothelial cells (BECs) were isolated from both groups. As we were particularly interested in proteins trafficked through the BBB, we extracted endosomal vesicles from these cells and analyzed proteins in these by Liquid Chromatography (LC) and a tandem mass spectrometer (MS/MS) method on QE-HF in DIA mode. A total of 2043 proteins were detected across all samples (Extended Data Table 1). Proteomic analysis of endosomes showed that proteins involved in 13 biological processes, 19 cellular components, and 16 molecular functions were significantly affected (Fig.1d). These proteins are involved in 10 functional pathways, including translation complex formation and initiation as well as metabolism of polyamines, amino acids and derivatives. Eighteen proteins were higher in the group that was allowed to sleep (SD plus recovery) than the sleep-deprived (SD) group (Fig.1b, c). Interestingly, this group included detectable level of Proteolipid protein 1 (PLP1), which are unlikely to originate from endothelial expression. PLP1 is a major constituent of central nervous system (CNS) myelin, produced almost exclusively by oligodendrocytes, and is not expressed in vascular endothelial cells^8,9^. Its detection therefore most likely reflects uptake of myelin-derived vesicles or debris from the brain parenchyma. Although we cannot exclude the possibility that it represents contamination of our samples, absence of other neuronal/glial proteins supports their active trafficking across the blood–brain barrier. Other proteins upregulated in the group allowed to sleep—Thumpd3, Dhx29, Ndor1, Tmed10, Prkra, Ube4b, Rpl22, Paip1, and Mtmr3— are highly expressed within brain endothelial cells. Meanwhile, 26 proteins were lower in expression in the SD plus recovery group than in the control group.

**Fig. 1:**
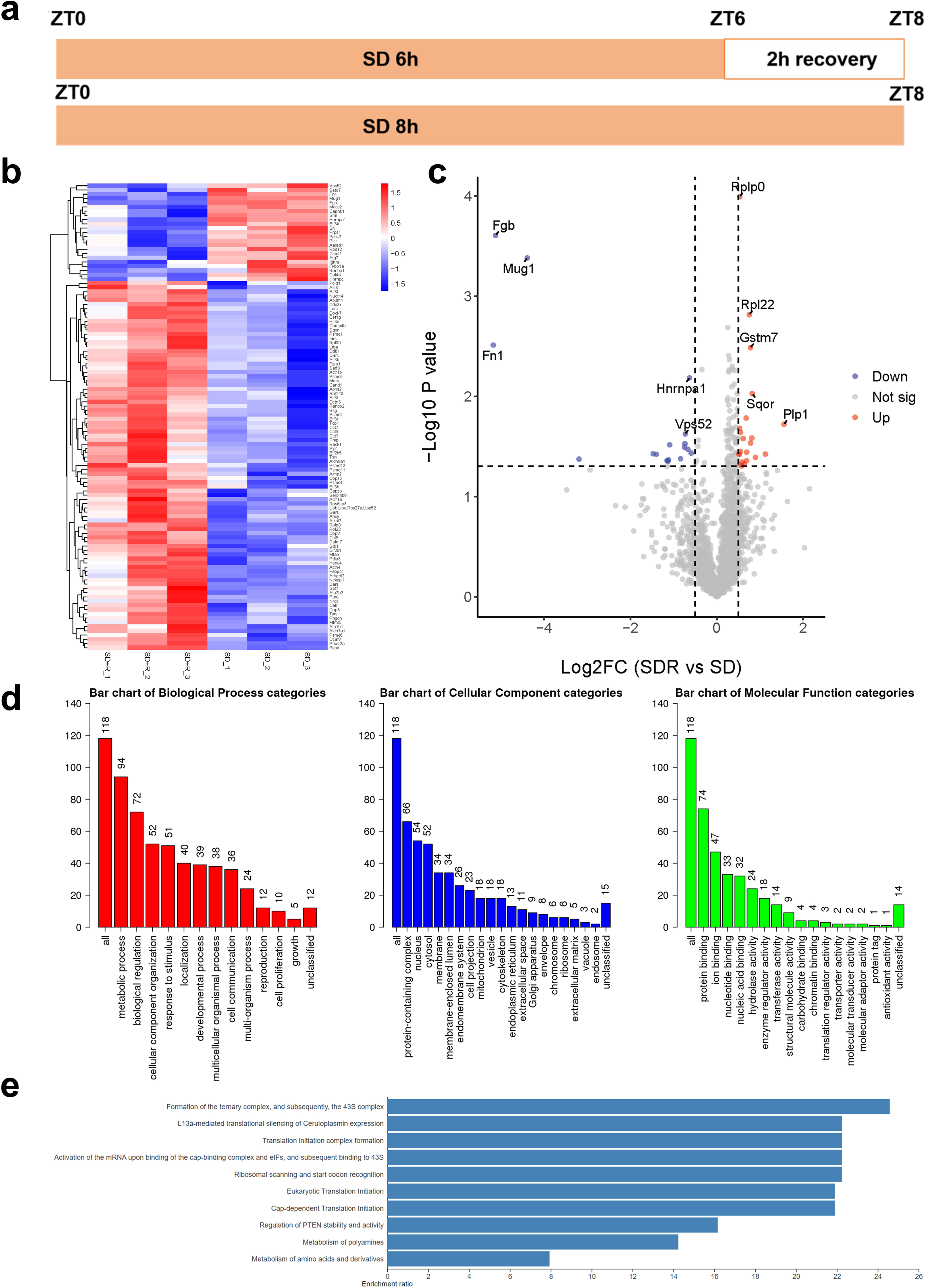
Proteomic analysis of endosomes isolated from brain endothelial cells following 6 hours of sleep deprivation with or without 2 hours of recovery. a. Sleep deprivation and recovery protocols for experimental and control groups of mice. b. Heat maps of relative protein levels, obtained via proteomics, in endosomes isolated from brain endothelial cells of sleep-deprived mice (SD 8h only) and mice allowed recovery sleep (SD followed 2 hours recovery). n=3 per group. c. Volcano plot showing proteins expressed differentially between the SD 8h and SD 6h and 2h recovery groups. Black dots represent the applied significance thresholds of a t-test p-value = 0.05. Details are shown in Table and the Supplementary Information. d. Bar graph of Gene Ontology (GO) analysis of the biological processes, cellular components and molecular functions of the proteins affected differentially in the SD with recovery group compared to SD only group. e. Reactome pathway analysis for proteins enriched in the SD plus 2h recovery group as compared to SD alone.

Given that PLP1 is not expressed in the BBB, we asked if it is trafficked to endothelial cells, perhaps to facilitate removal of excess protein. To investigate this, we overexpressed GFP-tagged PLP1 in the brain and assayed for its presence in brain endothelial cells (BECs). AAV-DJ-PLP1-GFP was administered via intracerebroventricular (ICV) injection into the lateral ventricles of C57BL/6 mice to induce PLP1 expression in CNS parenchymal cells. As control, AAV-DJ-GFP was injected at the same location under identical conditions. AAV-DJ is a capsid variant with strong CNS tropism, often achieving higher transduction efficiency in the brain and spinal cord than natural serotypes such as AAV9^10^. As PLP1 has a long half-life (∼100 days), we collected mouse brains three months later to determine whether it had been transported to BECs. Confocal imaging revealed PLP1-GFP signal in CD31⁺ BECs (Extended Data Fig.1b, c), suggesting transfer or uptake from surrounding parenchymal cells. Importantly, under identical conditions, AAV-DJ-GFP showed robust parenchymal expression but no detectable signal in BECs (Extended Data Fig.1a, c), indicating that PLP1 localization is cargo-dependent rather than a result of direct viral transduction. These findings support a model in which PLP1 produced in parenchymal cells can be selectively internalized by the BECs.

To determine if similar changes occur with sleep in a daily cycle, we collected brain endothelial cells from mice at two time points, ZT8 (sleep phase) and ZT20 (wake phase). Endosomes isolated from these cells were subjected to proteomic analysis using LC–MS/MS. In total, 2,133 proteins were detected across all samples (Extended Data Table 2). Among these proteins, 10 were specifically enriched in endosomes from mice in the sleep phase (Extended Data Fig. 3a, b). The most striking was the circadian and sleep-associated regulator, growth hormone (Gh1)^11^, which showed a 90-fold increase (Extended Data Fig. 3a, b). Another hormonally regulated molecule RACK1^12^ was highly expressed as well. Additionally, several proteins involved in endocytosis and vesicle trafficking, such as Arhgef18^13,14^, Lrrk2^15,16^, Gripap1^17^, and FKBP15, were upregulated between 2- and 8-fold. More interestingly, sleep phase BECs showed enrichment of metabolic and enzymatic components that regulate endosomal metabolism, membrane lipid remodeling, PI (3)P and autophagy-related endosomes, namely Hdhd3, Lypla1, Mtmr3 and Ears2 respectively. When comparing SD+R with SD and ZT8 with ZT20, we found that a few proteins-RACK1, Mtmr3, Eef1g, Gars, Mtap, Prep, Rplp0 –were consistently enriched under sleep conditions (SD+R and ZT8). Actbl2 protein was upregulated in the SD+R group and downregulated in the ZT8 group, suggesting that its regulation is not strictly sleep-dependent. Taken together, these findings suggest a sleep-specific endothelial program that enhances endocytosis and endosome maturation, raising the question of whether transport across the BBB is more active during sleep.

Waste products resulting from tissue activity are often secreted in the form of extracellular vesicles, or exosomes. To determine if this occurred with sleep deprivation, we examined vesicular changes in the mouse brain. Following 6 hours of sleep deprivation, brains were harvested, fixed, and subjected to immunofluorescence staining for the exosome marker CD63 and the brain endothelial cell marker CD31^18,19^. Quantification revealed a significant increase in CD63-positive exosomes, indicating robust accumulation of exosomal vesicles after sleep deprivation (Extended Data Fig. 3c). This accumulation suggests that sleep deprivation promotes the buildup of metabolic waste products or cellular debris, which are packaged and exported through exosomes.

To obtain direct evidence that endocytosis is enhanced during sleep, we used 10 kDa dextran as a tracer to monitor trafficking from the brain through brain endothelial cells (BECs). To avoid sleep-like effects induced by prolonged anesthesia, dextran was delivered into the brains of mice that had previously been cannulated via stereotaxic surgery. After a three-day recovery period following surgery, one set of mice underwent 6 hours of sleep deprivation (SD) followed by 2 hours of recovery sleep, whereas another set was subjected to 8 hours of continuous SD. Dextran–Alexa Fluor™ 488 was injected through the cannula into the second cortical layer (Fig. 2a). Blood and brain tissue were collected 3 h after injection. The fluorescence intensity of dextran in the blood of SD plus recovery mice was more than fivefold higher than in SD (Fig. 2e). Consistently, immunostaining of brain sections revealed prominent colocalization of Dextran-488 with BECs in SD plus recovery mice but not in SD mice (Fig. 2b, c, d).

**Fig. 2:**
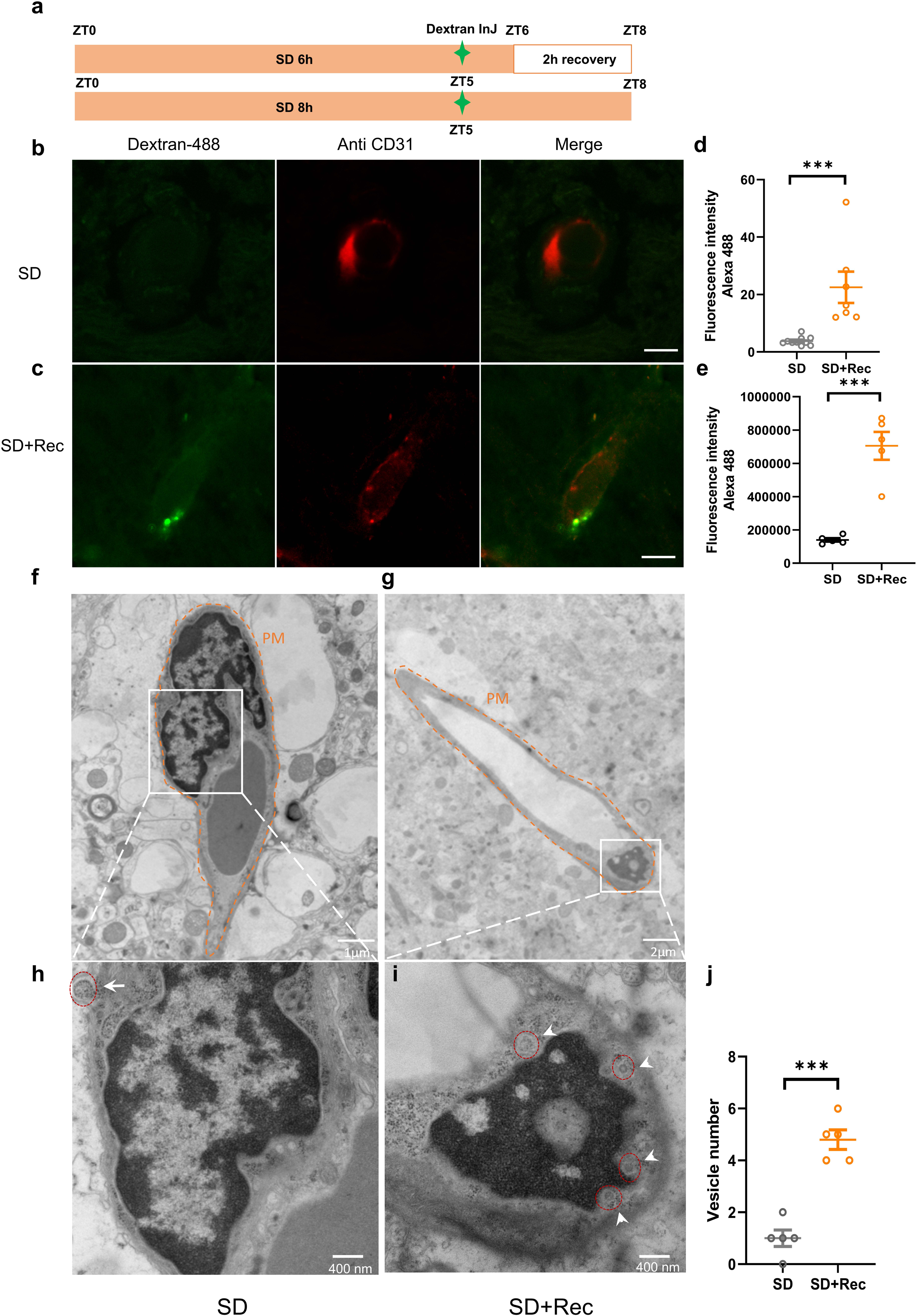
Vesicular transport is enhanced in the BBB during sleep. a. Sleep deprivation and recovery protocols for dextran injection and electron microscope imaging. b. Imaging of injected Dextran Alexa Fluor 488 in brain endothelial cells after 8 hours sleep deprivation only. c. Imaging of injected Dextran Alexa Fluor 488 in brain endothelial cells after 6 hours sleep deprivation and 2 hours recovery. d. Comparison of fluorescence intensity of brain endothelial cells between SD plus 2h recovery and SD only groups. n=10 cells for SD and n=7 cells for SD+Rec. e. The fluorescence intensity of dextran 488 was assayed in the blood at the same time (ZT8). n=5.f. Electron micrographs of brain endothelial cells after 8 h of sleep deprivation (SD). Direct magnification: 12,000×. g. Electron micrographs of brain endothelial cells after 6 h SD followed by 2 h recovery (Rec). Direct magnification: 6,000×. Each condition includes tissue from two mouse brains. h. Higher-magnification (40,000×) micrographs of the brain endothelial cells shown in panel f. i. Higher-magnification (40,000×) micrographs of the brain endothelial cells shown in panel g. j. Vesicles are outlined in red; arrowheads indicate individual vesicles. Quantification of vesicle number in panels f and g is shown (n = 6 per group). Scale bars in panels b and c represent 5 µm. SD, sleep deprivation; Rec, recovery; ZT, Zeitgeber time; PM, plasma membrane. \*\*\**P*<0.001; \*\**P*<0.01; \**P*<0.05.

To further confirm that brain endothelial cells (BECs) internalize dextran during sleep, we performed electron microscopy on mice subjected to the same treatment as in Fig. 1A. Electron micrographs revealed a marked increase in endocytic vesicles within BECs of experimental mice compared with controls (Fig. 2f–i). These vesicles frequently localized near the basolateral side, consistent with active uptake from the brain. Together, these findings provide ultrastructural evidence that endocytic activity in BECs is strongly enhanced during sleep.

Although these data supported sleep-dependent endocytosis through the BBB, we sought to exclude other mechanisms that could account for the sleep-dependent dextran transport. Thus, we blocked endocytosis by downregulating Dynamin 2 (DNM2), a GTPase essential for vesicle scission during endocytosis and exocytosis^20,21^. For this purpose, we used AAV-BR1-Dnm2_K44A (encoding a dominant-negative DNM2 mutant fused to GFP) where the BR1 serotype is a modified AAV2 variant with high specificity for the brain vascular endothelium^22,23^. The virus was delivered via tail vein injection, and then the mice were subjected to cortical cannulation followed by injection of Dextran-594. Three hours after injection, at a time of high sleep (ZT8), blood dextran levels were more than fivefold higher in control mice than in Dnm2_K44A mutants (Fig. 3d). Immunostaining further revealed that Dextran-594 colocalized with BECs in controls but not in Dnm2_K44A-expressing mice (Fig. 3a, b, c). Together these data indicate that blocking endothelial endocytosis suppresses both sleep-dependent dextran uptake by BECs and its clearance into the circulation.

**Figure 3:**
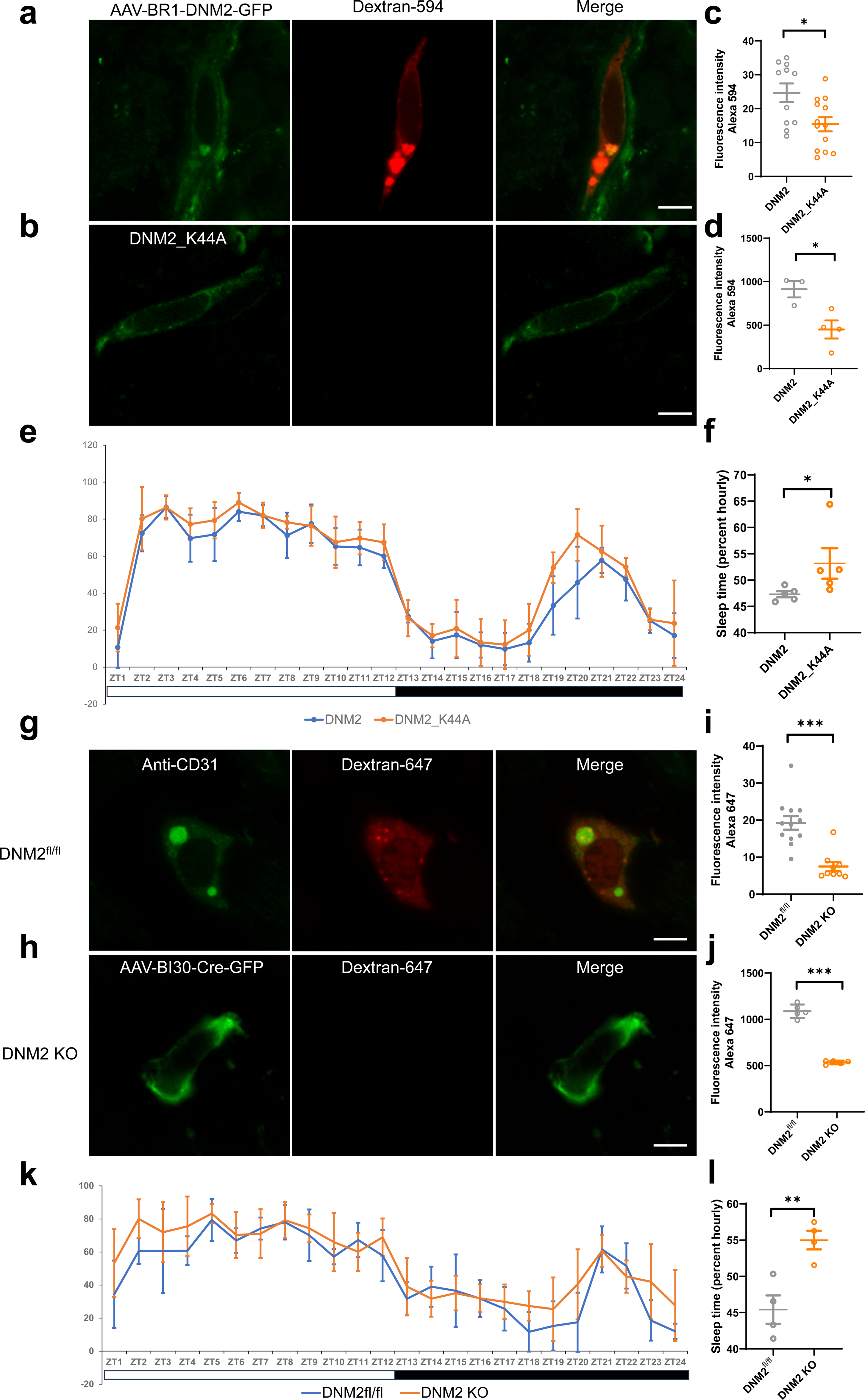
Mutant or knock out of Dynamin 2 in brain endothelial cells blocks endocytosis and increases sleep. a. Imaging of injected Dextran Alexa Fluor 594 in BECs of mice with wild type Dynamin 2 expressed by AAV-BR1-DNM2-GFP. b. Imaging of injected Dextran Alexa Fluor 594 in BECs expressing dominant negative dynamin with AAV-BR1-DNM2_K44A-GFP. c. Fluorescence intensity of dextran 594 in the BECs of Dynamin 2and Dynamin 2_K44A expressing mice. n=11 and 13. d. Fluorescence intensity of dextran 594 in the blood of Dynamin 2 and Dynamin 2_K44A expressing mice. n=3 and 4. e. Dynamin 2_K44A dominant negative mutant in the BECs increases sleep in mice. n=5. f. Quantification of sleep time in wide type and Dynamin 2_K44A mice. n=5 mice per group. h. Imaging of injected Dextran Alexa Fluor 647 in BECs of wild type mice and Dynamin 2 knockout mice. i. Imaging of injected Dextran Alexa Fluor 647 in the BBB of mice with Dynamin 2 knocked out. j. Fluorescence intensity of dextran 647 in the blood of Dynamin II and Dynamin 2 KO mice. Fluorescence intensity of dextran in brain endothelial cells of DNM2^flox/flox^ and dynamin 2 knock out mice. n=5. k. Knock out of Dynamin 2 in the BECs increases sleep in mice. n=4 in each group. l. Quantification of sleep time in wild type and Dynamin 2 knock out mice. n=4 in each group. fl, flox. ZT, Zeitgeber Time. DNM, Dynamin. Scale, 5 μm. \*\*\**P*<0.001; \*\**P*<0.01; \**P*<0.05.

We surmise that endocytosis from the brain through the BBB is promoted by sleep, and asked whether blocking it affects sleep amount. We allowed four weeks of expression of the AAV-BR1-Dnm2_K44A, and then assessed sleep using the Piezo sleep monitoring system. Mice expressing Dnm2_K44A displayed significantly increased total sleep compared with wild-type Dnm2 controls (Fig. 3e, f). This is similar to what is seen in Drosophila ^24^, and may indicate an increased need for sleep when a function of sleep is compromised.

To further validate the role of dynamin 2 in brain endothelial cells (BECs), we tried to knock out dynmin2 in BECs using a method described previously ^6^. However, we found that knockout of *Dnm2* in BECs is developmentally lethal in mice. We thereby chose an alternate strategy in which we injected AAV-BI30-Cre-GFP into Dnm2^flox/flox^ mice to selectively delete Dnm2 in the adult endothelium. AAV-BI30 is a capsid variant with high tropism for CNS endothelial cells and efficient penetration of the blood–brain barrier (BBB)^25^. After 4 weeks, the loss of Dnm2 expression in BECs following knockout was confirmed by immunostaining (Extended Data Fig. 5a, b). As before, we cannulated the mice and injected Dextran-647 at ZT5, during the sleep phase. Blood collected three hours later (ZT8) revealed markedly reduced fluorescence in Dnm2 knockout mice compared with controls (Fig. 3j). Immunostaining confirmed that Dextran-647 rarely entered BECs in knockout animals, whereas it was robustly internalized in controls (Fig. 3g, h, i). Conditional knockout of Dnm2 in BECs significantly increased total sleep time (Fig. 3k, l). These results demonstrate that loss of Dnm2 attenuates endothelial endocytosis at the BBB, perhaps impairing transport from the brain and increasing sleep need.

To determine whether endocytosis through the BBB has a diurnal rhythm, we first monitored sleep in C57BL/6 mice under LD12:12 conditions. As expected, mice predominantly slept around ZT2 and were largely awake at ZT14 (Fig. 4a). Accordingly, we injected Dextran-488 at ZT5 or ZT17 and collected blood and brains at ZT8 or ZT20. Fluorescence in the blood was dramatically higher at ZT8 than at ZT20 (Fig. 4e), and Dextran-488 colocalization with BECs was likewise greater during sleep (Fig. 4b, c, d). Further evidence for this came from comparison of endocytosis at ZT2 and ZT 14, with higher endocytosis observed at ZT2. Thus, endocytosis at the BBB is enhanced during daily sleep, facilitating the transport of brain-derived substances into BECs and their subsequent release into circulation.

**Fig. 4:**
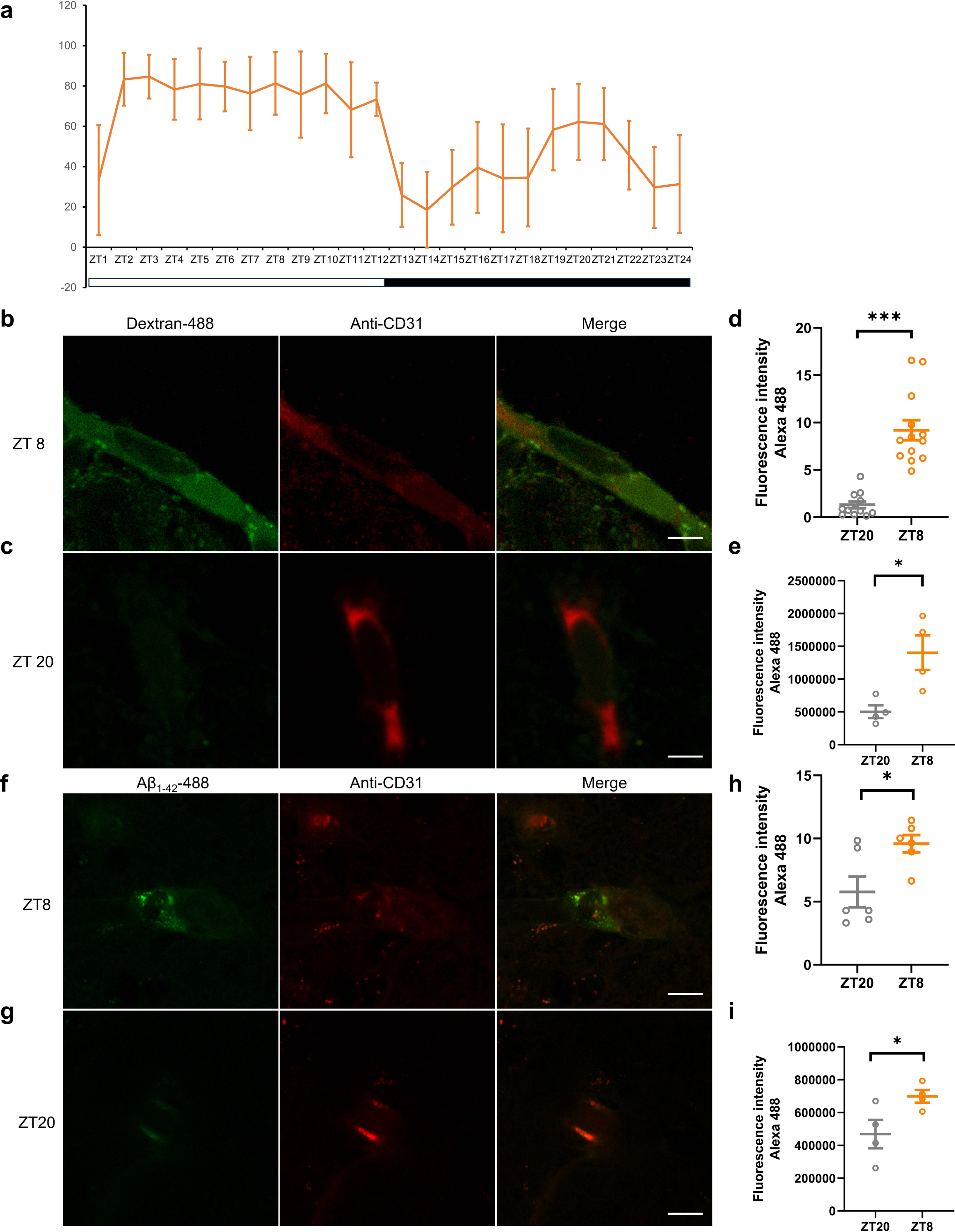
Transport of Aβ_1-42_ 488 through brain endothelial cells is enhanced during sleep. a. Sleep of C57BL/6J mice. b. Imaging of injected Dextran Alexa Fluor 488 at ZT8. c. Imaging of injected Dextran Alexa Fluor 488 at ZT20. d. Fluorescence intensity of Dextran in brain endothelial cells at ZT8 and ZT20. n=13 in ZT8 group and n=12 in ZT20 group. e. Fluorescence intensity of dextran 488 in the blood at ZT2 and ZT14. n=4 in ZT8 and ZT20 group. f. Imaging of injected Aβ_1-42_ 488 at ZT8. g. Imaging of injected Aβ_1-42_ 488 at ZT20. h. Fluorescence intensity of Aβ_1-42_ 488 in brain endothelial cells at ZT8 and ZT20. n=6 in each group. i. Fluorescence intensity of Aβ_1-42_ 488 in the blood at ZT8 and ZT20. n=4 in each group. ZT, Zeitgeber Time. Scale, 5 μm. \*\*\**P*<0.001; \*\**P*<0.01; \**P*<0.05.

Studies of clearance from the brain frequently examine the transport of Aβ_1-42,_ which contributes to amyloid plaque formation in Alzheimer’s disease (AD) ^26^. Thus, we tested whether sleep-dependent endocytosis through the brain also mediates the clearance of Aβ_1–42_. Aβ_1–42_-488 was injected into the cortex at ZT5 or ZT17, and blood and brains were harvested at ZT8 or ZT20, respectively. Consistent with the tracer results, Aβ_1–42_-488 levels were significantly higher in the blood at ZT8 than at ZT20 (Fig. 4i), and localization in BECs was greater during the sleep phase (Fig. 4f, g, h).

Although dynamin is considered a mediator of endocytosis, its loss also affects exocytosis by impeding vesicle trafficking. Our data suggest that endo and exocytosis are both important for sleep-regulated passage through the BBB. To specifically test for a role of exocytosis, we disrupted Exo70, a core component of the exocytotic machinery ^27,28^, in BECs using CRISPR-Cas9. AAV-BI30-saCas9-GFP and AAV-BI30-gRNA_Exo70_ were co-administered via retro-orbital injection. The knockout of Exo70 in BECs was confirmed by immunostaining for Exo70 (Extended Data Fig. 5c, d). Dextran-594 was infused at ZT5 (sleep phase) using cannulation. Blood collected three hours later (ZT8) showed markedly reduced fluorescence in Exo70 knockout mice compared with controls (Extended Data Fig. 3d). However, immunostaining demonstrated high accumulation of Dextran-594 within BECs of knockout mice, consistent with the idea that endocytosis is required for the uptake of dextran, and likely other material, from the brain but its release into the blood requires exocytosis (Extended Data Fig. 4a, b, c). As with the block in endocytosis, blocking exocytosis significantly increased sleep (Extended Data Fig. 4e, f).

Together, these findings demonstrate that endo and exocytosis through the BBB occur during sleep, likely to clear waste from the brain. We previously demonstrated that the activity of efflux transporters in the BBB is regulated by a circadian clock in endothelial cells and is timed to occur during wake ^6^. Sang *et al*. subsequently replicated our findings and showed too that sleep deprivation enhances efflux, indicating that efflux does not just occur during wake but is promoted by it ^7^. Interestingly, Aβ removal through the BBB is thought to involve both low-density lipoprotein receptor-related protein 1 (LRP1), which would act through an endocytic mechanism, as well as efflux transport ^29,30^. Given that we saw higher endocytosis during sleep, we speculate that Aβ is taken up by BECs via endocytosis during sleep and its extrusion from these cells can occur via exocytosis or efflux.

Our findings establish a direct, causal link between sleep, endothelial endocytosis, and transport of material, likely waste, from the brain. Unlike glymphatic flow, which relies on CSF-ISF exchange, this pathway operates at the vascular interface, allowing metabolites to be rapidly sequestered from the parenchyma and exported into systemic circulation. The rhythmic upregulation of endocytic machinery (e.g., Arhgef18, Lrrk2) and hormonal signals (Gh1) during sleep suggests an active, rhythmic program for vascular detoxification. This may explain why chronic sleep disruption is a risk factor for Alzheimer’s disease: impaired BEC endocytosis could promote Aβ and tau accumulation unless complementary enzymatic degradation mechanisms are engaged. PLP1-GFP expressed in CNS parenchymal cells was detected in CD31⁺ brain endothelial cells (BECs), whereas GFP alone showed no endothelial signal, indicating cargo-dependent transfer or uptake. This suggests that PLP1 can move from parenchymal cells to the BBB, likely for its clearance. The observation that blocking BEC endocytosis increases sleep reveals a homeostatic feedback loop: when clearance is inefficient, the brain compensates by demanding more sleep. This positions the BBB not merely as a static barrier, but as a dynamic sensor and effector of brain metabolic state. Future work should explore whether enhancing BEC endocytosis pharmacologically or by promoting sleep can reduce amyloid burden or rescue cognitive deficits in neurodegenerative models. If so, targeting the vascular clearance axis may offer a novel therapeutic strategy— one that works with, rather than against, the brain’s natural rhythms.

## Method

### Animal

Male C57BL/6J mice were obtained from Jackson laboratory (#000664, 013542, Jackson lab). All mice were housed in a temperature-controlled facility maintained at 22 ± 1°C with a 12-hour light/dark cycle (lights on at 07:00 and off 19:00 or lights on at 19:00 and off 07:00) with free access to food and water and were 10-12 weeks old at the time of surgery. All procedures were approved by Institutional Animal Care and Use Committees of the University of Pennsylvania and were done in accordance with the federal regulations and guidelines on animal experimentation (National Institutes of Health Offices of Laboratory Animal Welfare Policy).

The B6.129S1(Cg)-*Dnm2^tm^*^1^*^.1Pdc^*/J mice were obtained from Jackson laboratory (#013542, Jackson lab) and were born, bred and housed at University of Pennsylvania, Smilow Research Center of Translation animal facility. Breeding male and female B6.129S1(Cg)-*Dnm2^tm^*^1^*^.1Pdc^*/J mice and genotype offspring by PCR with the forward primer 5’-CCCTGCTAGTGACCTTTCTTGA-3’ and the reverse primer 5’-GCAGGAAGACACACAACTGAAC-3’. The Dnm2*^fl/fl^* mice were injected the AAV-CAG-Cre-GFP (AAV Serotype BI30) (SL116327, SignaGen Laboratories) to generate Dnm2 knock outs specifically in BECs. All animals included in experiments were also included in data analysis and results.

AAV-CAG-Dnm2-GFP (AAV-BR1) and AAV-CAG-Dnm2_K44A-GFP (AAV-BR1) were produced by the Vector Core of the Gene Therapy Program, Perelman School of Medicine at the University of Pennsylvania. The viral titers were 1.73×10¹² and 2.54×10¹² PFU/mL, respectively. Each virus was administered to 4 weeks old mice by retro-orbital injection using a total volume of 150 µL per animal. AAV-CAG-Exo70-GFP (AAV-BI30) and AAV-U6-gRNA-CBh-mCherry (AAV-BI30) were produced following the method described by Negrini et al ^31^. Viral titers, determined by qPCR, were 3.7×10¹² and 5.41×10¹² PFU/mL, respectively. Both vectors were delivered to 4-week-old mice via retro-orbital injection at a total volume of 200 µL per animal (100 µL of each virus).

An expression cassette encoding PLP1 alone was cloned into a pAAV-CAG-GFP transfer vector under the control of the CAG promoter. At the same time, pAAV-CAG-GFP empty vector was used the control vector. AAV-DJ particles were produced by co-transfecting the transfer plasmids with the pAAV-DJ capsid plasmid (#130878, Addgene) and helper plasmids in HEK293T cells. Viral titers, determined by qPCR, were 4.5×10¹² and 4.34×10¹² PFU/mL, respectively. For in vivo experiments, C57BL/6 mice (4 weeks old) received intracerebroventricular (ICV) injections into the lateral ventricles under stereotaxic guidance. Each group containing 5 mice was injected with either AAV-DJ-PLP1-GFP or AAV-DJ-GFP at the same site (bregma as the 0 point: AP = -0.2 mm, ML = ±1.0 mm, DV = -2.2 mm) and under identical conditions to serve as experimental and control groups, respectively.

### Isolation of brain endothelial cells (BECs) from mouse brain tissue

All steps were performed on ice unless otherwise specified. Initially, 6-well plates containing 6-8 mL of complete media per well (RPMI-1640 (Life Tech, 11875093) were supplemented with 10% Newborn Calf Serum (Sigma, N4673-500ML) and 1% Penicillin–Streptomycin (Life Tech, 15140-122)). Mice were euthanized with 3-5% Isoflurane, brains were rapidly dissected and immediately placed into ice-cold media. Using fine forceps, brain tissue was finely minced and transferred into a 50-mL conical tube. To the minced tissue, we added Collagenase/Dispase (100 mg/mL stock in H₂O) (Fisher, 17-104-019), 200 µL of DNase I Grade II (1 mg/mL stock in H₂O) (Sigma,10104159001), and 20 µL of Endosidin 2 (40 mM in DMSO) (Cayman, 21888). The total volume was brought to 20 mL with complete media. The suspension was incubated at 37°C with shaking at 250 rpm for 20 minutes to facilitate tissue dissociation. While the tissue was digesting, a fresh 22% Percoll solution (Fisher, 45-001-748) was prepared for myelin and debris removal. To do this, a Percoll bottle was vortexed thoroughly, then 11 mL of Percoll was added to a 50 mL conical tube, followed by 5 mL of 10×PBS and 34 mL of sterile water. Following thorough mixing, the solution was vacuum-filtered to remove particulates. Following the 20-minute digestion, the dissociated brain suspension was centrifuged at 500 × g (∼1250 rpm) for 5 minutes at 4°C. The supernatant was carefully aspirated and the pellet was resuspended in 1 mL of complete media (resulting in a total volume of approximately 2 mL including the cell pellet). For myelin and debris removal, 4 mL of the freshly prepared 22% Percoll solution was aliquoted into each of two 15-mL conical tubes per brain. 1 mL of the resuspended brain slurry was transferred into each tube and centrifuged at 1000 × g for 10 minutes at 4°C. After centrifugation, a compact cell pellet forms at the bottom of the tube, while myelin and debris accumulate at the interface or top. The supernatant was carefully poured off and a pipette was used to remove any residual myelin or debris without disturbing the cell pellet. If the cell suspension was clumpy, it was passed through a 100-µm cell strainer. Samples were pooled as needed (e.g., from multiple brains), then 5 mL of complete media was added and centrifuged again to pellet the cells. For positive selection of CD31⁺ endothelial cells, the final cell pellet was resuspended in 90 µL of PEB buffer (AutoRinse Buffer supplemented with BSA at a 1:20 dilution, stored at 4°C). 10 µL of CD31 MicroBeads (Miltenyi Biotec, 130-097-418) were added, mixed gently, and incubated on ice for 15 minutes. The MiniMACS Separator (Miltenyi Biotec, 130-042-102) was set up by placing an MS Column in the magnetic field. The column was pre-wet by rinsing it with 1 mL of PEB buffer. The labeled cell suspension was applied onto the column. The flow-through, which contains unlabeled (CD31⁻) cells, was collected. The column was washed with an additional 1 mL of PEB buffer and this effluent was collected as part of the CD31⁻ fraction. To recover the positively selected CD31⁺ cells, the column was removed from the magnetic separator and placed over a new collection tube. Immediately the column was flushed by applying 1 mL of PEB buffer and firmly pushing the plunger into the column to elute the magnetically retained CD31⁺ cells. This fraction represents the CD45⁻/CD31⁺ brain endothelial cell population. Finally, the isolated cells were counted. Typically, one sample consisting of pooled brains from five mice yields approximately 2 × 10⁶ CD31⁺ endothelial cells for proteomic analysis.

### Endosome collection from the brain endothelial cells

For endosomal proteins, brain endothelial cells were washed with cold PBS. The pellet was resuspended in 500 μl of Buffer solution of Minute Endosome Isolation (Invent biotechnologies, ED-028). The endosomal protein extraction was performed in accordance with manufacturer instructions. The endosomes were collected and sent to the Children’s Hospital of Philadelphia (CHOP) proteomics core for proteomic analysis.

### Stereotaxic surgery and dextran injection

To implant cannulae for dextran injection, mice were anesthetized with 3-4% isoflurane and placed on a stereotaxic frame. The guide cannula (INC22-1.5, World Precision Instruments) was inserted into the skull 1.2 mm from midline and 0.04 mm posterior to the bregma. The guide cannula was sticked to the skull with super glue and capped with a dummy cannula. After 3 days of the cannulation, 1 μl of 10000 Da dextran conjected with the Alexa Fluor 488 (D22910, Invitrogen) or the Alexa Fluor 594 (D22913, Invitrogen) or the Alexa Fluor 647 (D22914, Invitrogen) was injected into the mouse brain via a cannula by inserting into the internal cannula (INC22-1.5, World Precision Instruments).

### Sleep recordings

Sleep of C57BL/6J, Dnm2 knockout, and Dnm2_K44A mutant mice was recorded in home cages to which animals had been habituated, under a 12-h light/dark cycle (lights on at 07:00, off at 19:00). Sleep–wake states were monitored using a Piezo Adapt-A-Base sensor (AAB-Sense; Signal Solutions) connected to a 16-channel data acquisition system (16ch-DAQ; Signal Solutions). Sleep–wake classification was performed automatically using PiezoSleep 2.8 software and analyzed with SleepStats 2.18 (Signal Solutions).

### Blood fluorescent dextran measurement

Blood was collected via cardiac puncture into 1-mL lithium heparin–coated tubes (#450477, Greiner Bio-One). Samples were centrifuged at 2,000 × g for 20 min at 4 °C to obtain plasma. A 100-µL aliquot of each plasma sample was transferred to a 96-well plate, and fluorescence was measured at 488 or 647 nm by using a microplate reader (SpectraMax iD5, Molecular Devices).

### Proteomics of endosomes

Endosomal protein extracts were denatured, reduced, alkylated, and digested into peptides. The peptides were analyzed in single shot (per sample) on QE-HF in DIA mode by LC–MS/MS. Peptides were identified by using Spectronaut software (DIA, Biognosys) to compare with mouse reference proteome (25,125 entries, Reviewed canonical and isoforms). Perseus was used for statistical data analysis (peptides and proteins filtered at 1% FDR; minimum 2 razor+unique peptides for protein quantification). Contaminants, reverse hits and proteins identified only by site were removed. LFQ intensities were log₂ transformed. Differential abundance was assessed using a two-sample moderated t-test implemented in limma; p-values were adjusted by Benjamini–Hochberg (FDR). Proteins with |log₂ fold change| ≥ 1 and FDR < 0.05 were considered significant. Heatmaps were generated from row-scaled log₂ LFQ intensities (z-score per protein) and clustered by Euclidean distance with complete linkage. Volcano plots show log₂ fold change versus −log₁₀(FDR) with significant proteins highlighted. Functional enrichment on significant hits was performed using STRING or DAVID. All analyses and plotting were performed in R.

### Electron-microscopy

Following either 6 h of sleep deprivation plus 2 h of recovery or 8 h of continuous sleep deprivation, mice were deeply anesthetized and were fixed with 2.5% glutaraldehyde, 2.0% paraformaldehyde in 0.1M sodium cacodylate buffer, pH7.4, overnight at 4℃. After subsequent buffer washes, the samples were post-fixed in 2.0% osmium tetroxide for 1 hour at room temperature, and rinsed in dH2O prior to *en bloc* staining with 2% uranyl acetate. After dehydration through a graded ethanol series, the tissue was infiltrated and embedded in EMbed-812 (Electron Microscopy Sciences, Fort Washington, PA). Thin sections were stained with uranyl acetate and lead citrate and examined with a JEOL 1010 electron microscope fitted with a Hamamatsu digital camera and AMT Advantage NanoSprint500 software. Embedding, sectioning, and staining were performed by the Electron Microscopy Resource Laboratory at the University of Pennsylvania.

Representative images were obtained from a single brain per group. All image analyses were performed blinded to experimental conditions. For vesicle quantification, vesicles or subcellular structures were counted per cell using Fiji/ImageJ. Statistical analyses were conducted on biological replicates (n values are indicated in figure legends) using unpaired t-tests as described in the main text.

### Plasmid cloning

For the AAV-CAG-Dnm2-GFP and AAV-CAG-Dnm2_K44A-GFP plasmids construction, the coding region of the Dnm2 and Dnm2_K44A genes were amplified by PCR using RFP Dynamin2 Wild Type (#128152; Addgene) and RFP Dynamin2 K44A (#128153; Addgene) plasmids as templates and the forward primer ‘AAATCTAGAATGGGCAACCGCGGGATGGAAG’ and the reverse primer ‘AAAGGATCCGTCGAGCAGGGACGGCTCGGCT’. The PCR products were gel purified, digested with *Xba*I and BamHI, and ligated into the pAAV-CAG-GFP (#37825; Addgene) vector precut with the same enzymes. The AAV-CAG-saCas9-GFP plasmid construction, the coding region of SV40 NLS, saCas9 and GFP genes were amplified by PCR using pSaCas9_GFP (#64709; Addgene) as template and the forward primer ‘AAAGGATCCATGGCCCCGAAGAAAAA’ the reverse primer ‘AAAAAGCTTTTACTTGTACAGCTCGT’. The PCR products were gel purified, digested with BamHI and HindIII, and ligated into the pAAV-CAG-GFP (#37825; Addgene) vector precut with the same enzymes. The guide RNA vector of Exo70 were generated by annealing the oligos ‘CACCCCTTG CTG TGC CGG GTCATC’ and ‘AAACGATGACCCGGCACAGCAAGG’ and insert pSpCas9(BB)-2A-GFP (PX458) (#48138; Addgene) with the precut BbsI (BpiI) sites. After the KO effect was confirmed in the Neuro-2a cells, the guide RNA vector of Exo70- pSpCas9(BB)-Exo70-2A-GFP was subcloned into the pAAV-U6-gRNA-CBh-mCherry (#91947; Addgene) with the NdeI sites. The cloning of mouse PLP1 gene by using the cDNA from Neuro-2a cell line with the forward primer ‘ATG GGC TTGTTAGAGTGTTGT’ and the reverse primer ‘TCAGAA CTTGGTGCCTCGGCCCA’. The coding region of PLP1 gene was inserted the plasmid pAAV-CAG-GFP (#37825; Addgene) vector.

### Histology for endocytosis

Three hours after dextran injection, mice were deeply anesthetized and subjected to cardiac blood sampling. For histology of BECs, brains were fixed overnight in 4% paraformaldehyde (PFA) in PBS and then transferred to 30% sucrose in PBS solution for at least two nights. Brains were embedded and mounted with Tissue-Tek OCT compound (Sakura Finetek) and frozen. 20 μm sections were cut using a cryostat (CM3050, Leica) and mounted onto glass slides. Brain sections were washed in PBS, permeabilized using PBST (0.3% Triton X-100 in PBS) for 30 min and then incubated in blocking solution (3% normal calf serum (NCS) in PBST) for 1 h. Slices were then incubated in primary antibodies (1:200, anti-CD31 antibody, MA1-26196, Invitrogen) overnight at 4 °C. 18 h later, slices were washed with PBS and incubated with the corresponding secondary antibody (1:1000, Alexa Fluor 594 - Highly Cross-Adsorbed Goat anti-Mouse IgG (H+L), A32742 or Alexa Fluor 647 - Highly Cross-Adsorbed Goat anti-Mouse IgG (H+L) A-21236 Invitrogen). Brain sections were washed with PBS and mounted onto glass slides with mounting medium (#H1200, Vector Labs). Slides were cover-slipped with micro cover glass (#48404-455, VWR).

### Histology for knockout of Dynamin2 and Exo70 in BECs

Four weeks after injection of AAV-BI30-Cre-GFP or AAV-BI30-saCas9 together with AAV-BI30-gRNAEx070-CBh-mCherry, experimental and control mice were deeply anesthetized, and brains were harvested. Brains were fixed overnight in 4% paraformaldehyde (PFA) in PBS, then transferred to 30% sucrose in PBS for at least two nights. Fixed brains were embedded in Tissue-Tek OCT compound (Sakura Finetek) and frozen. Coronal sections (20 μm) were cut using a cryostat (CM3050, Leica) and mounted onto glass slides. Sections were washed in PBS, permeabilized with PBST (0.3% Triton X-100 in PBS) for 30 min, and blocked in 3% normal calf serum (NCS) in PBST for 1 h. Slices were then incubated overnight at 4 °C with primary antibodies: anti-Dynamin2 (1:50, PA5-29658, Invitrogen) or anti-Exo70 (1:100, 12014-1-AP, Proteintech). After 18 h, sections were washed with PBS and incubated with secondary antibody (1:1000, Alexa Fluor 594 Highly Cross-Adsorbed Goat anti-Rabbit IgG (H+L), A32740, Invitrogen). Finally, sections were washed with PBS, mounted with medium (#H1200, Vector Labs), and cover-slipped with micro cover glass (#48404-455, VWR). Loss of Dynamin2 or Exo70 immunoreactivity in GFP brain endothelial cells was used to confirm efficient gene knockout.

### Confocal microscopy of brain endothelial cells

All fluorescent images of cells were acquired with an inverted Leica SP8 confocal microscope equipped with a motorized and piezo z stage, an HC PL APO CS2 40×/1.30 oil-immersion objective (Leica Microsystems, Mannheim, Germany), Laser Diodes 405, 488, 594, and 647, and HyD detectors. The settings for each image were Pinhole 1 Airy unit, 16-bit acquisition, 4x zoom, 2048x2048 pixels field of view, 400 Hz scan speed, resulting in a pixel size of 0.07 μm. All channels were detected on HyD detectors in multiple individual series to reduce bleed through: (1) Dextran (Alexa 488) and BECs (Alexa 594) channels (2) Dextran (Alexa 647) and BECs (Alexa 594) channels (3) AAV-BI30-Cre-GFP (Alexa 488) and Dextran (Alexa 594) channels (4) AAV-BI30-saCas9-GFP and Dextran (Alexa 594) channels (5) AAV-BI30-Cre-GFP (Alexa 488) and Dynmin2 (Alex 647) channels (6) AAV-BI30-saCas9-GFP and Exo70 (Alexa 594) channels detected. The detector gains and the laser intensity for the dextran (Alexa 488 or Alexa 647) channel were constant for all images acquired. Detector gains and laser intensities for the other channels were minimally adjusted between different plates and cell lines, if needed.

### Image analysis

Confocal micrographs of dextran and the BECs were selected randomly. After the background subtraction, using the BG Subtraction from ROI plugin of ImageJ software (version 1.44; National Institutes of Health), signal was quantified with the Intensity Analysis plug-in of ImageJ software.

### Quantification and statistical analysis

Sleep comparisons between control and experimental groups (Fig. 3 and Extended Data Fig. 4) were evaluated using Welch’s *t*-tests, unless otherwise specified in the figure legends. Comparisons of dextran fluorescence intensity in blood or brain endothelial cells (Figs. 2–4) were analyzed using Mann–Whitney *t-*tests, as were comparisons of exosome numbers (Extended Data Fig. 1) and endosomal vesicle counts (Fig. 2h). All statistical analyses were performed in GraphPad Prism. Detailed information on statistical tests, sample sizes, and significance thresholds is provided in the figure legends.

## Acknowledgements

We thank the Neurobehavior Testing Core at UPenn/ITMAT and IDDRC at CHOP/Penn P50 HD105354 for help with cannulations.

## Author contributions

Conceptualization: F. L. and A.S.; Methodology: F. L. and A.S.; Investigation: F. L. and D. C.; Writing: F.L. and A.S.

## Extended Data Figure legends

**Extended Data Table 1.**
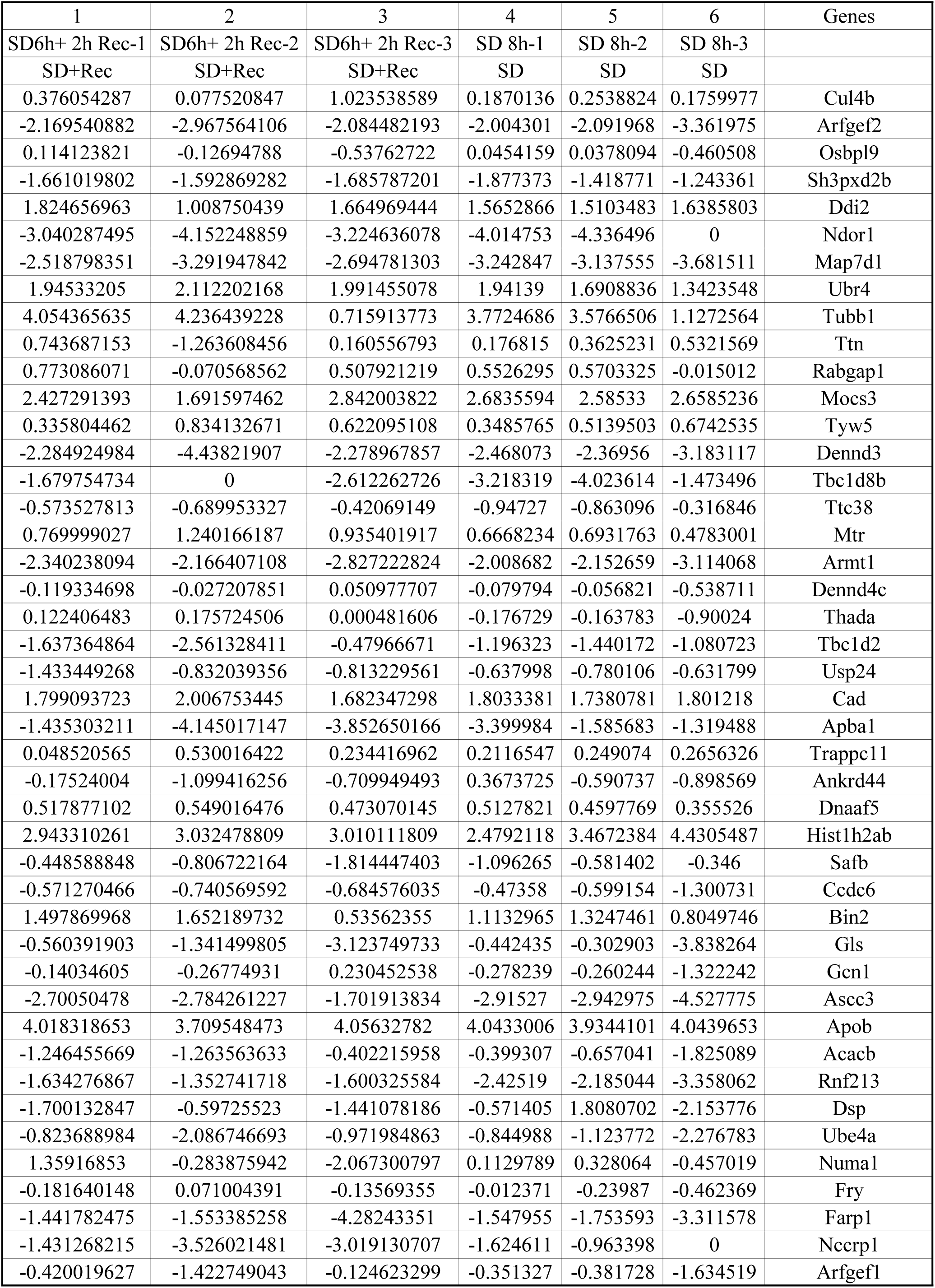
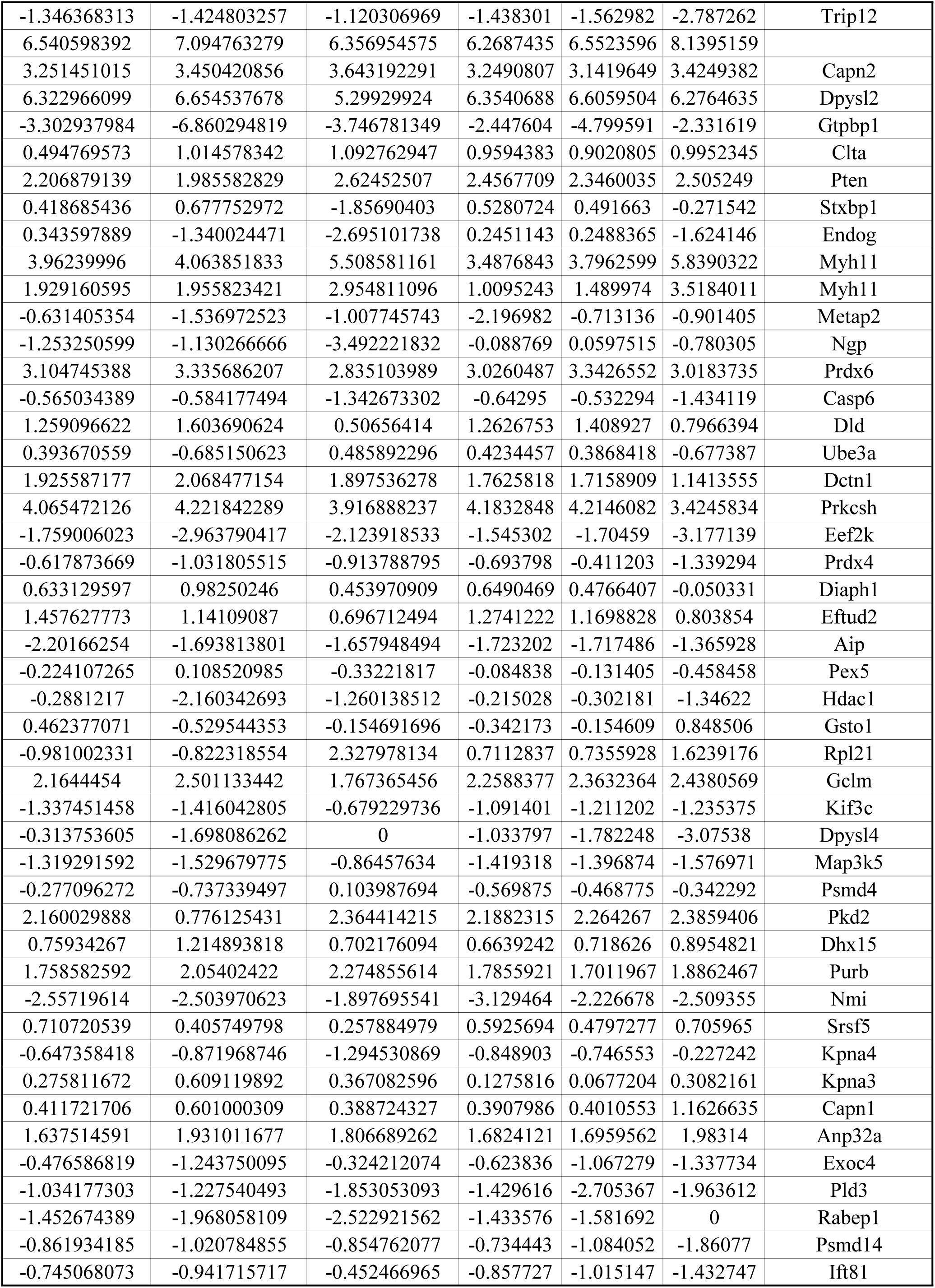
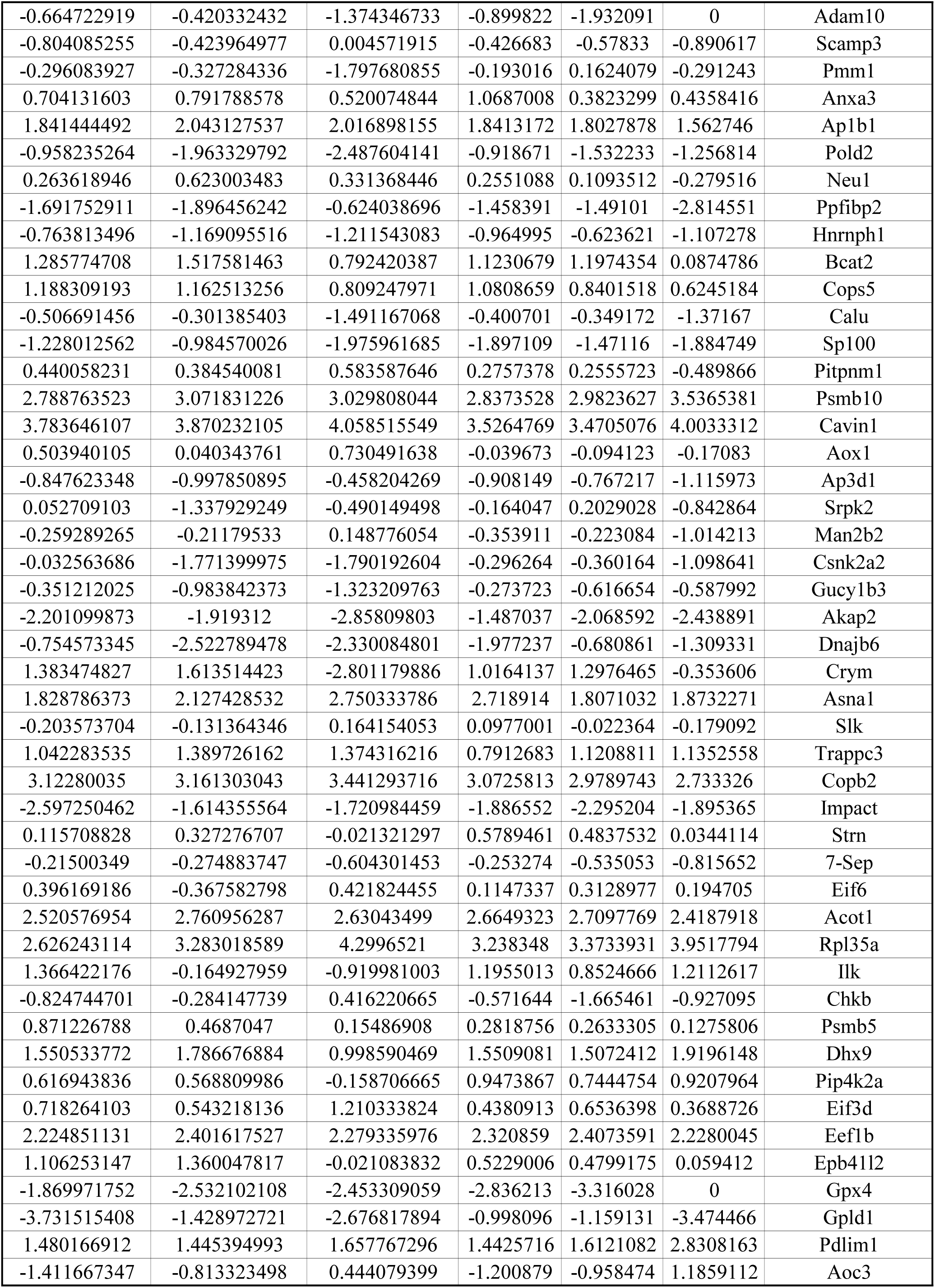
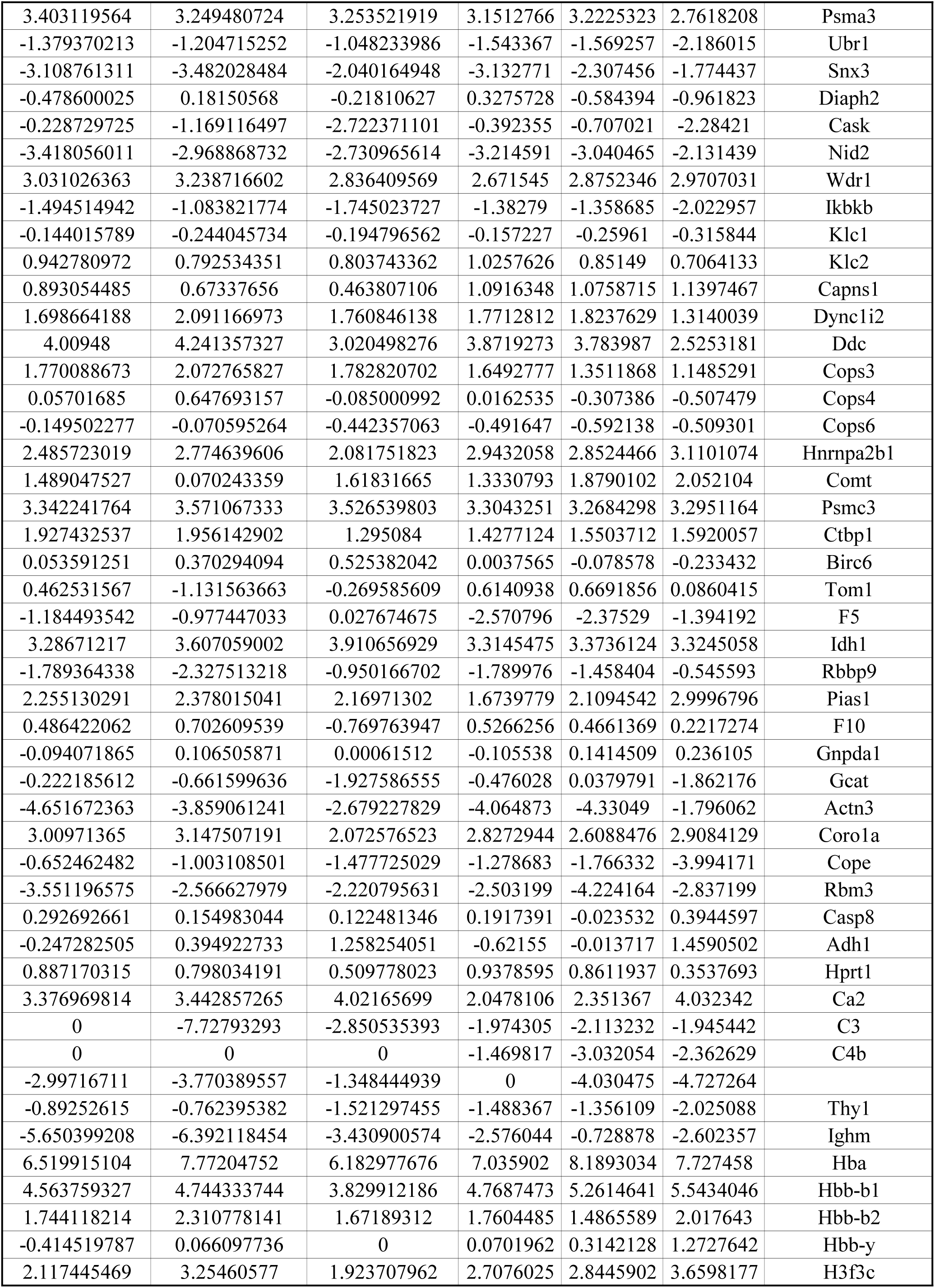
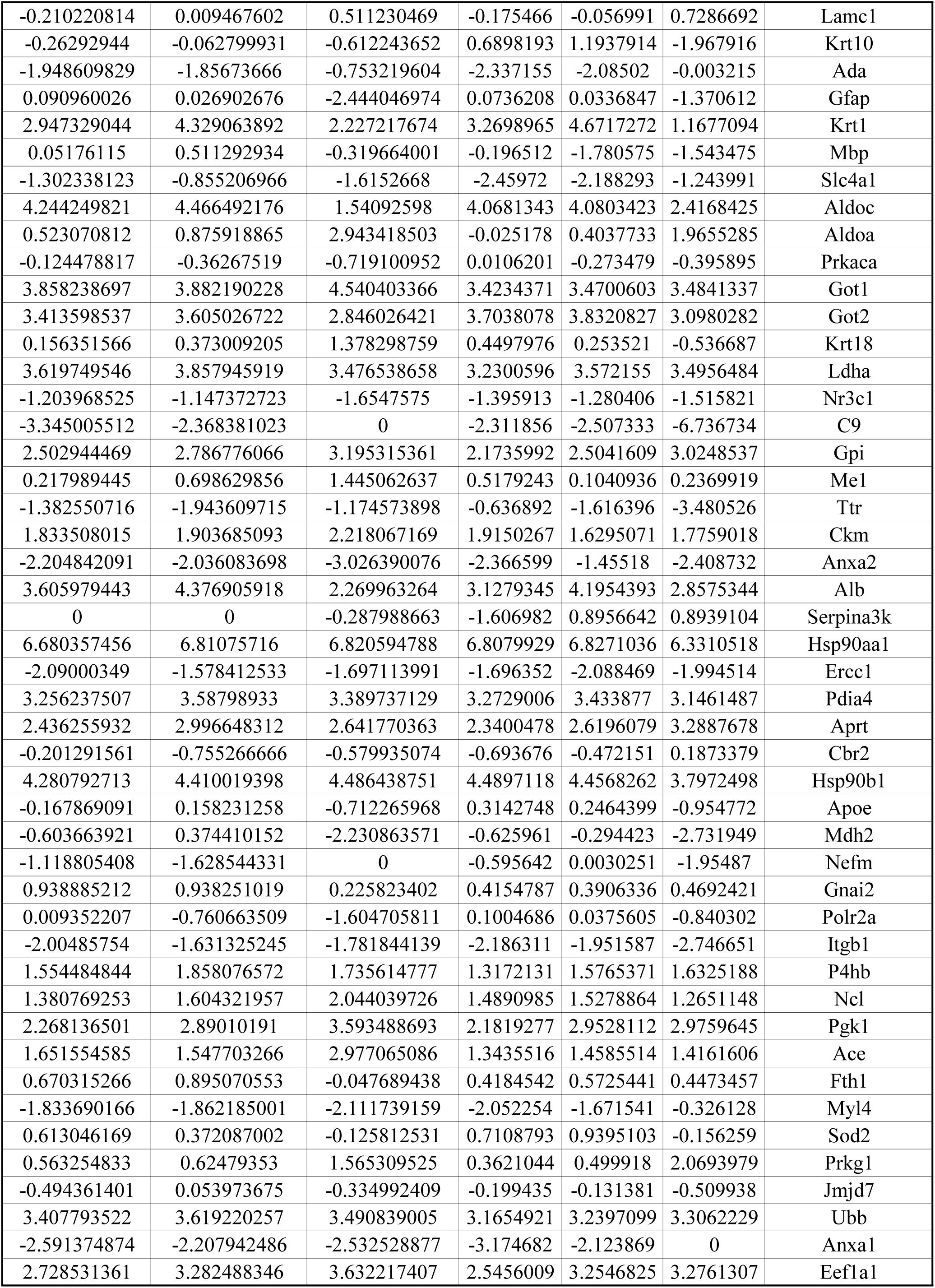
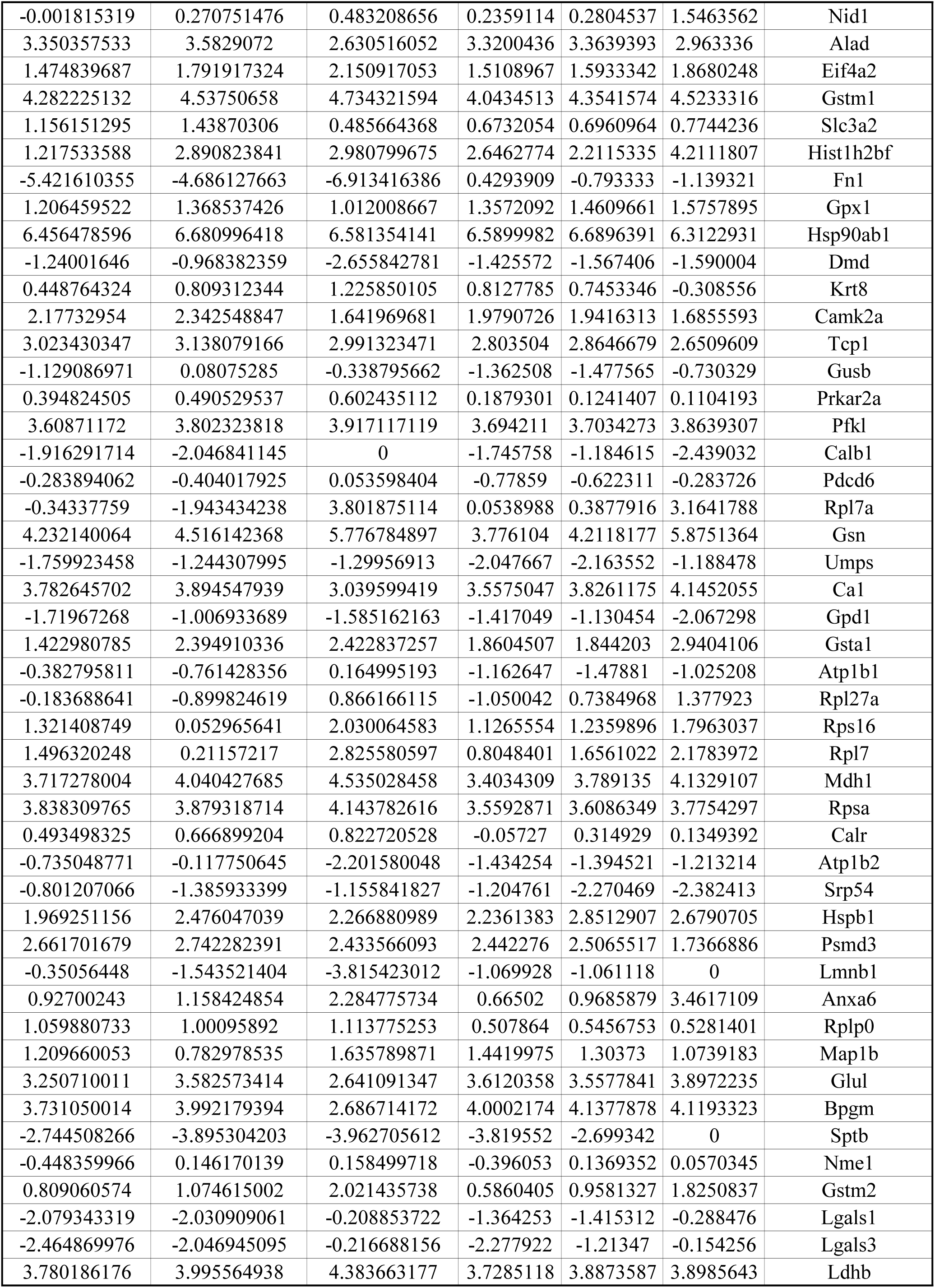
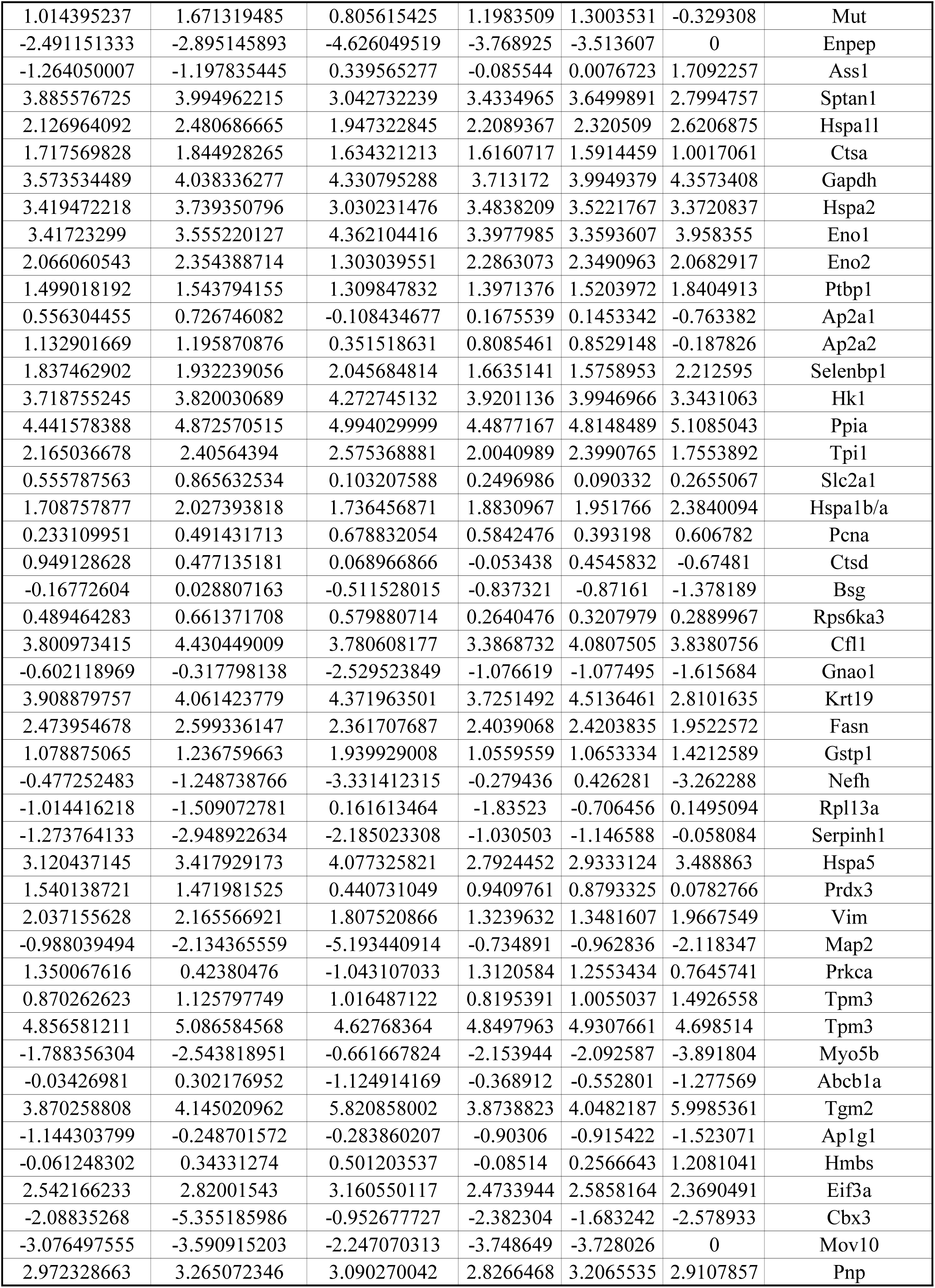
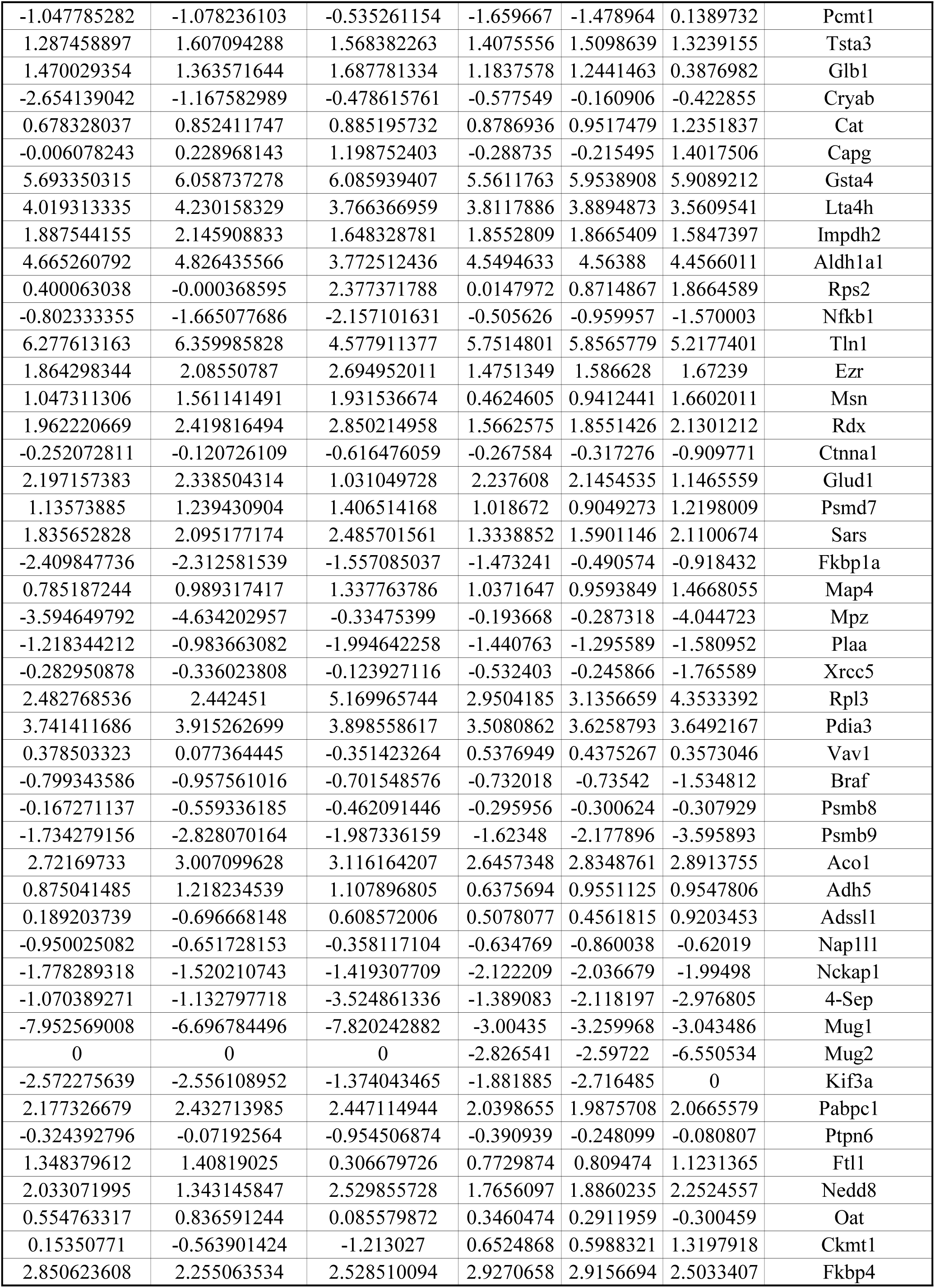
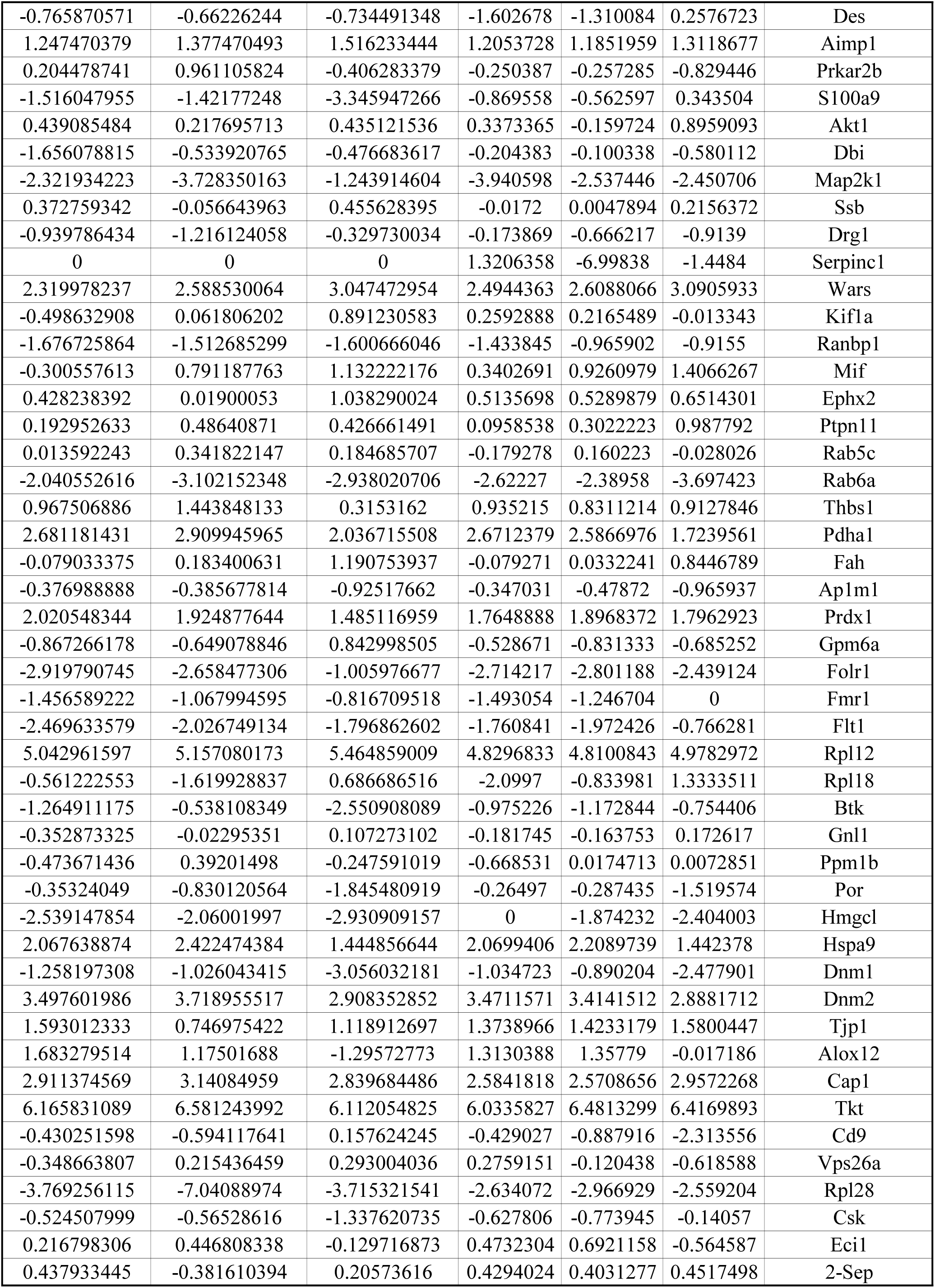
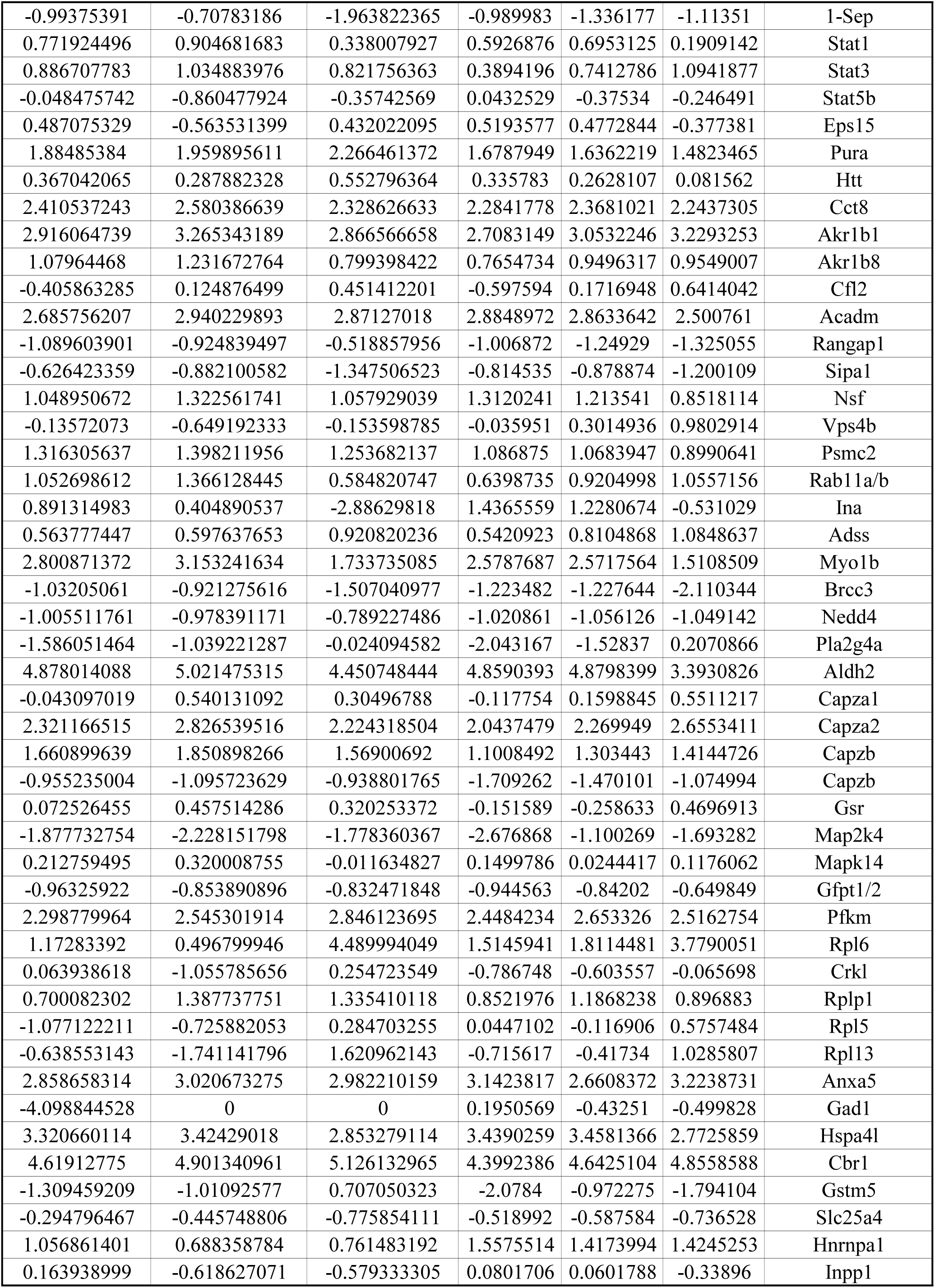
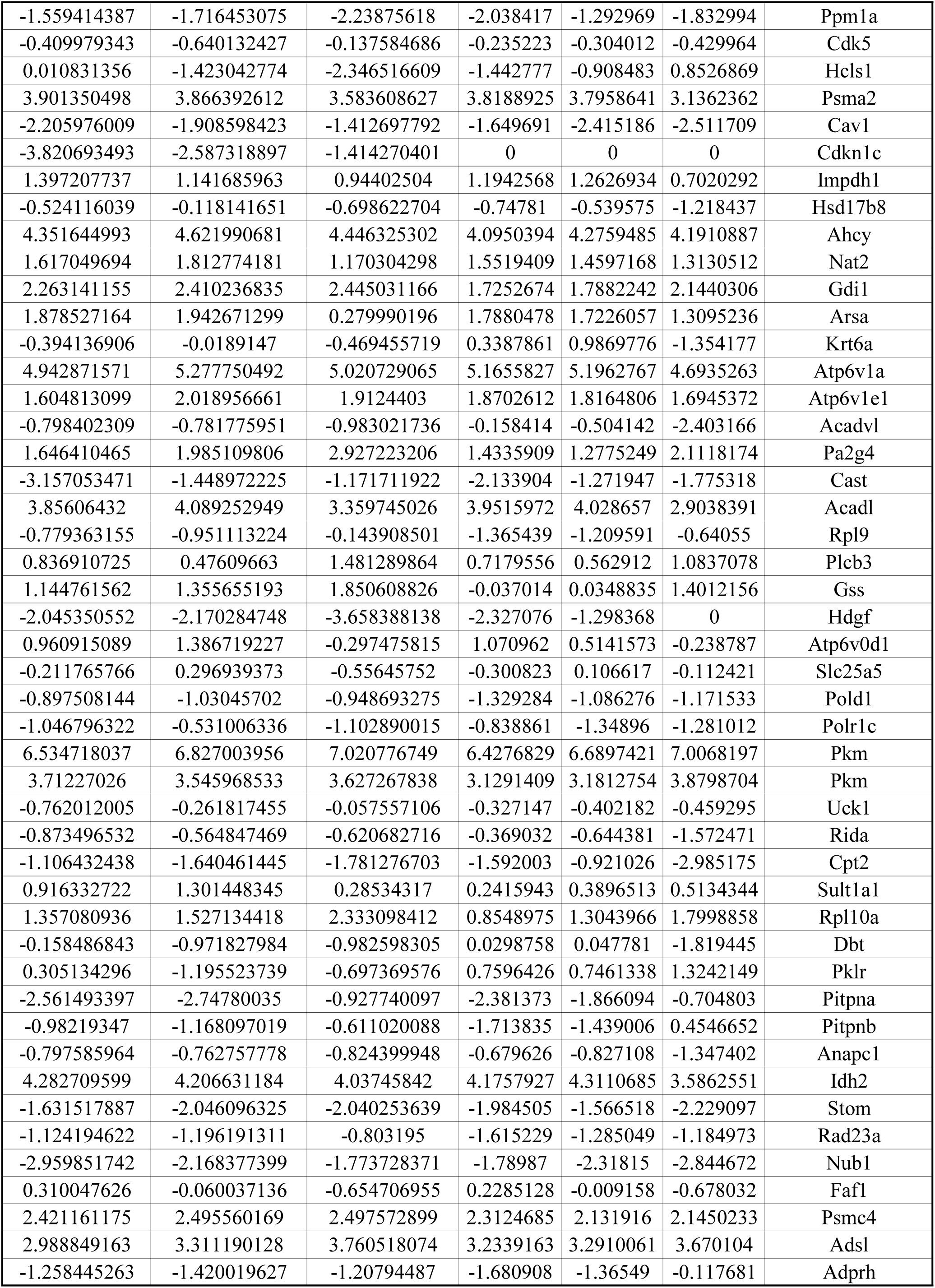
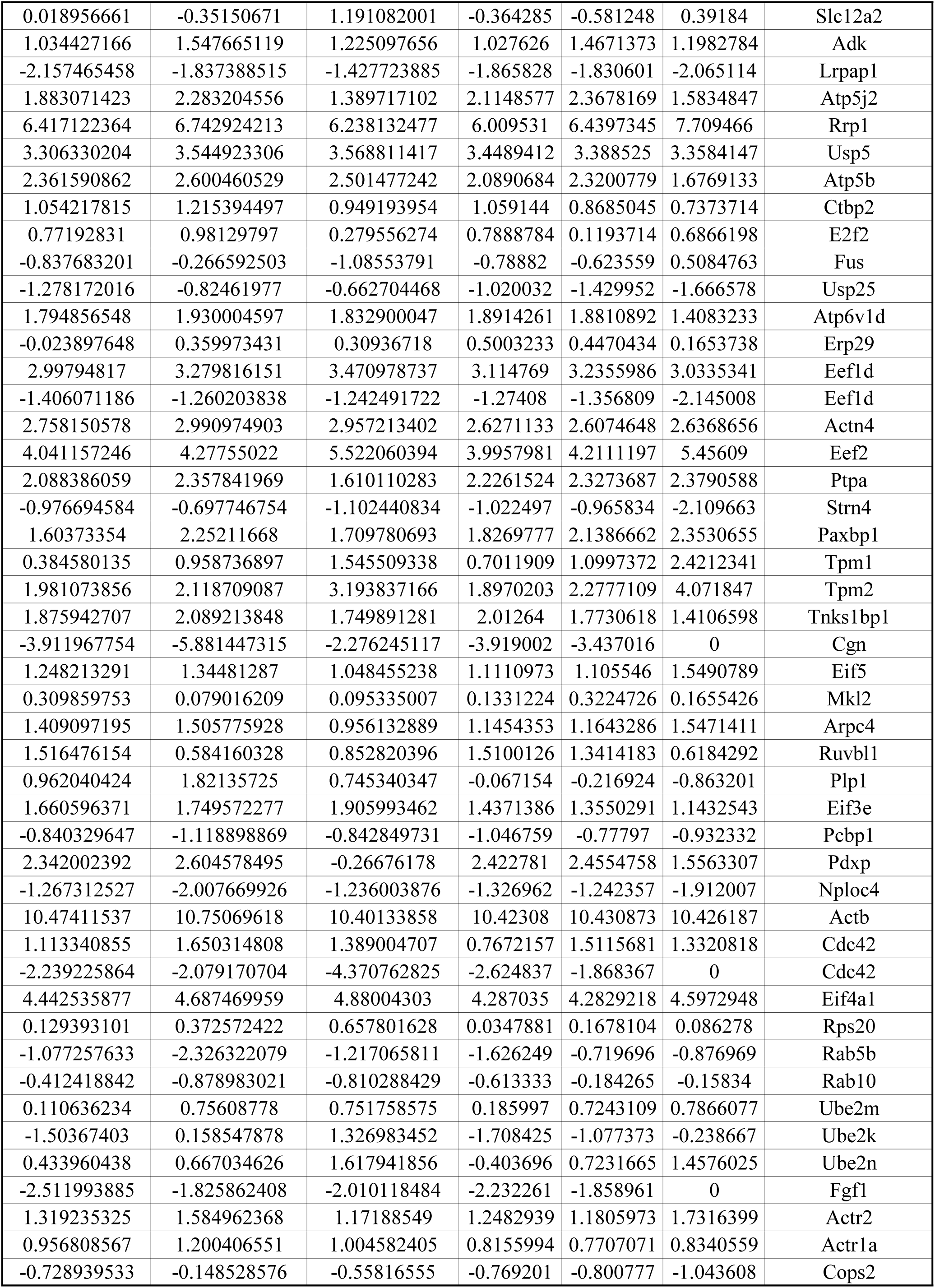
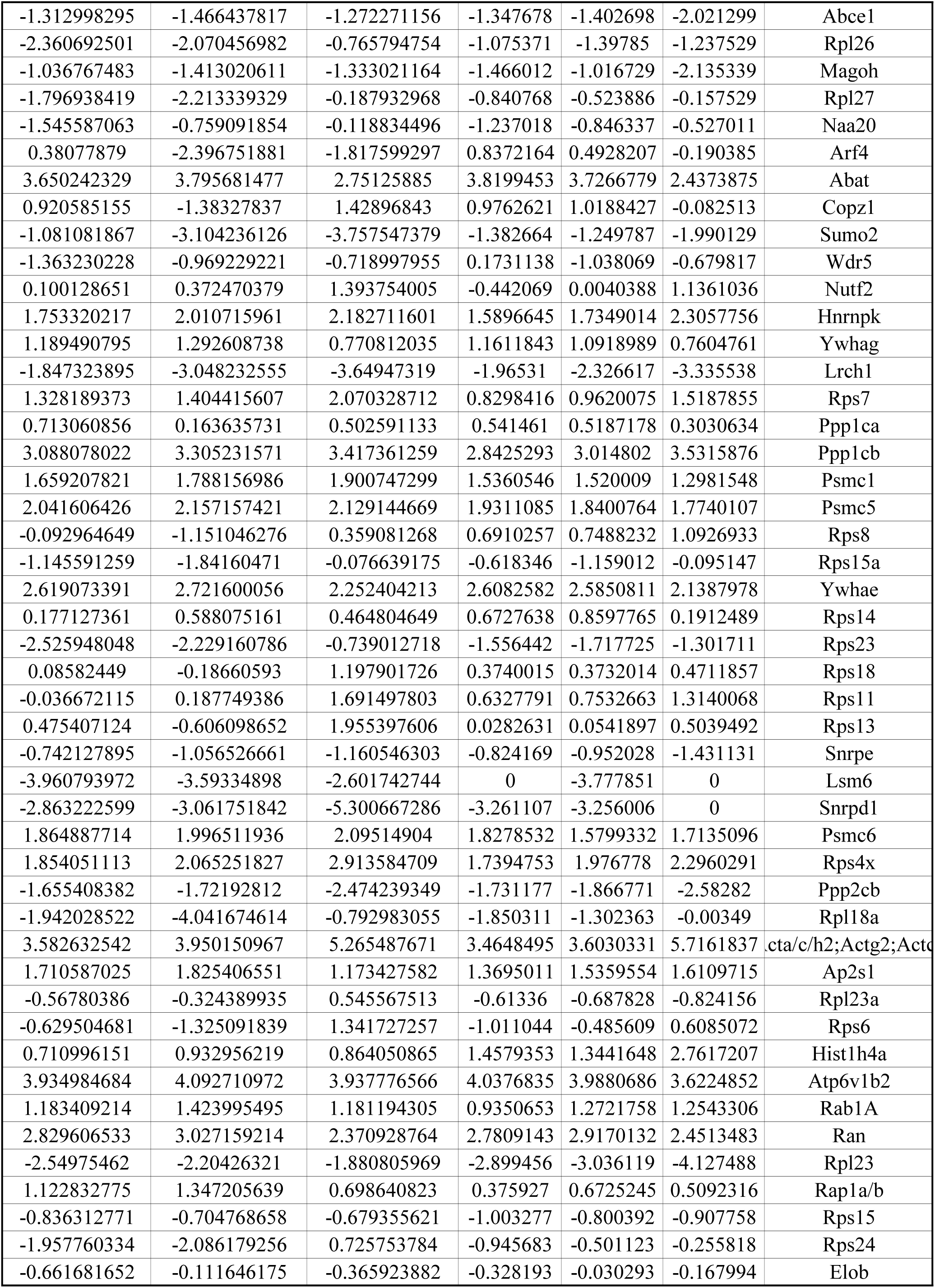
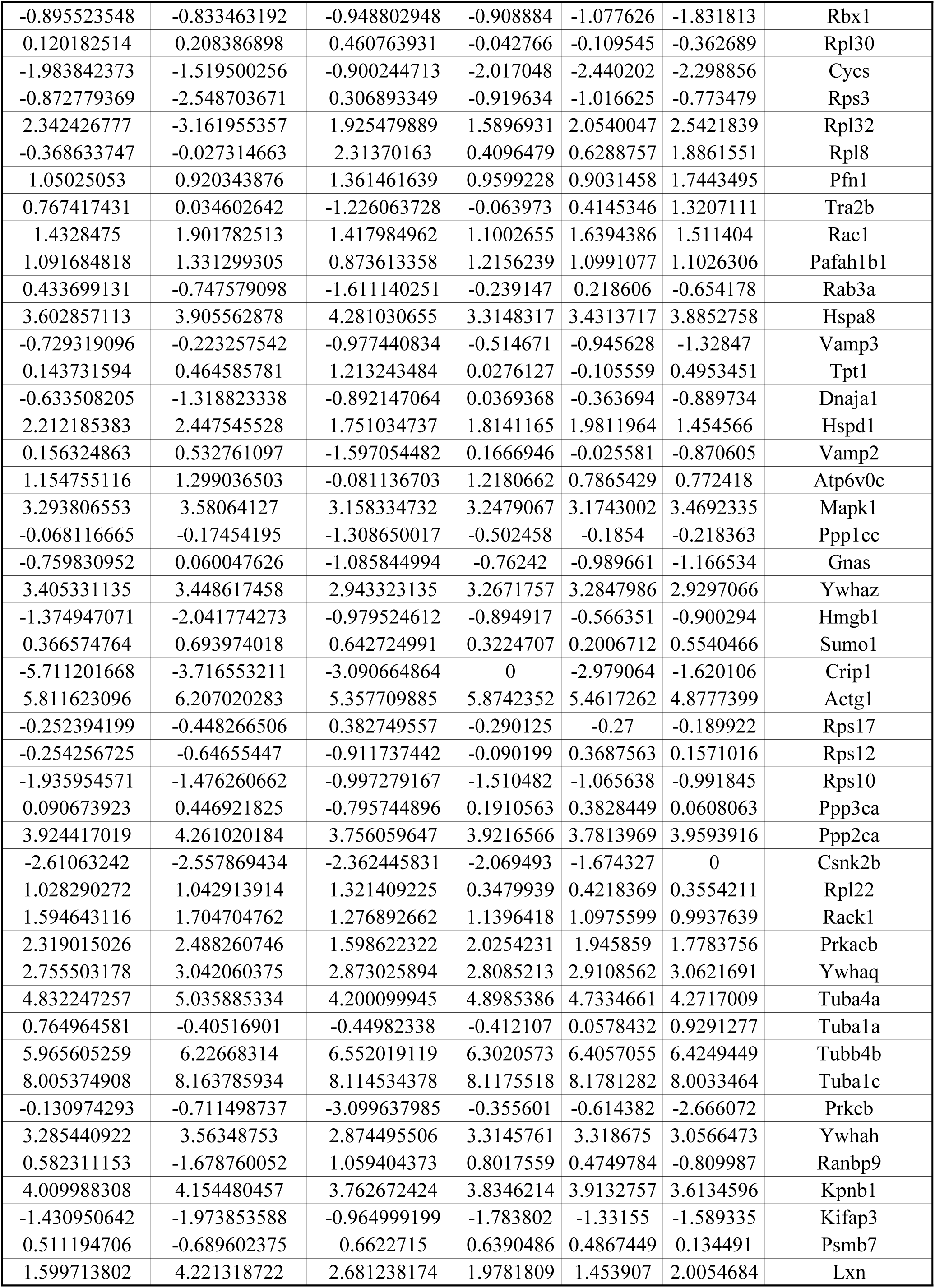
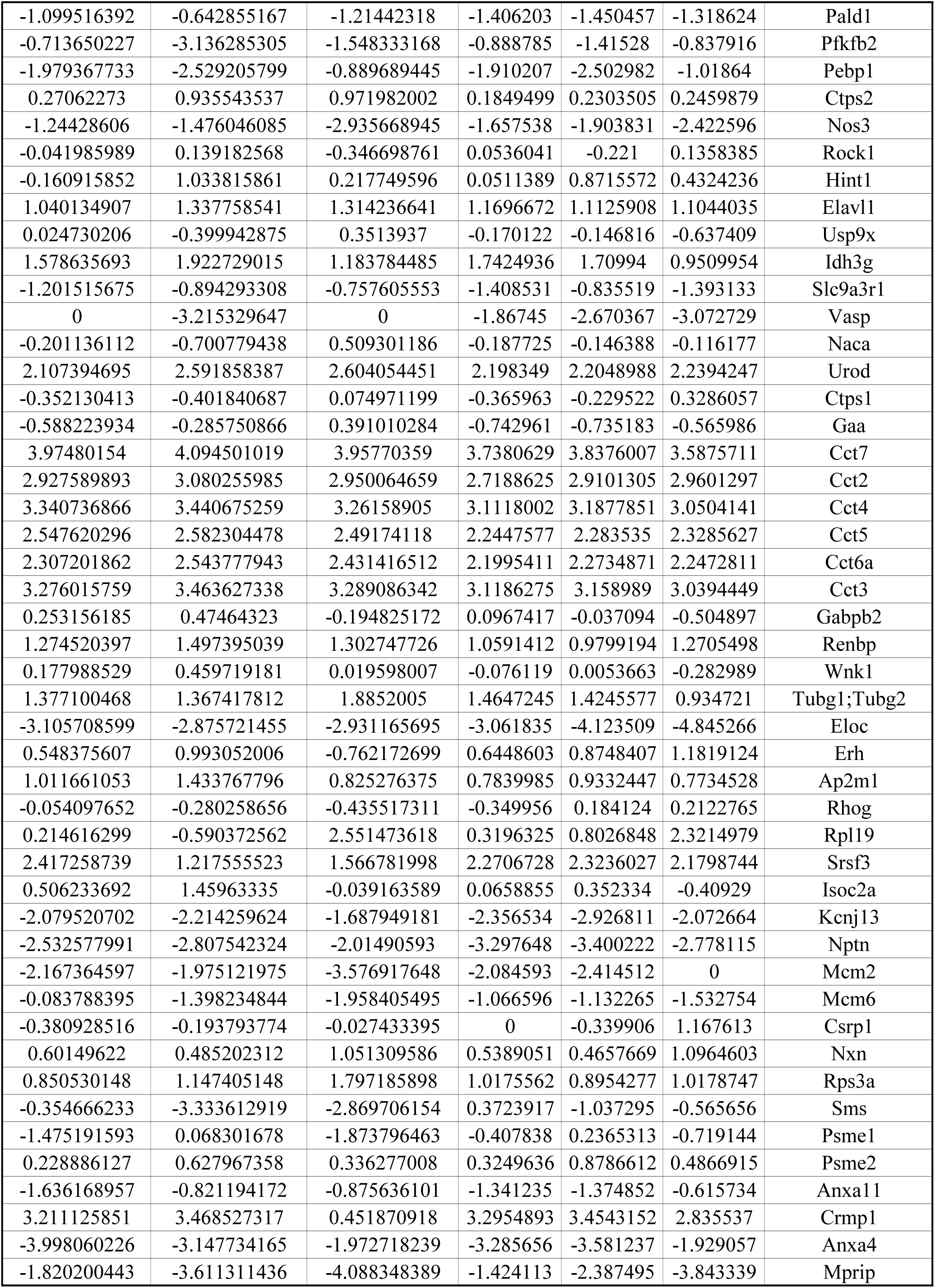
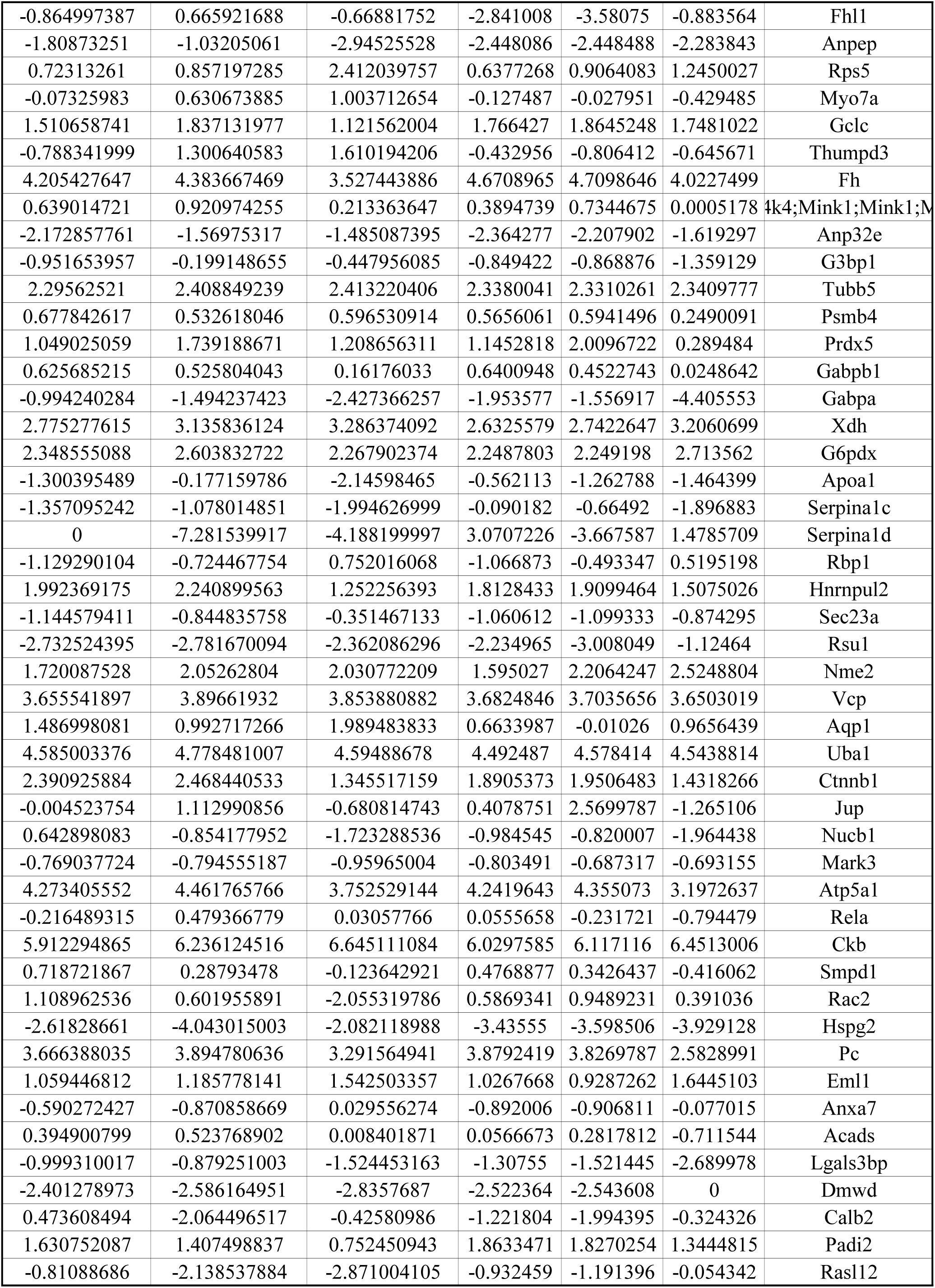
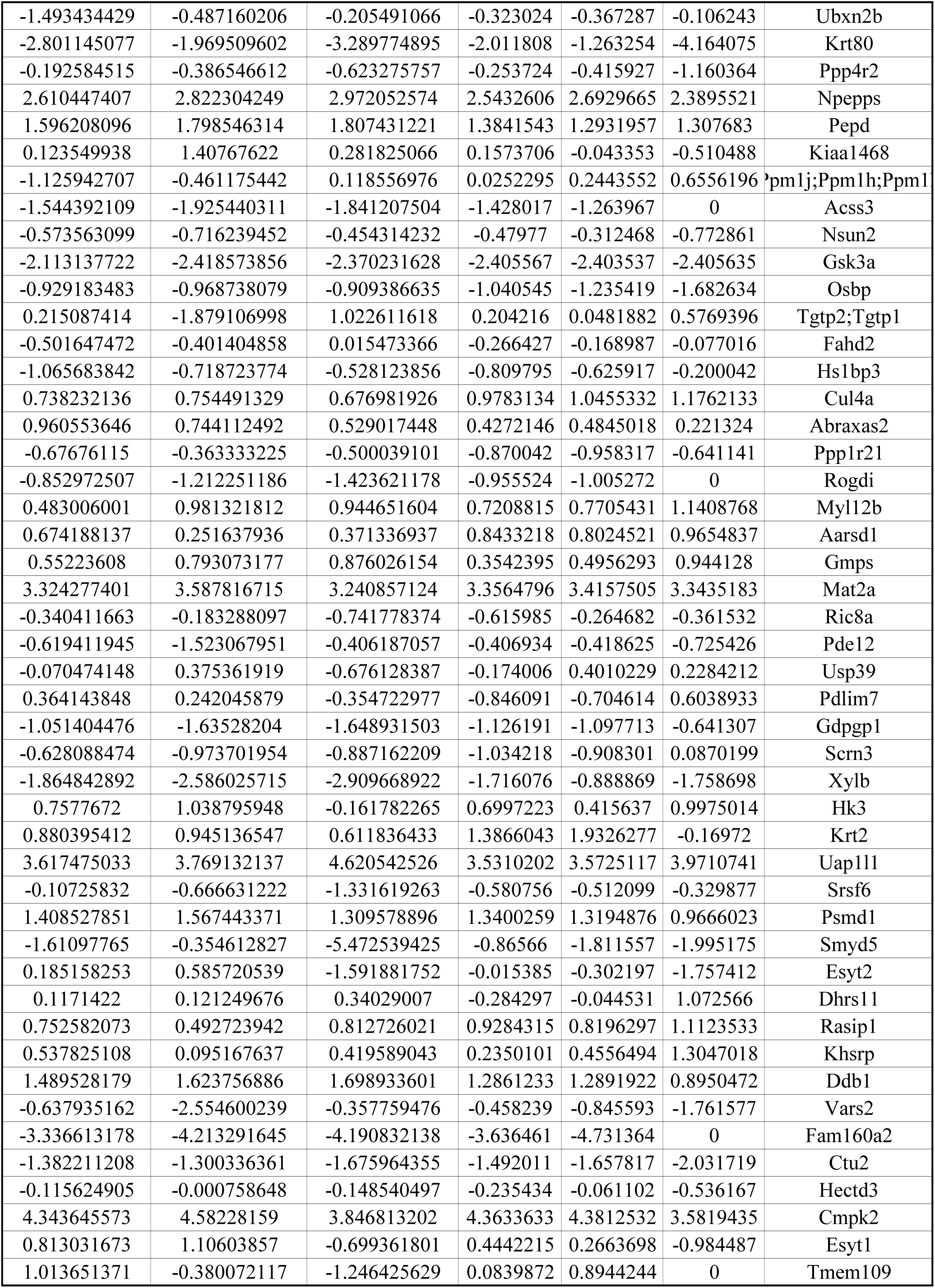
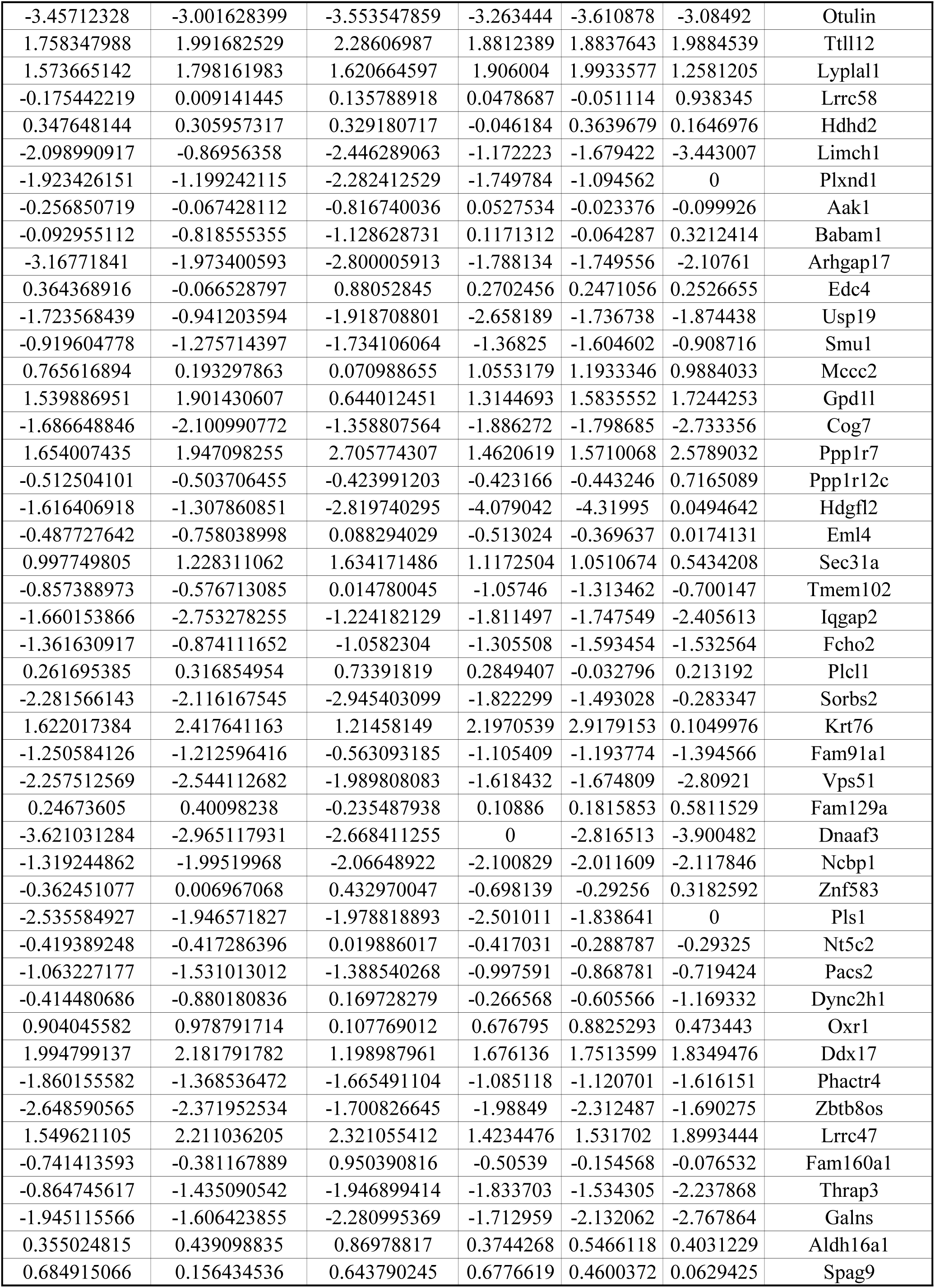
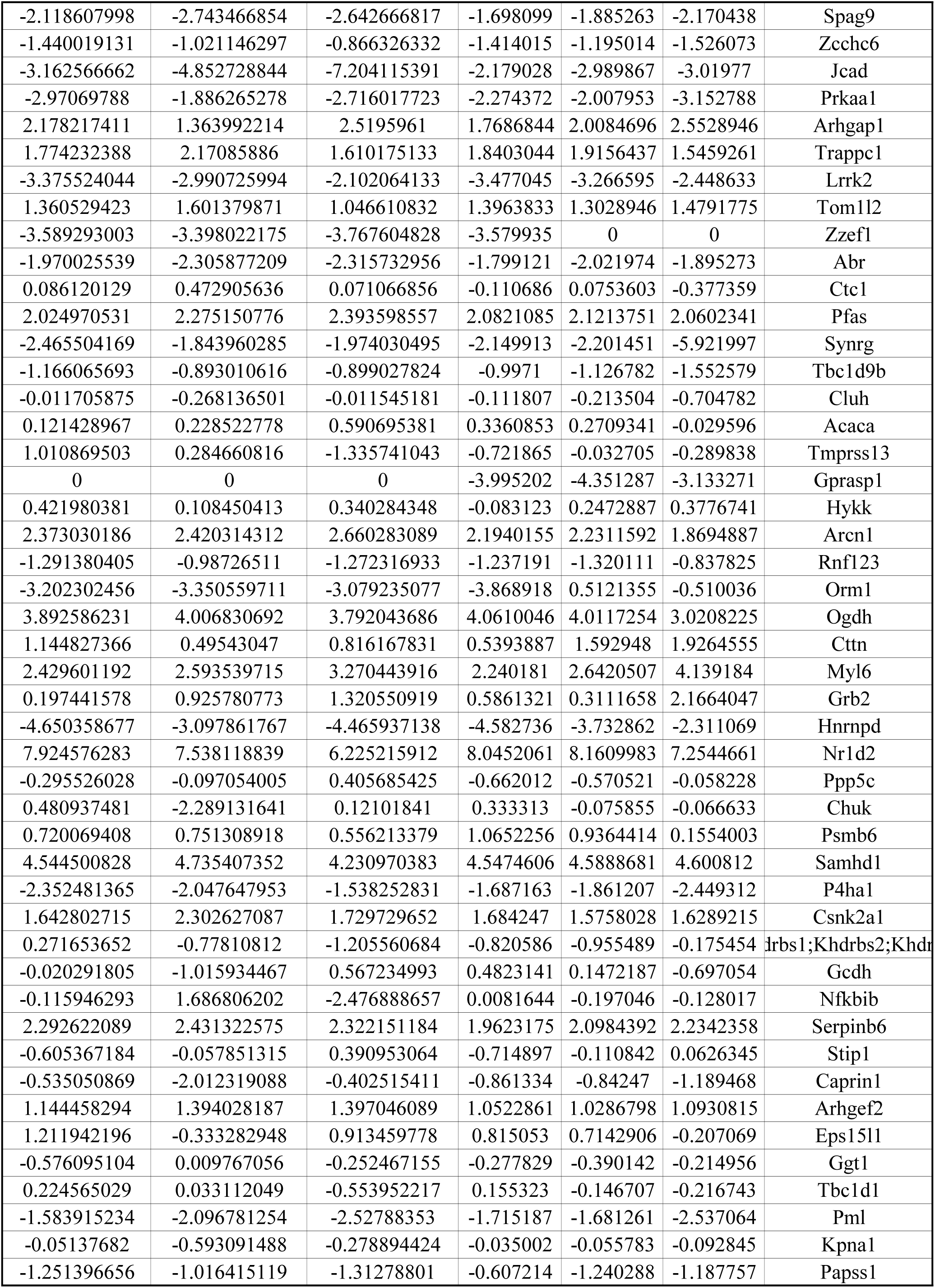
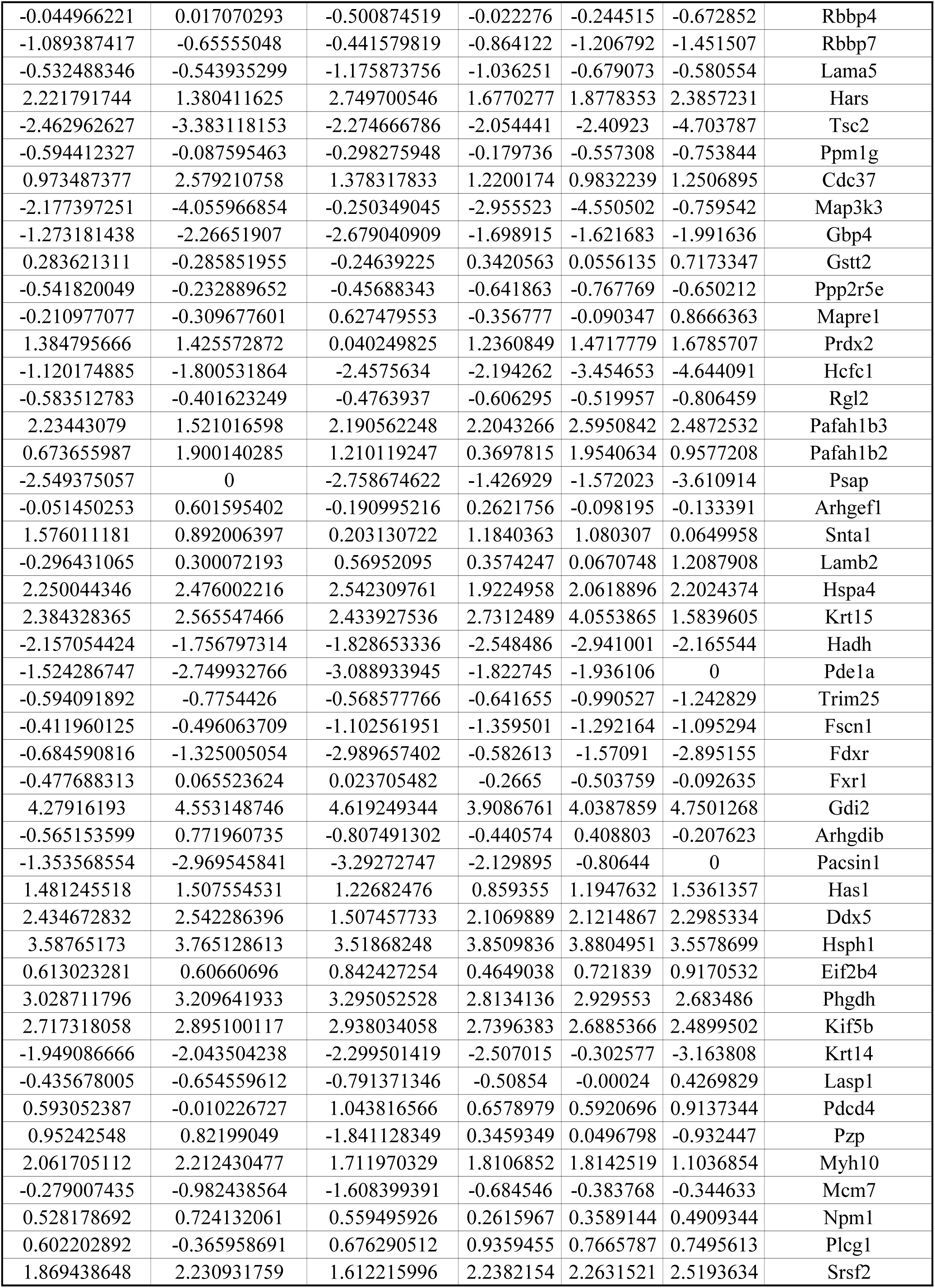
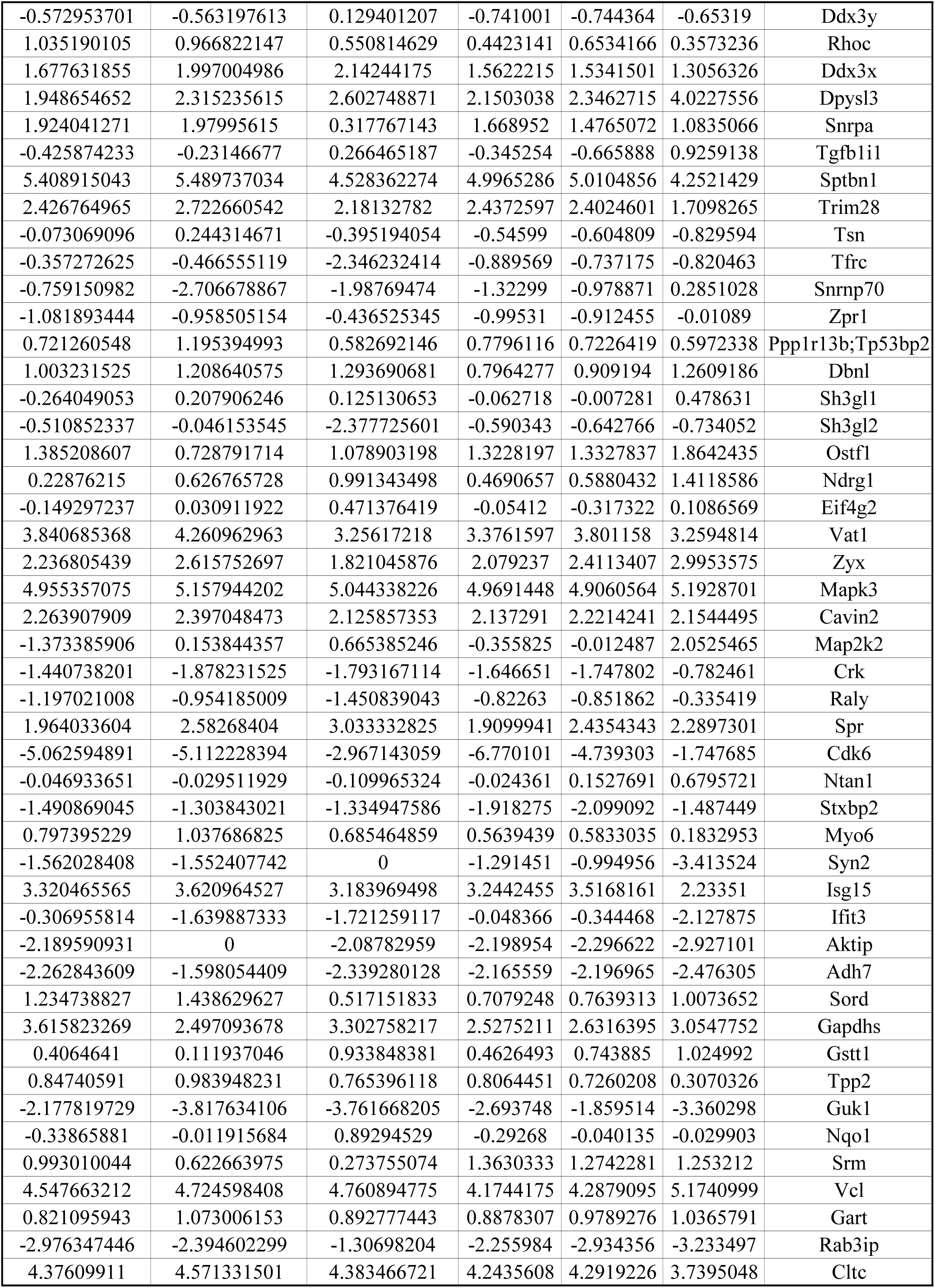
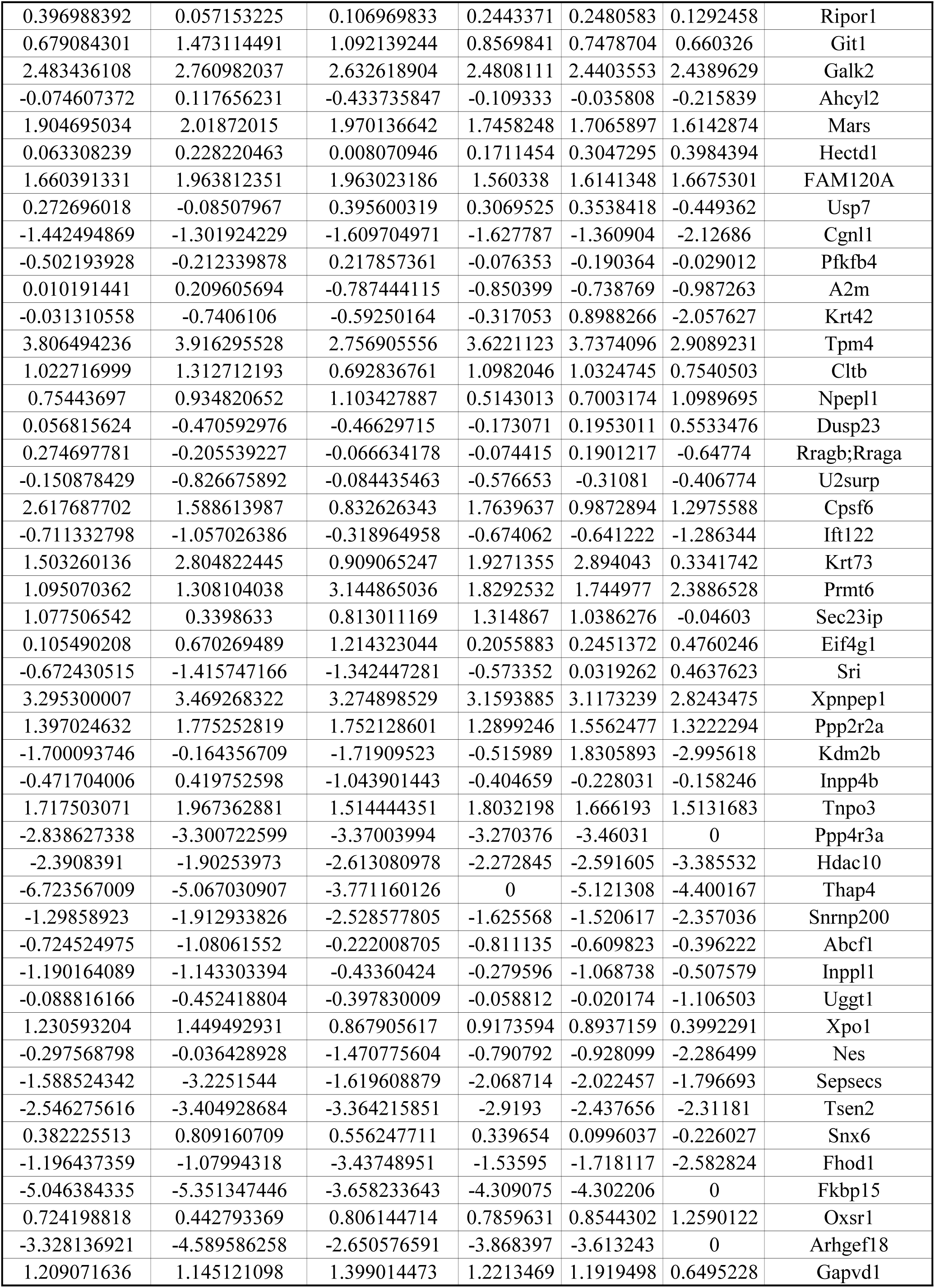
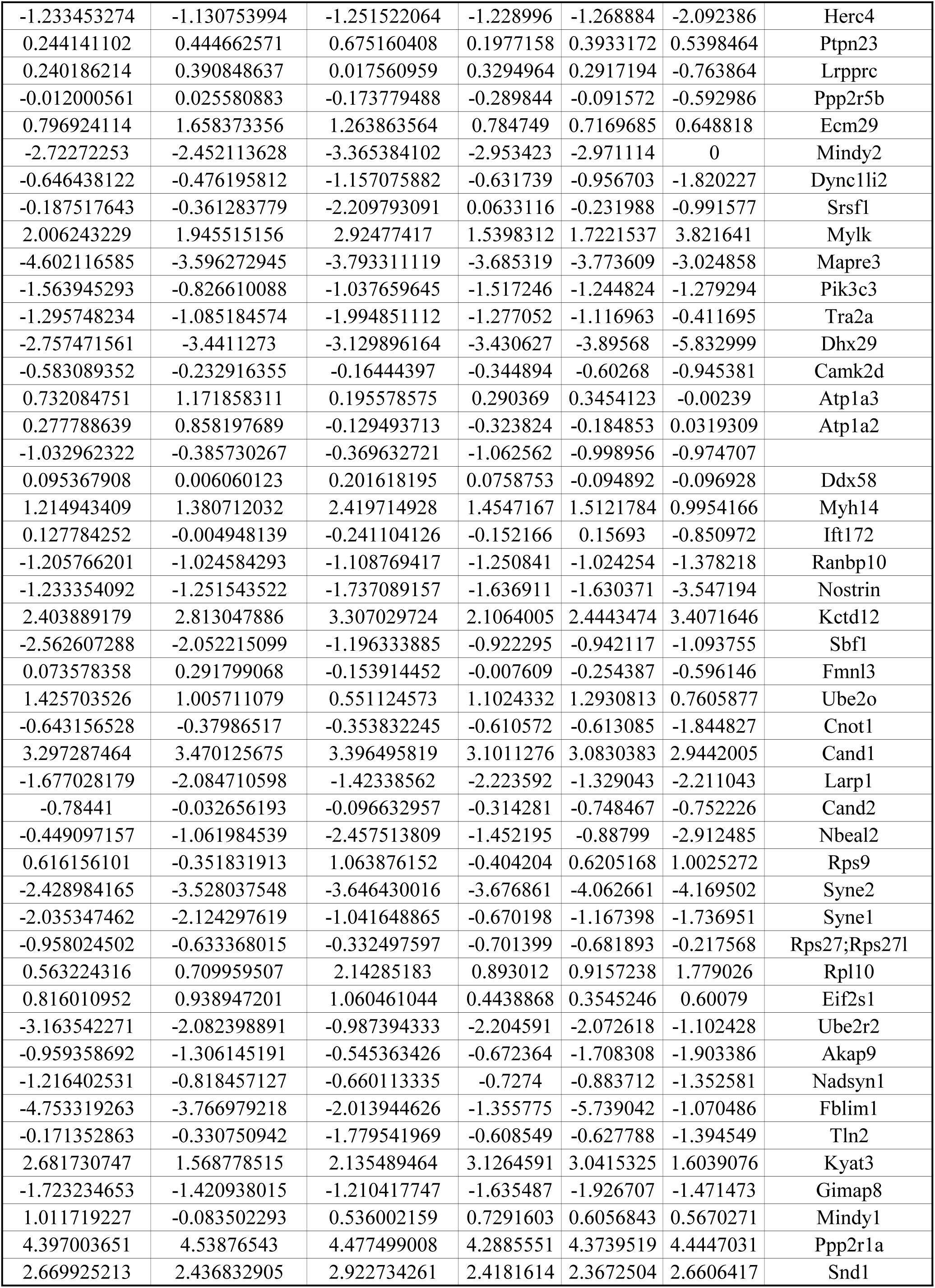
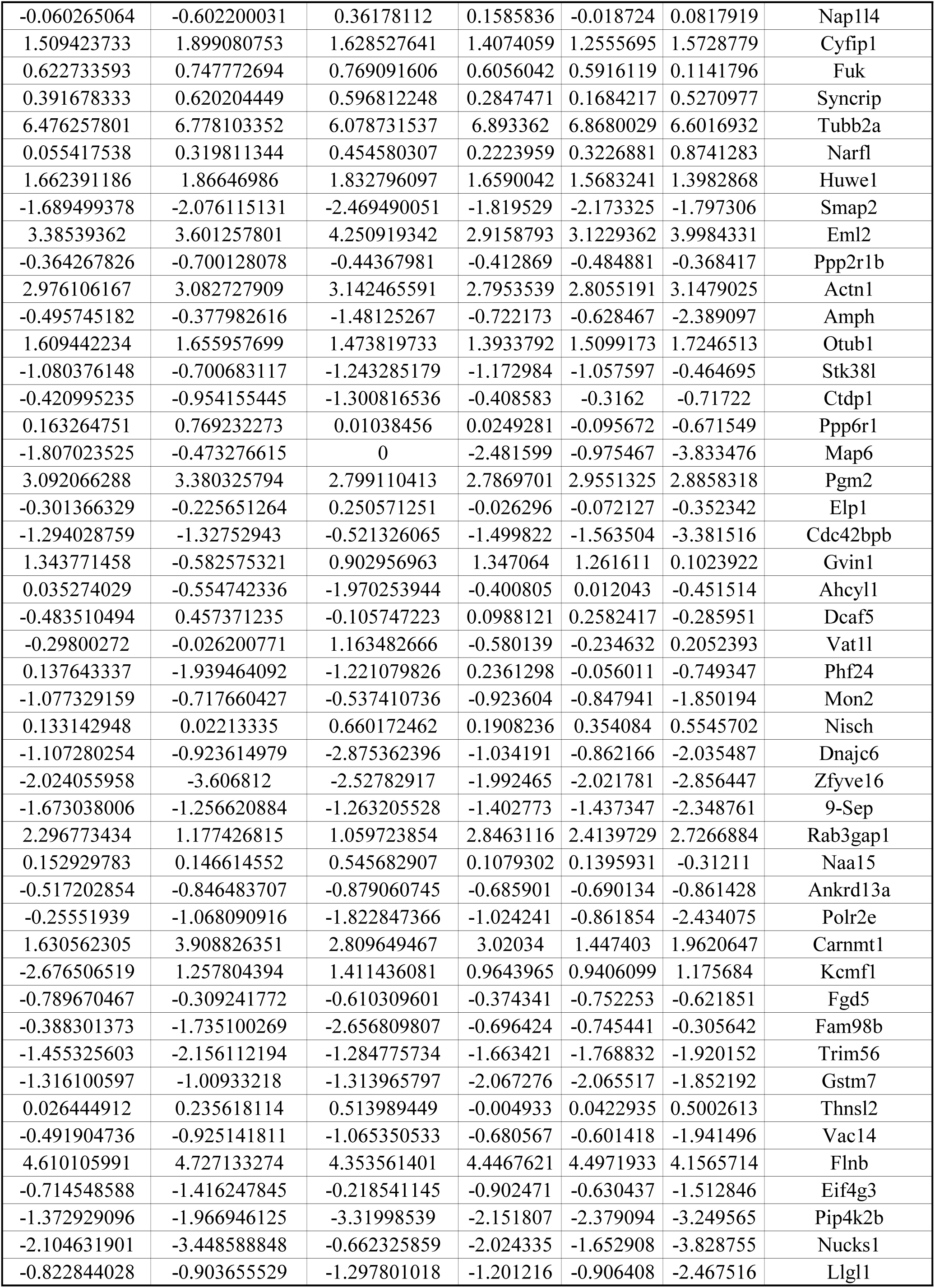
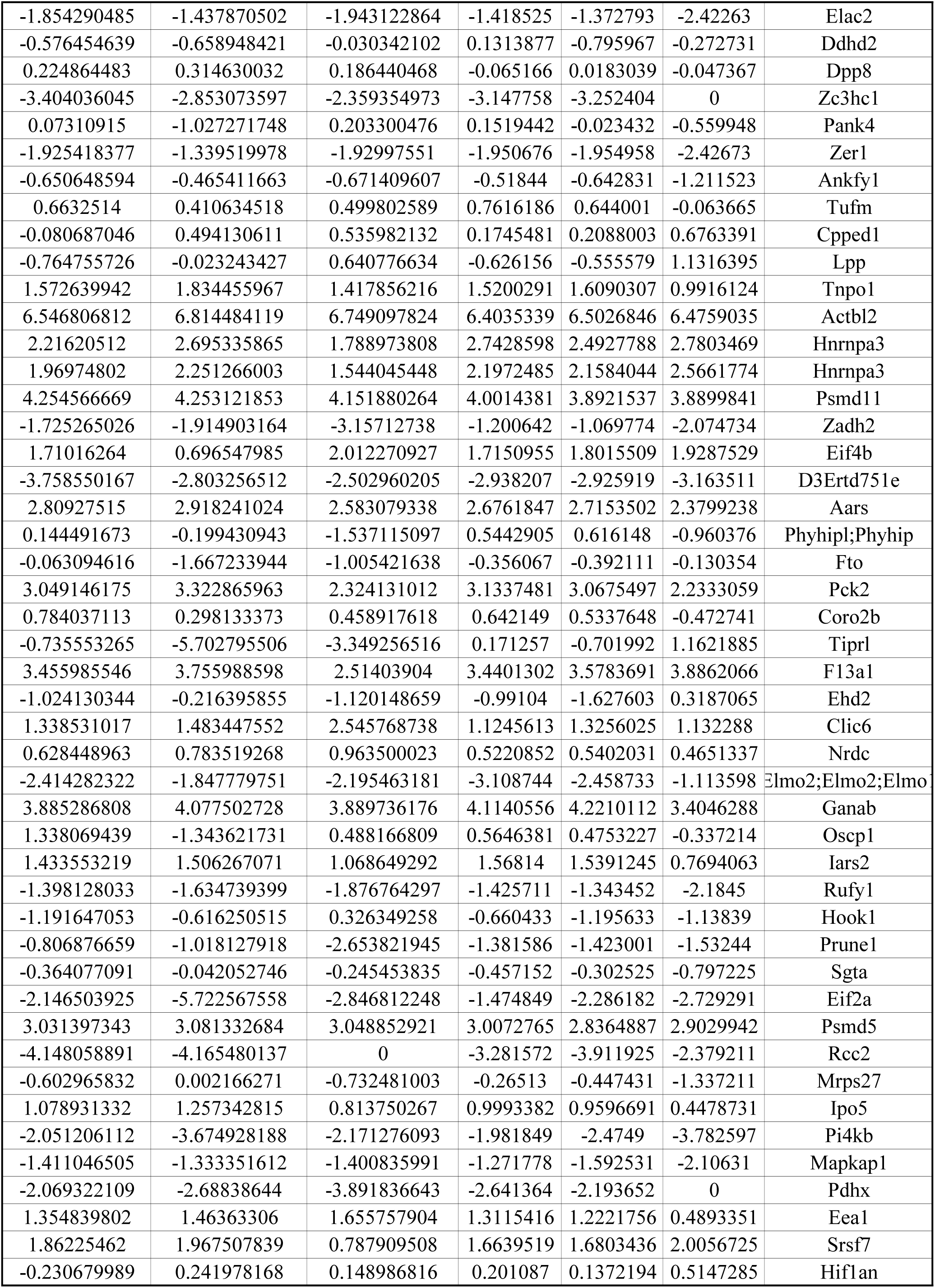
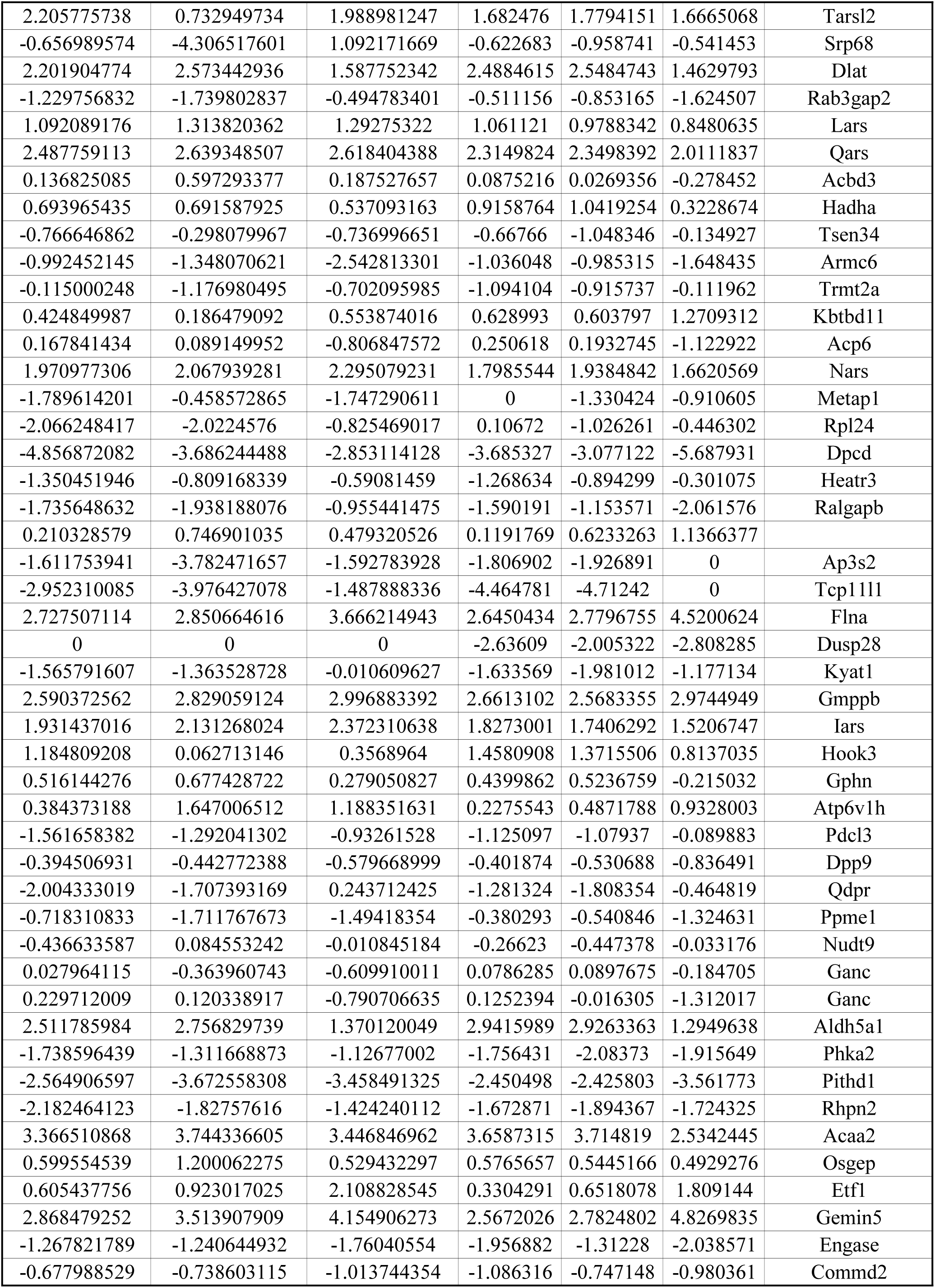
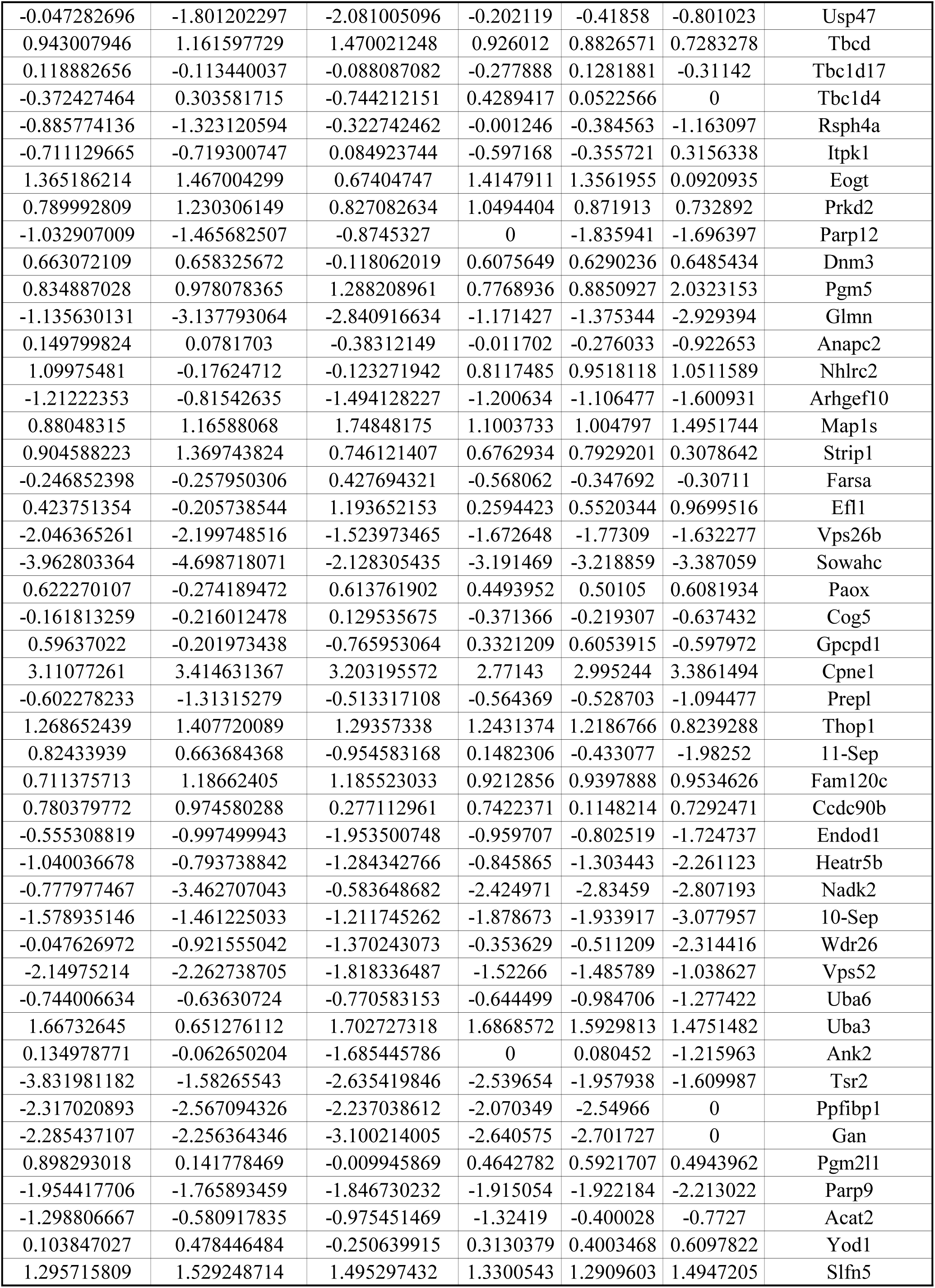
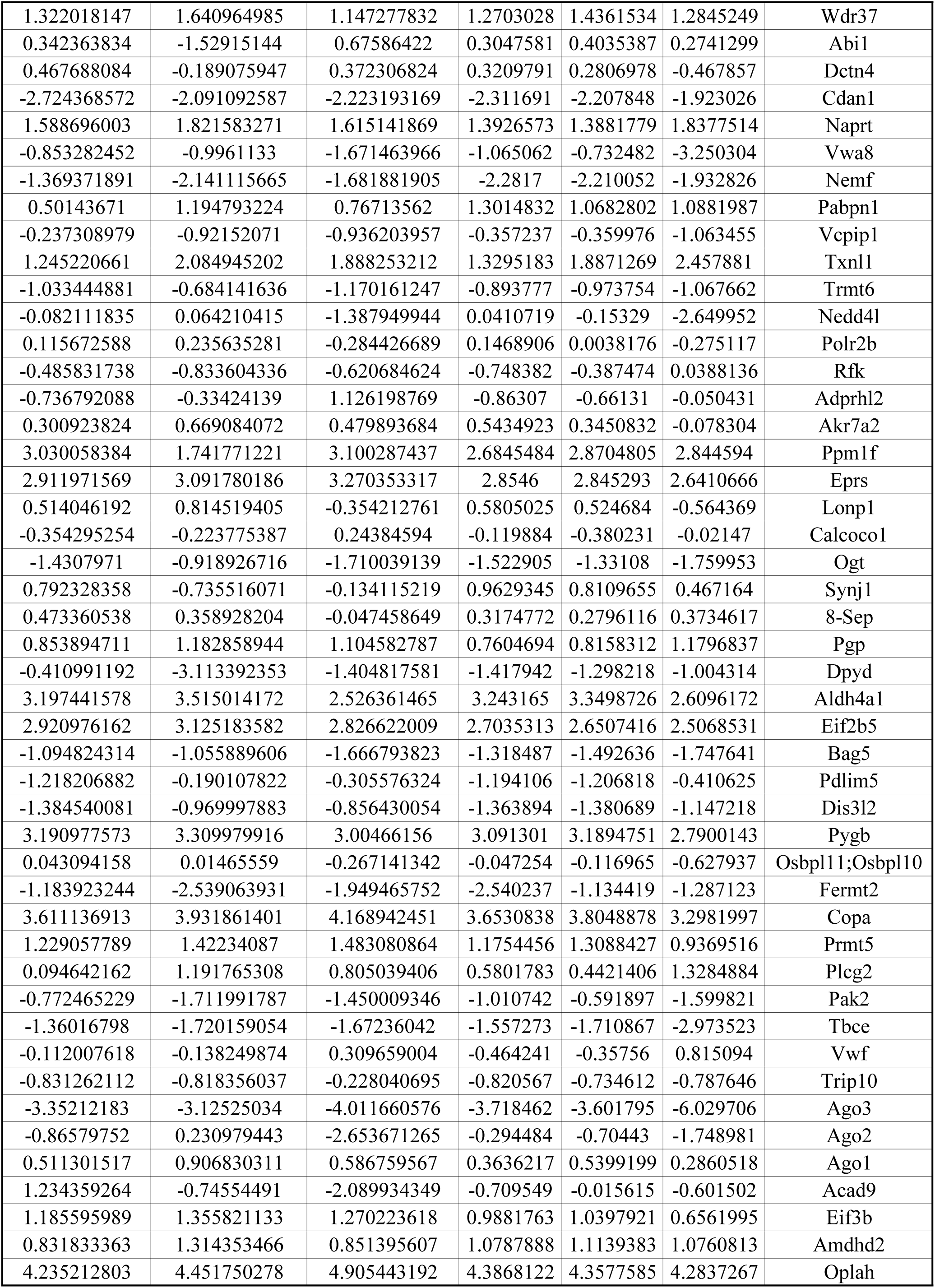
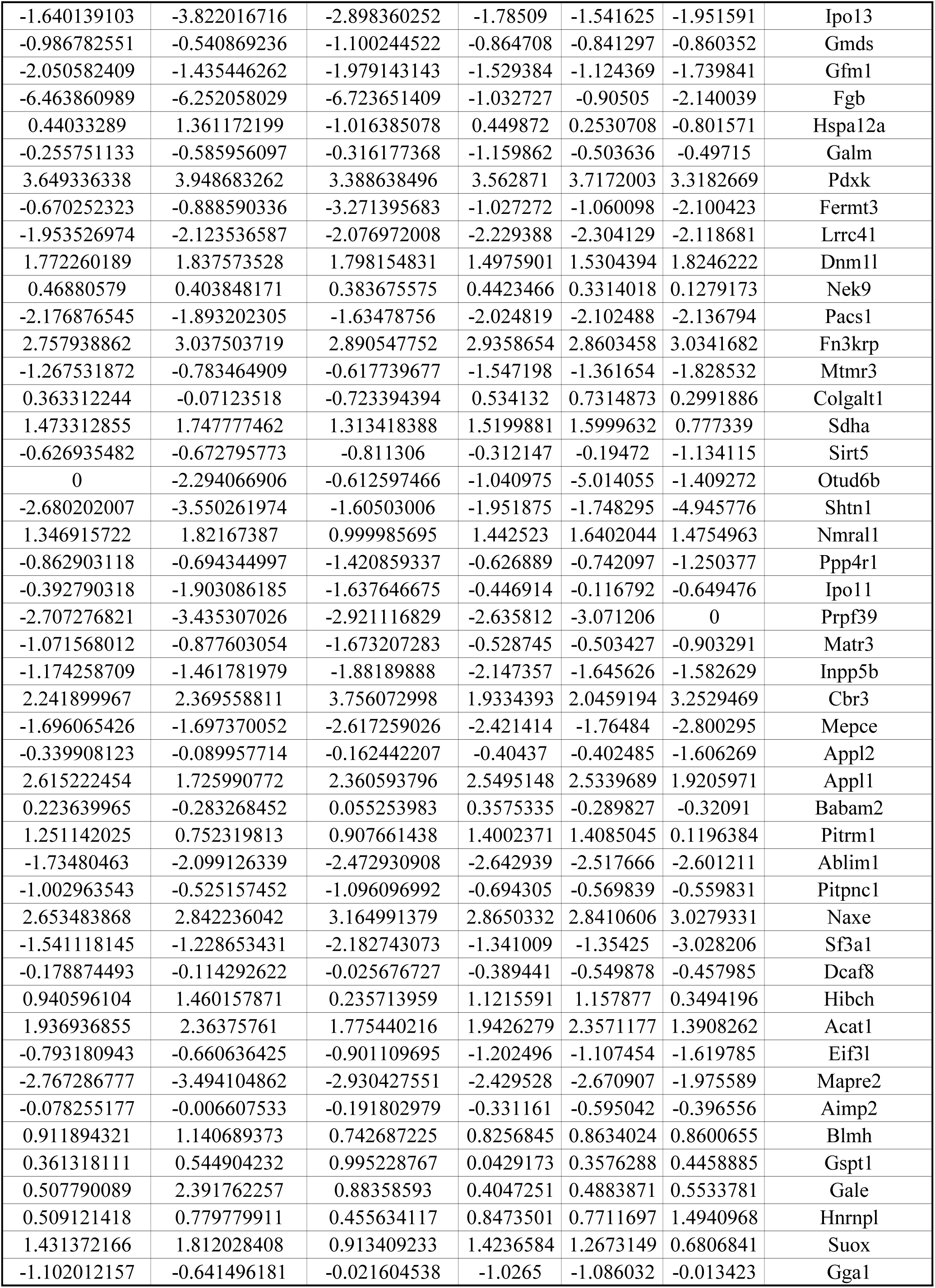
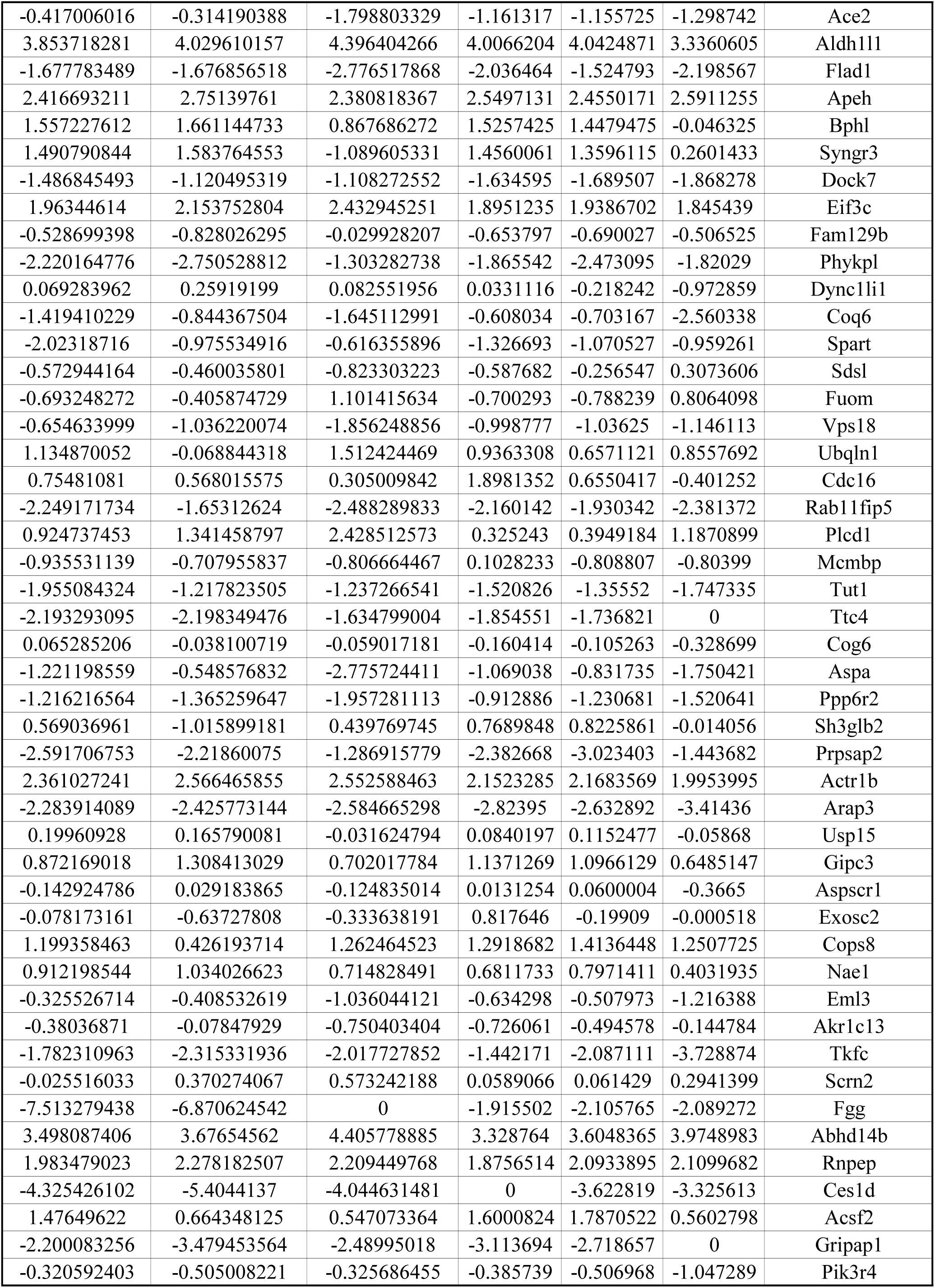
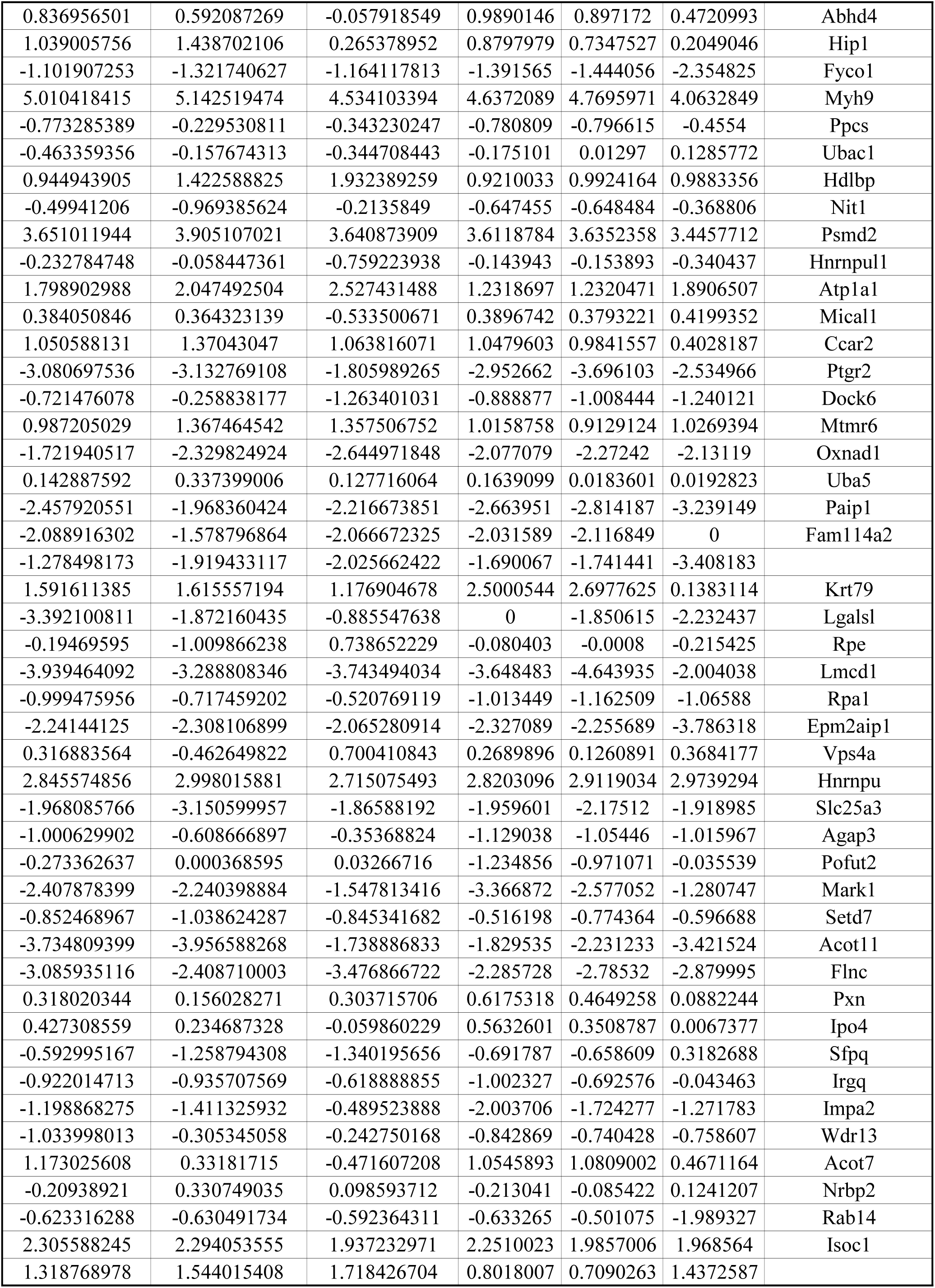
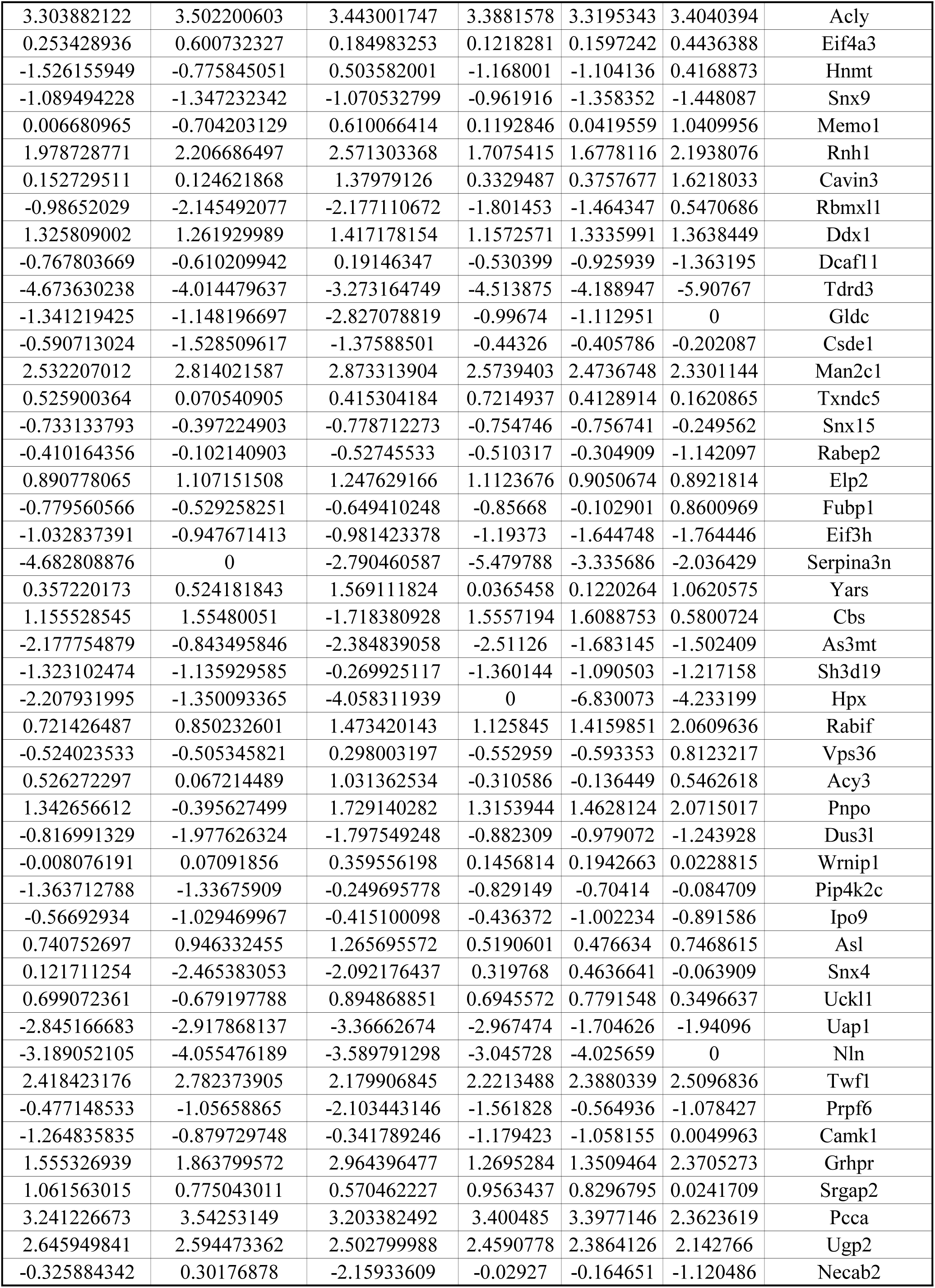
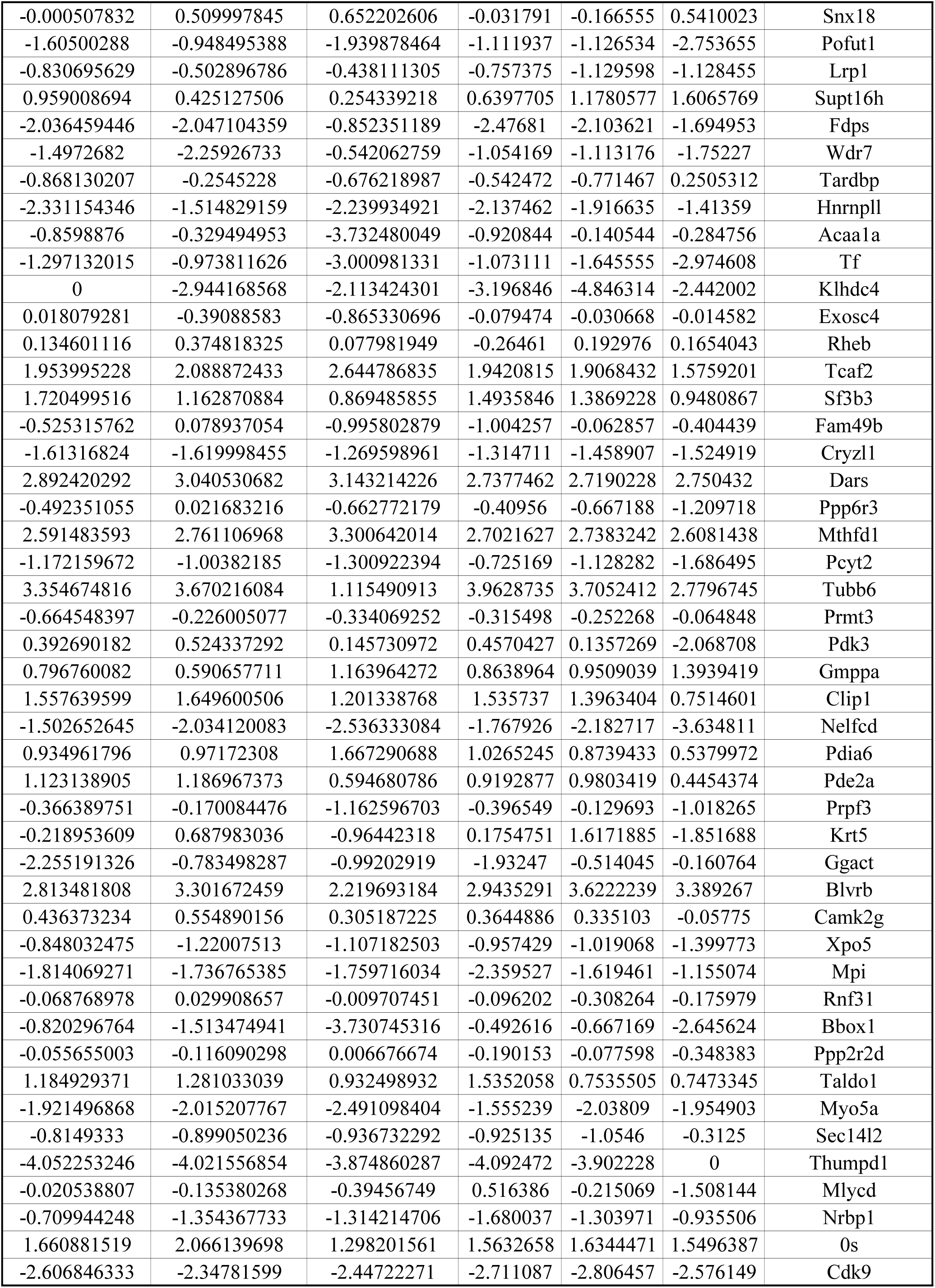
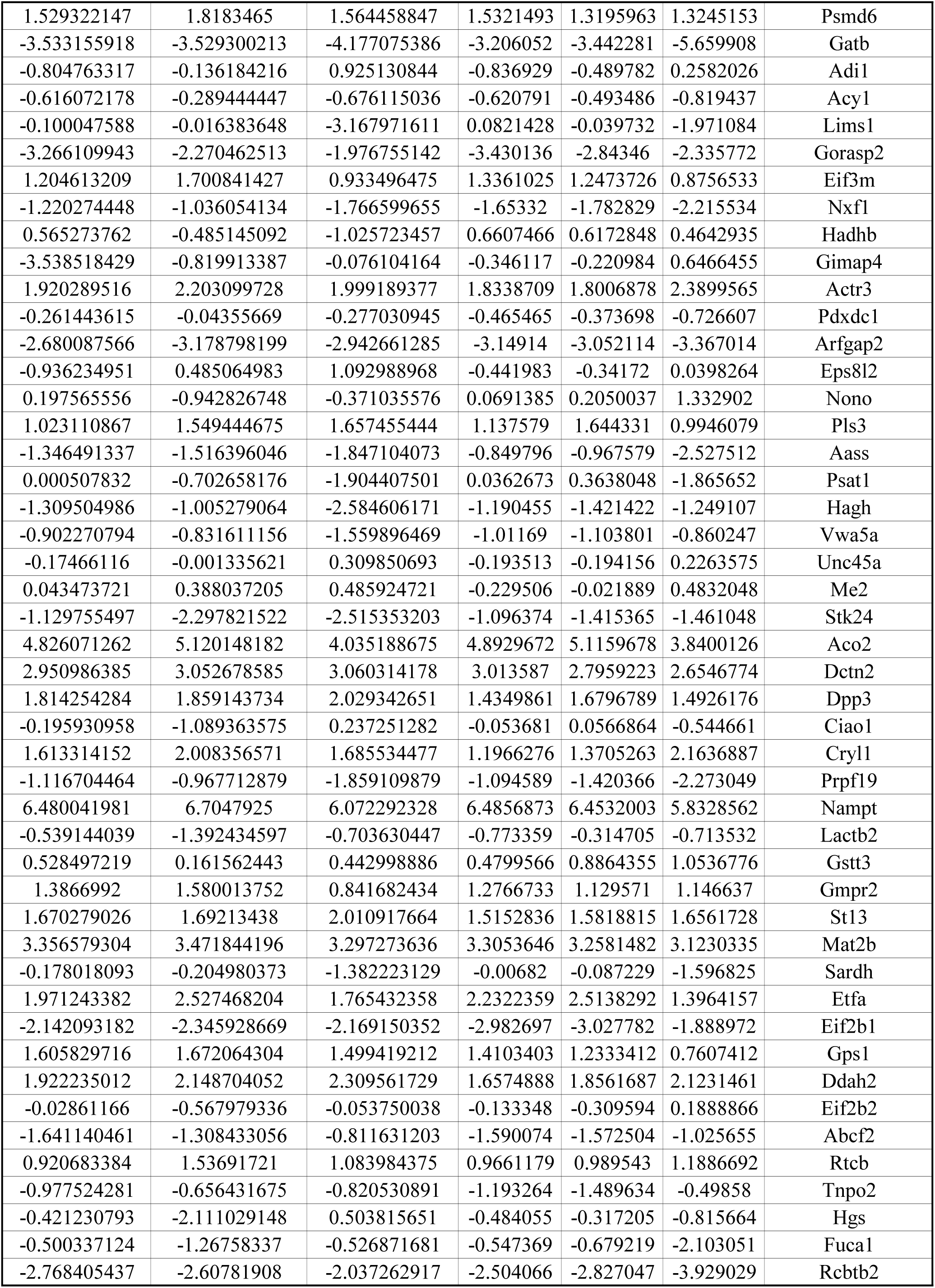
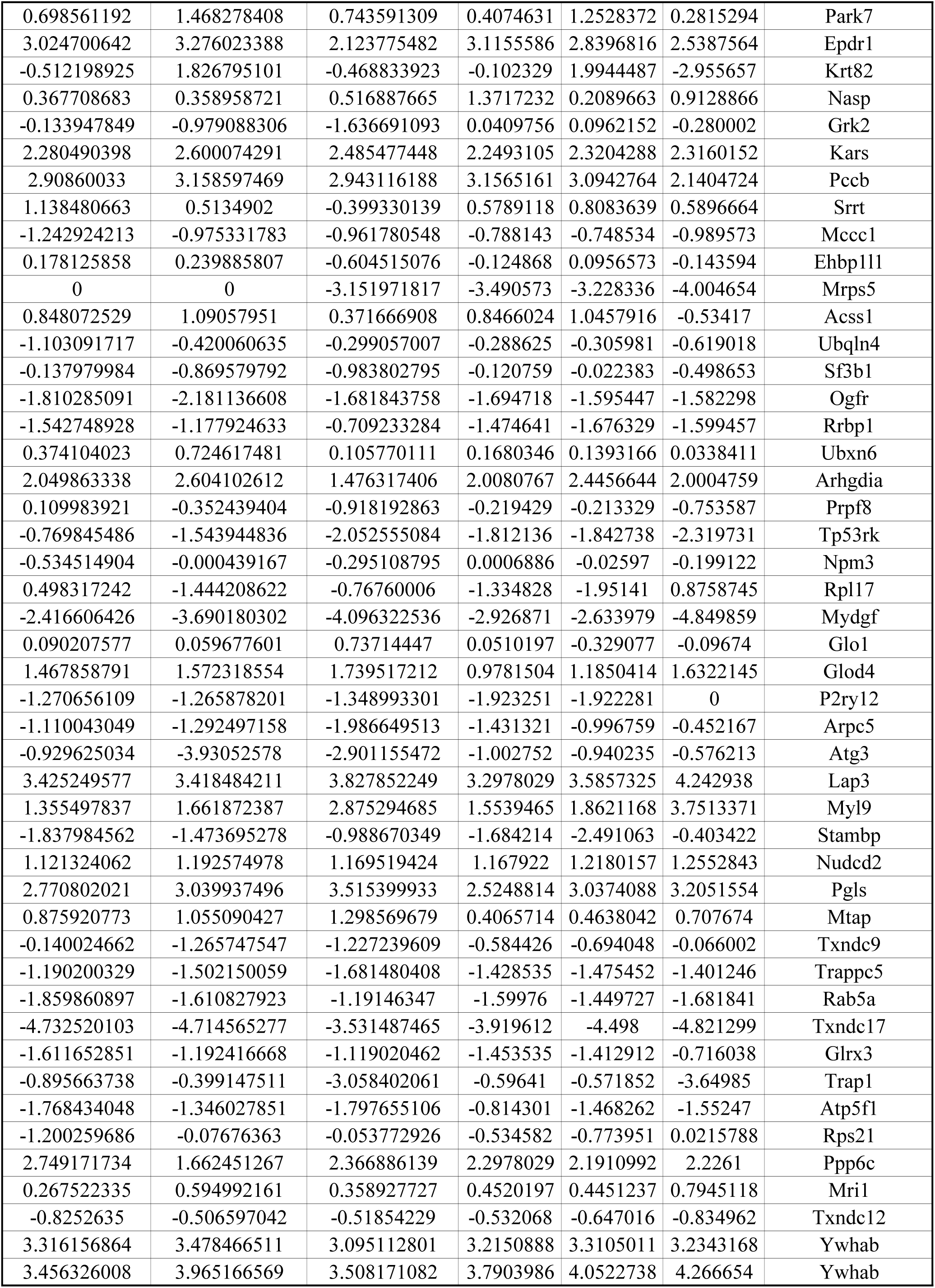
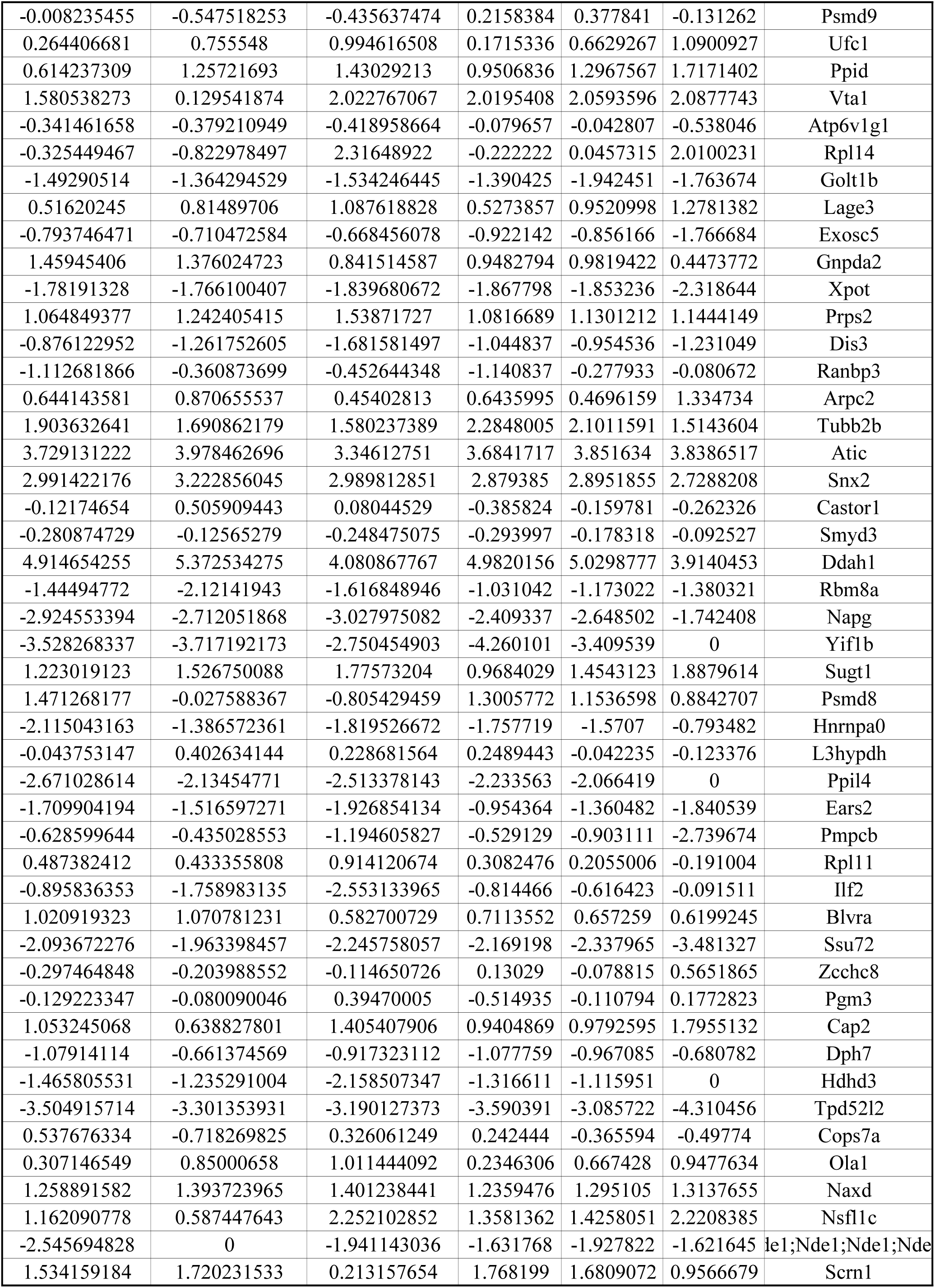
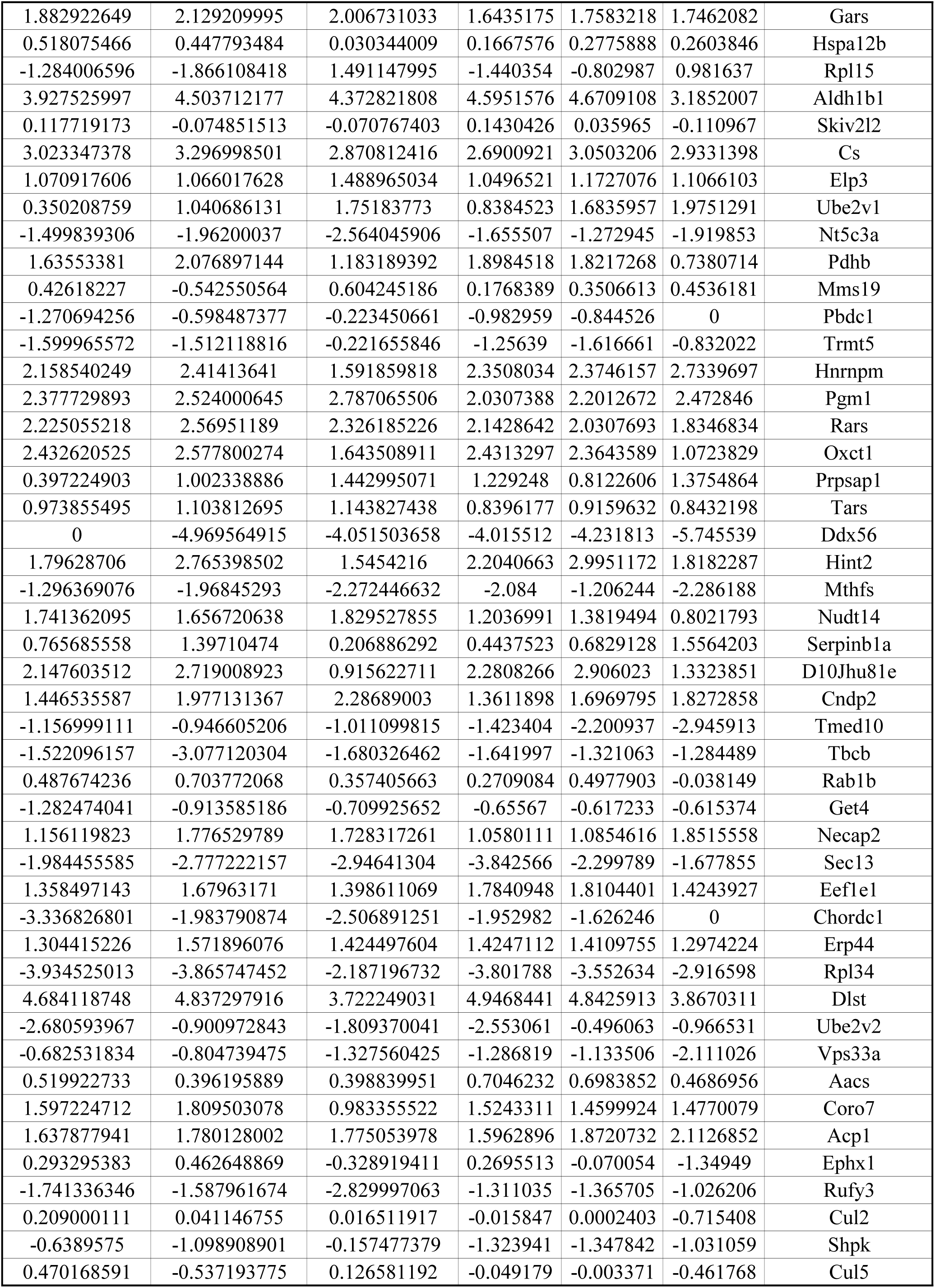
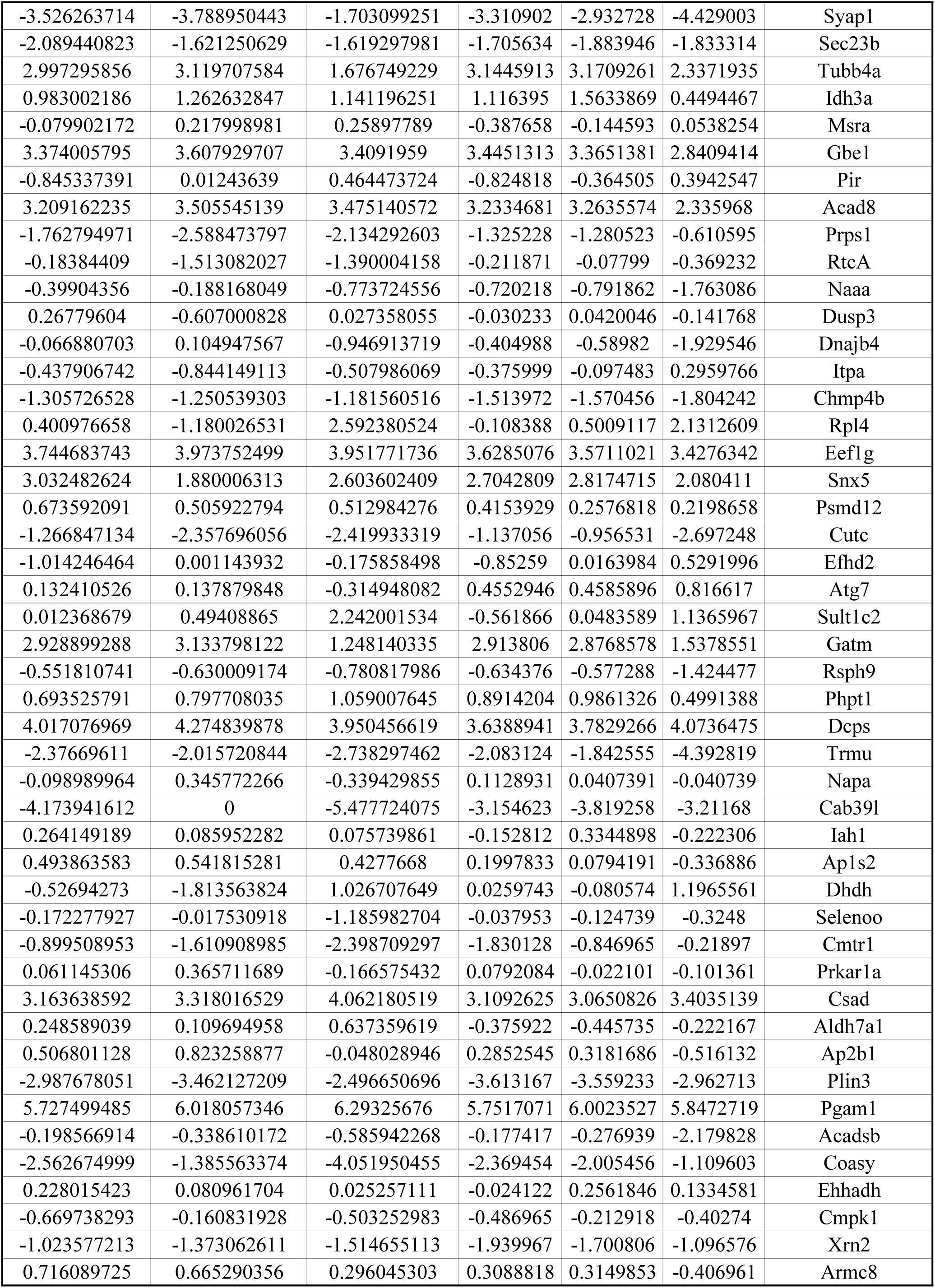
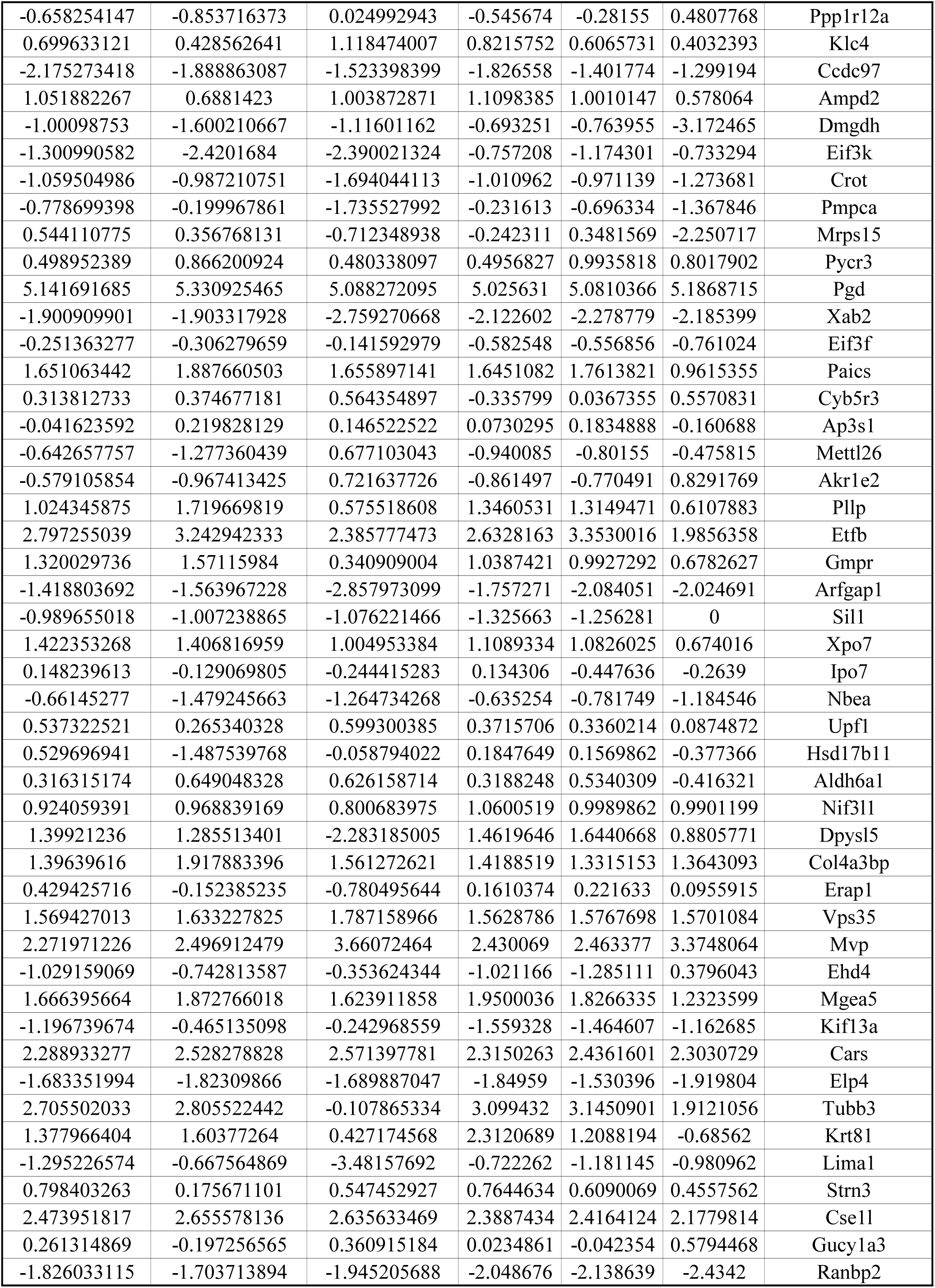
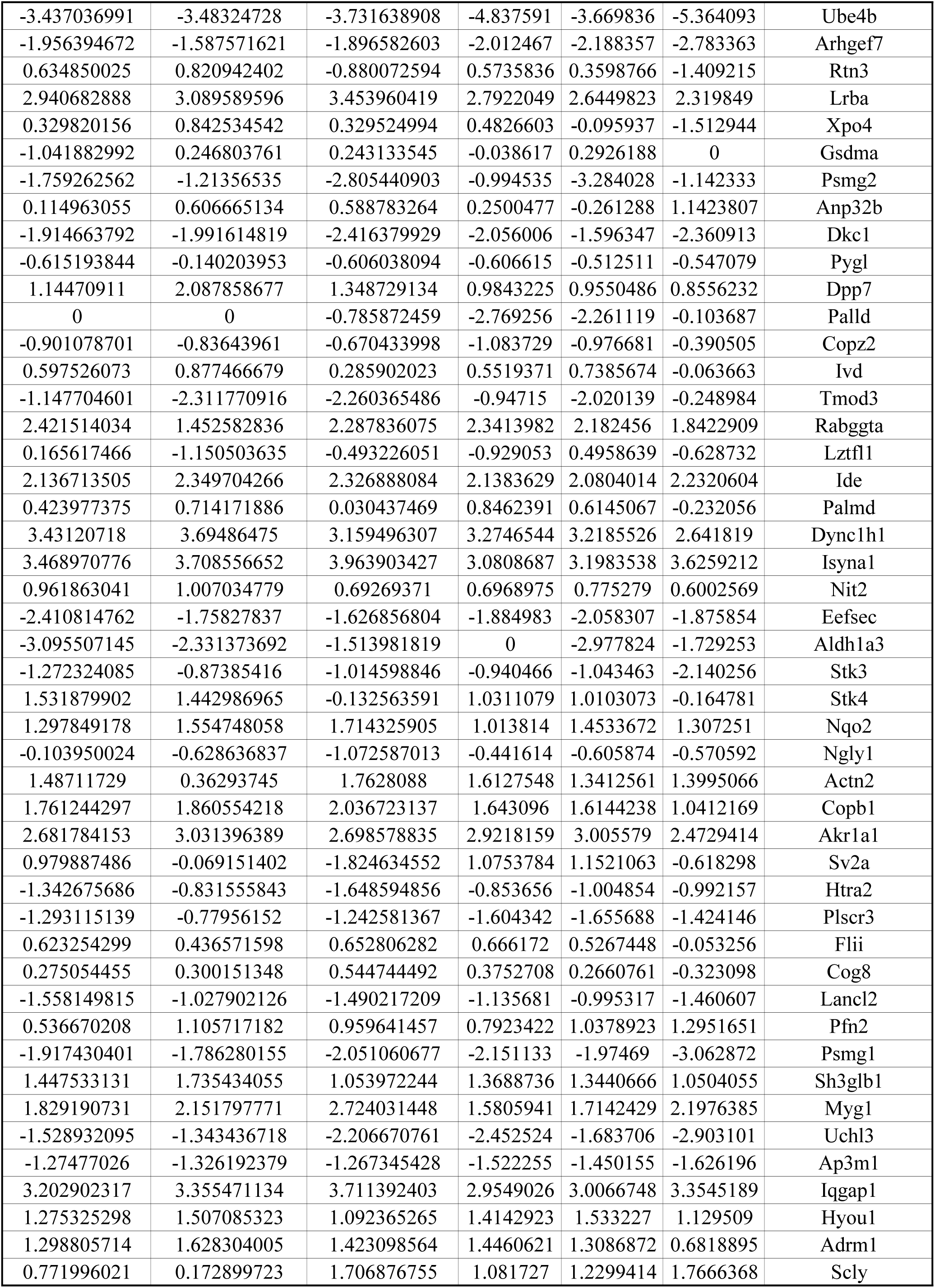
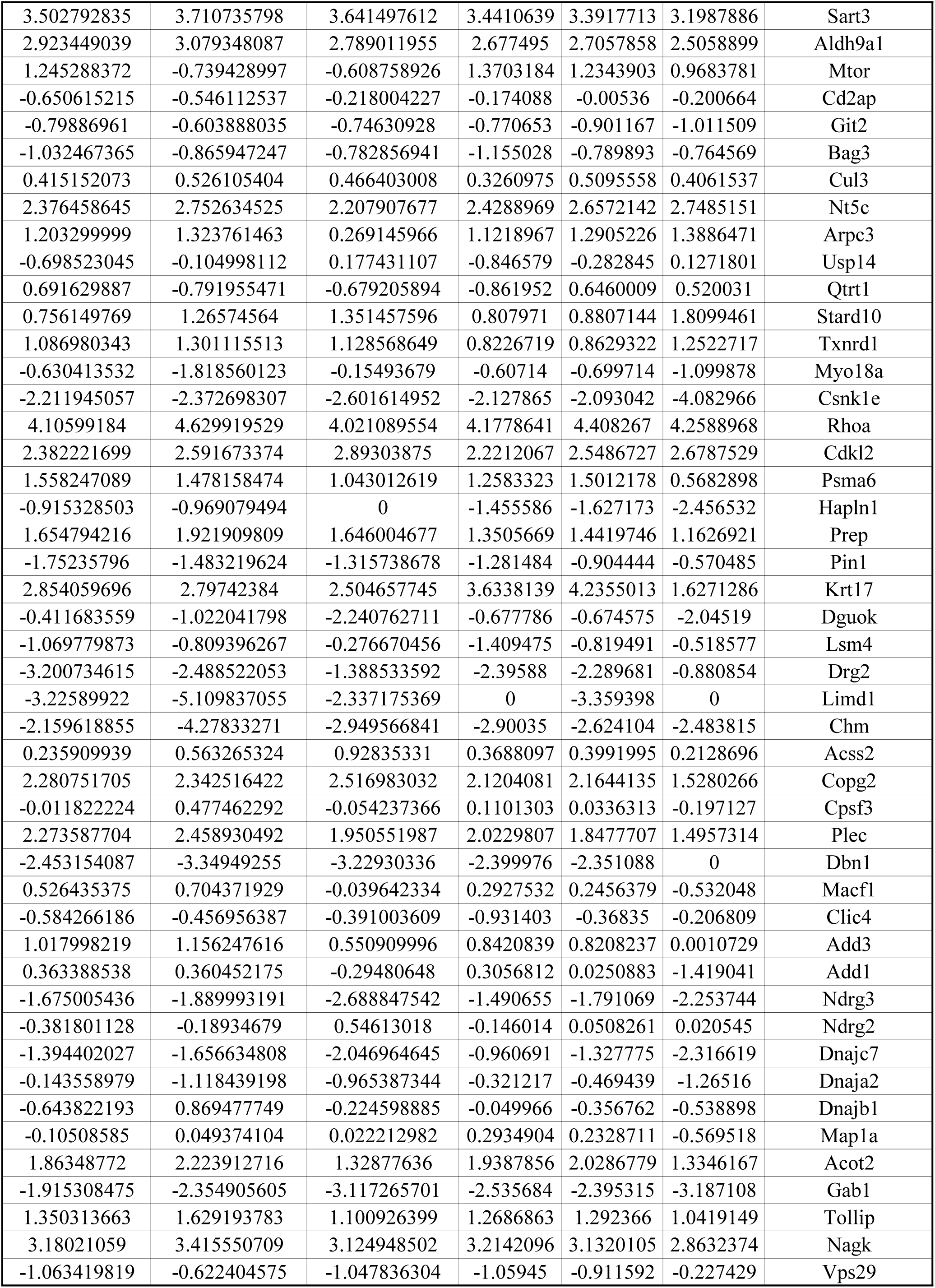
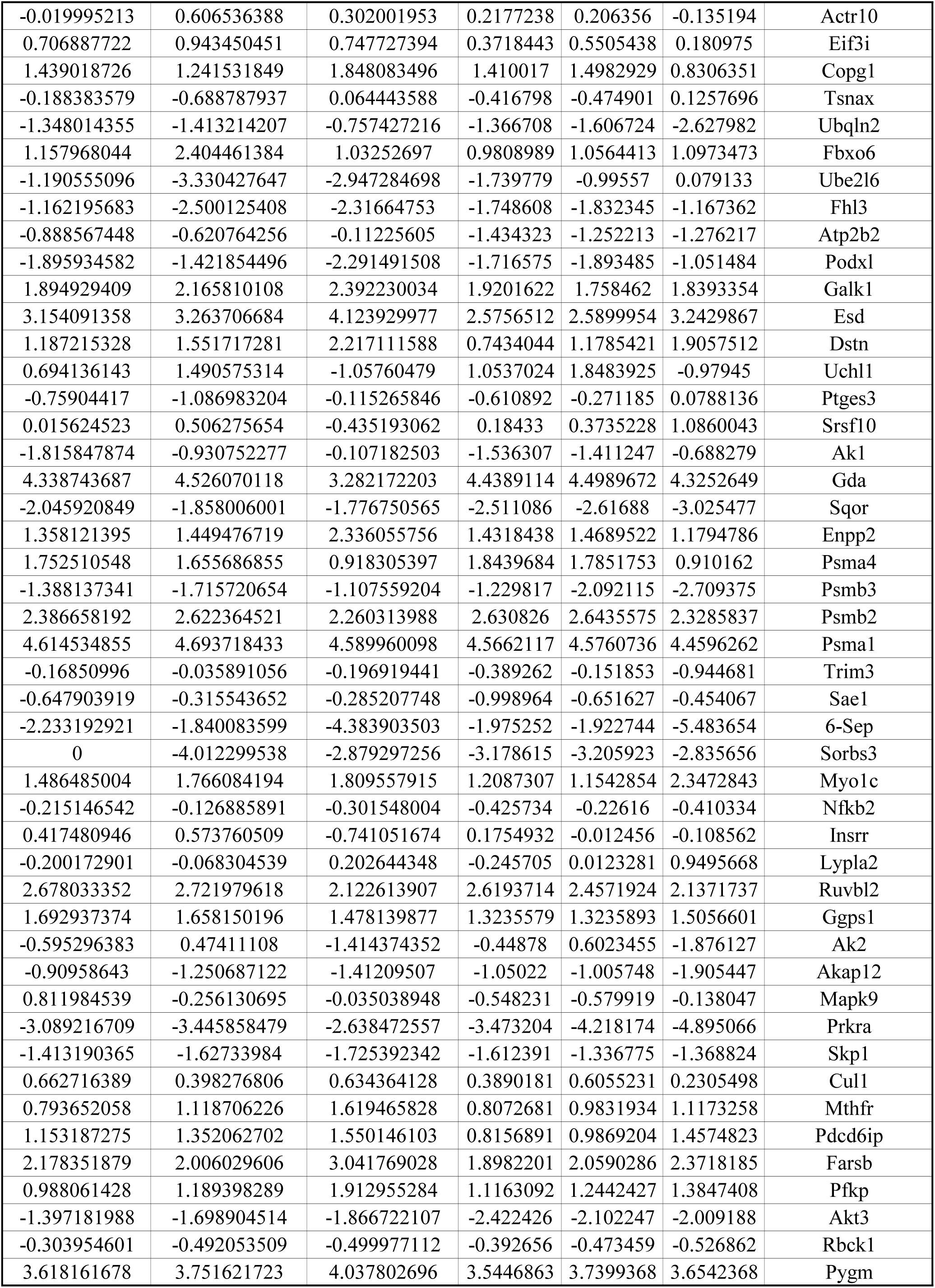
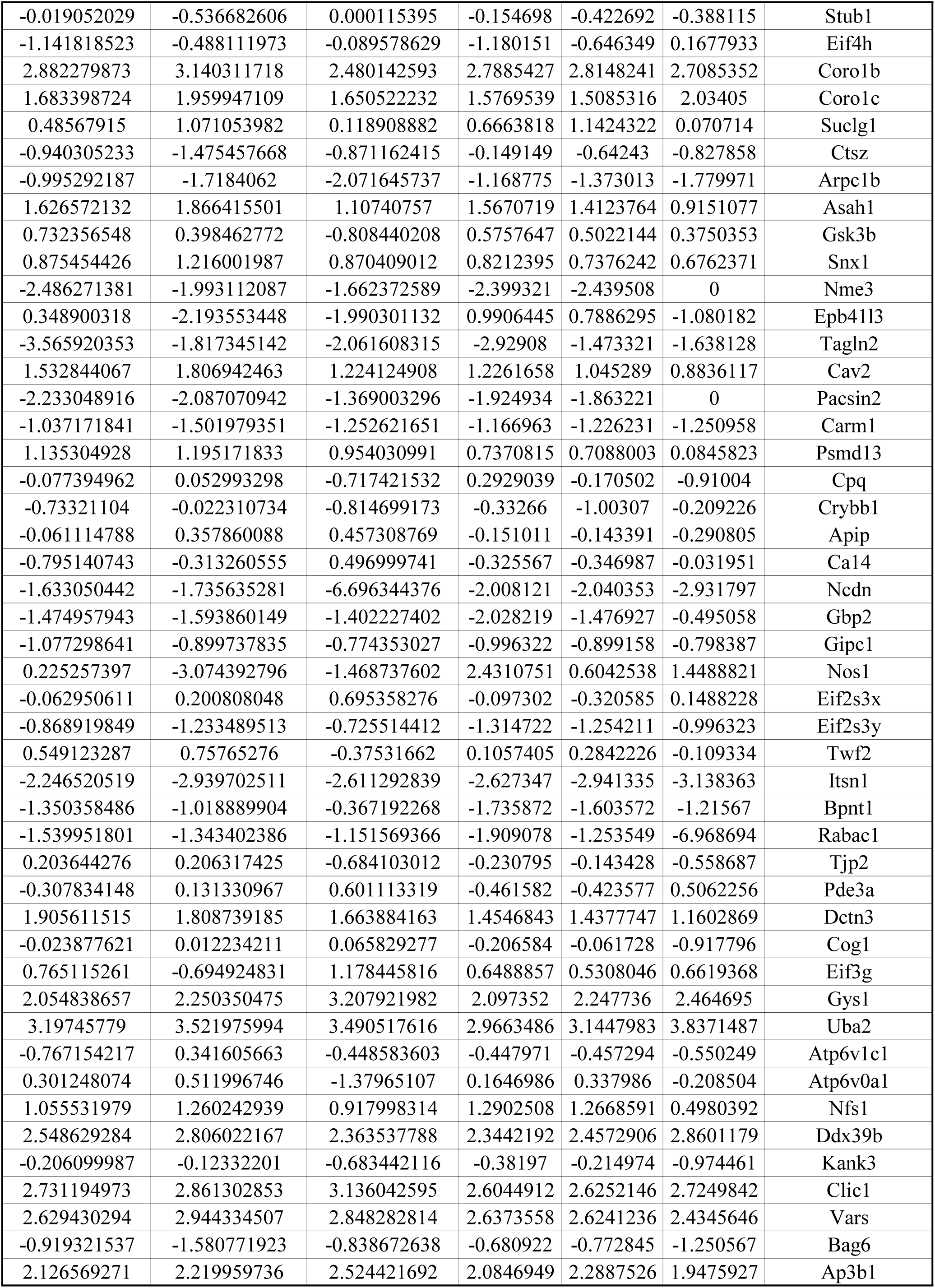
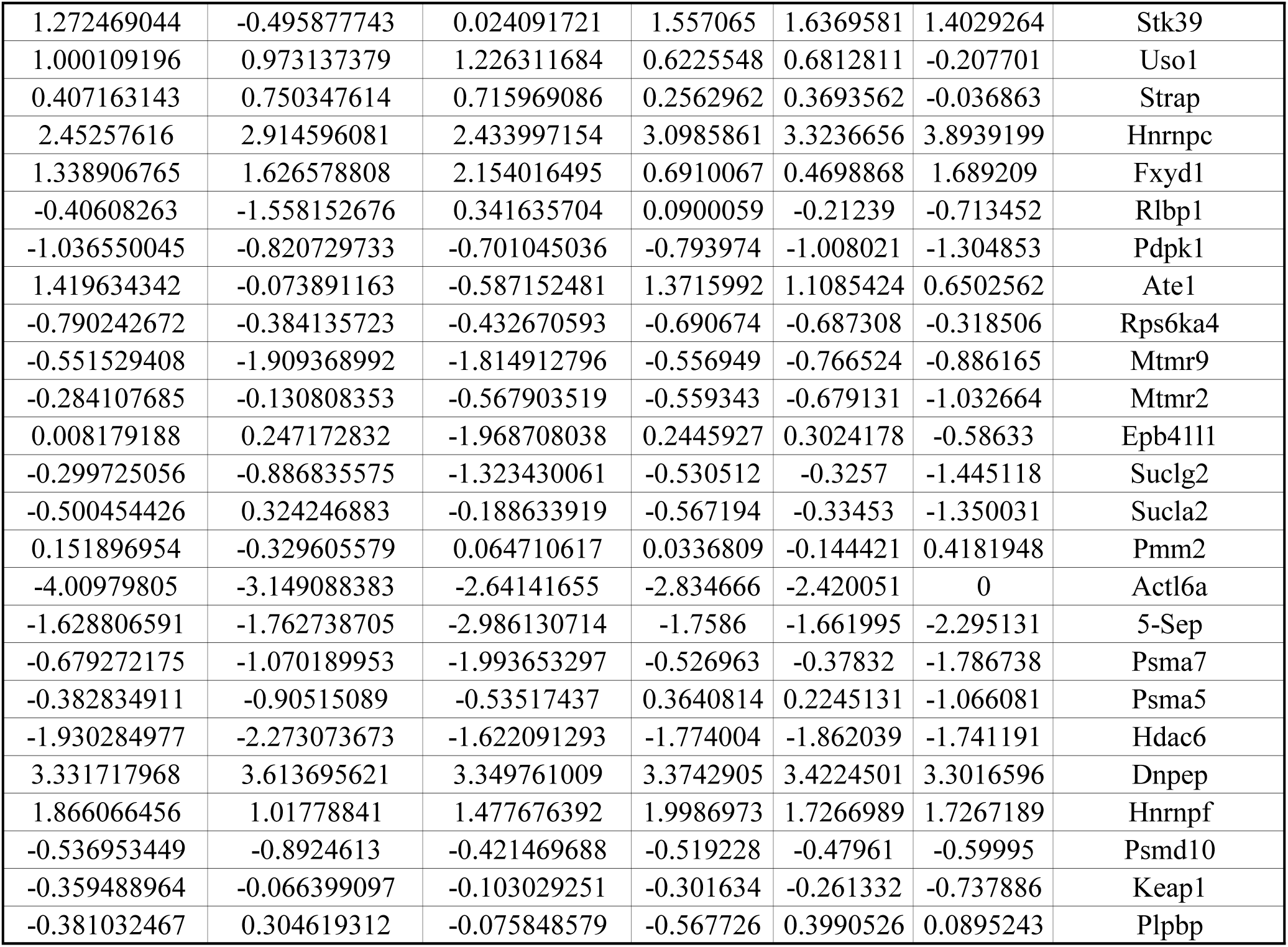
Proteonmics analysis of endosomes from sleep deprivation followed recovery and sleep deprivation only mice.

**Extended Data Table 2.**
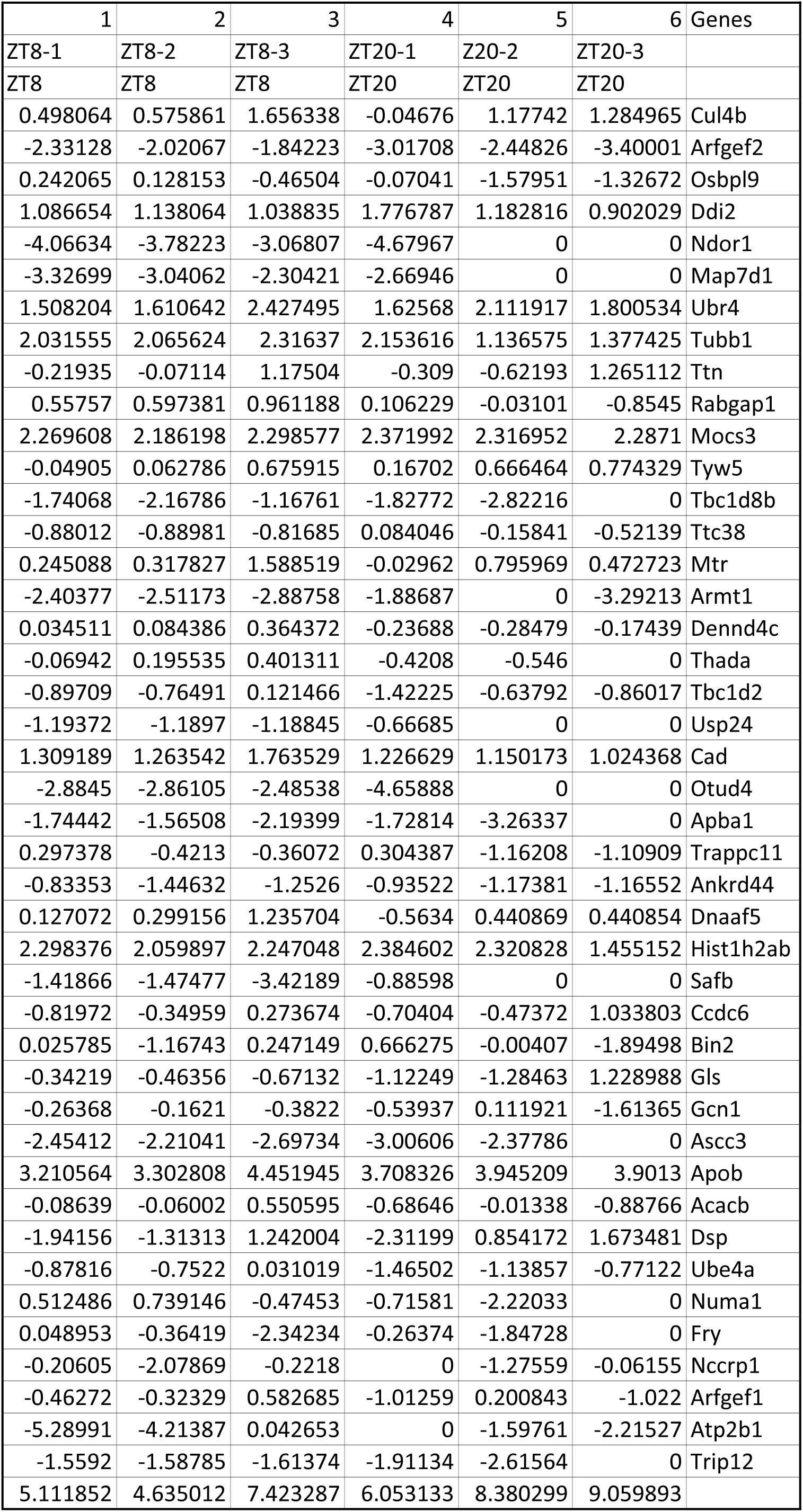
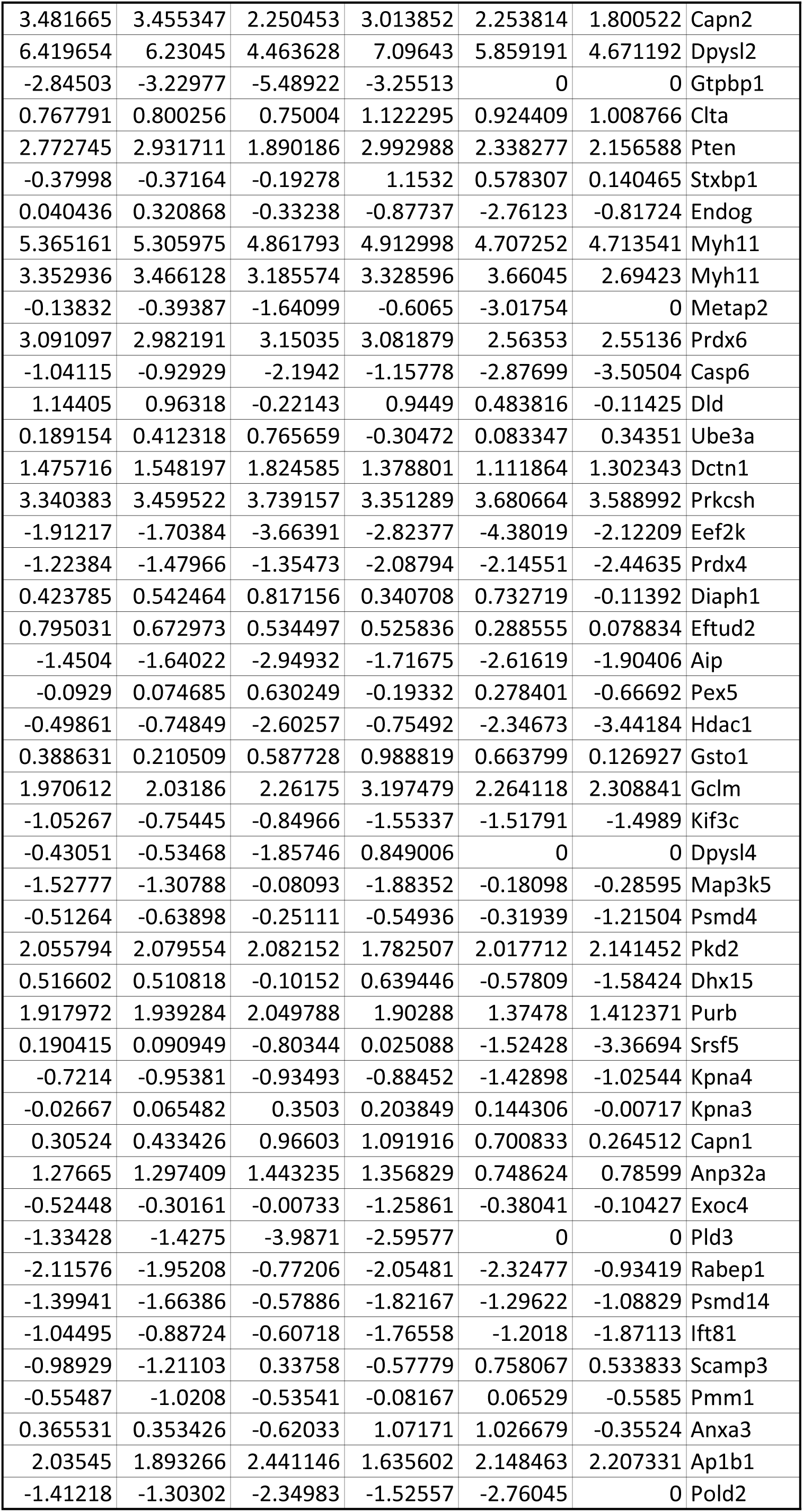
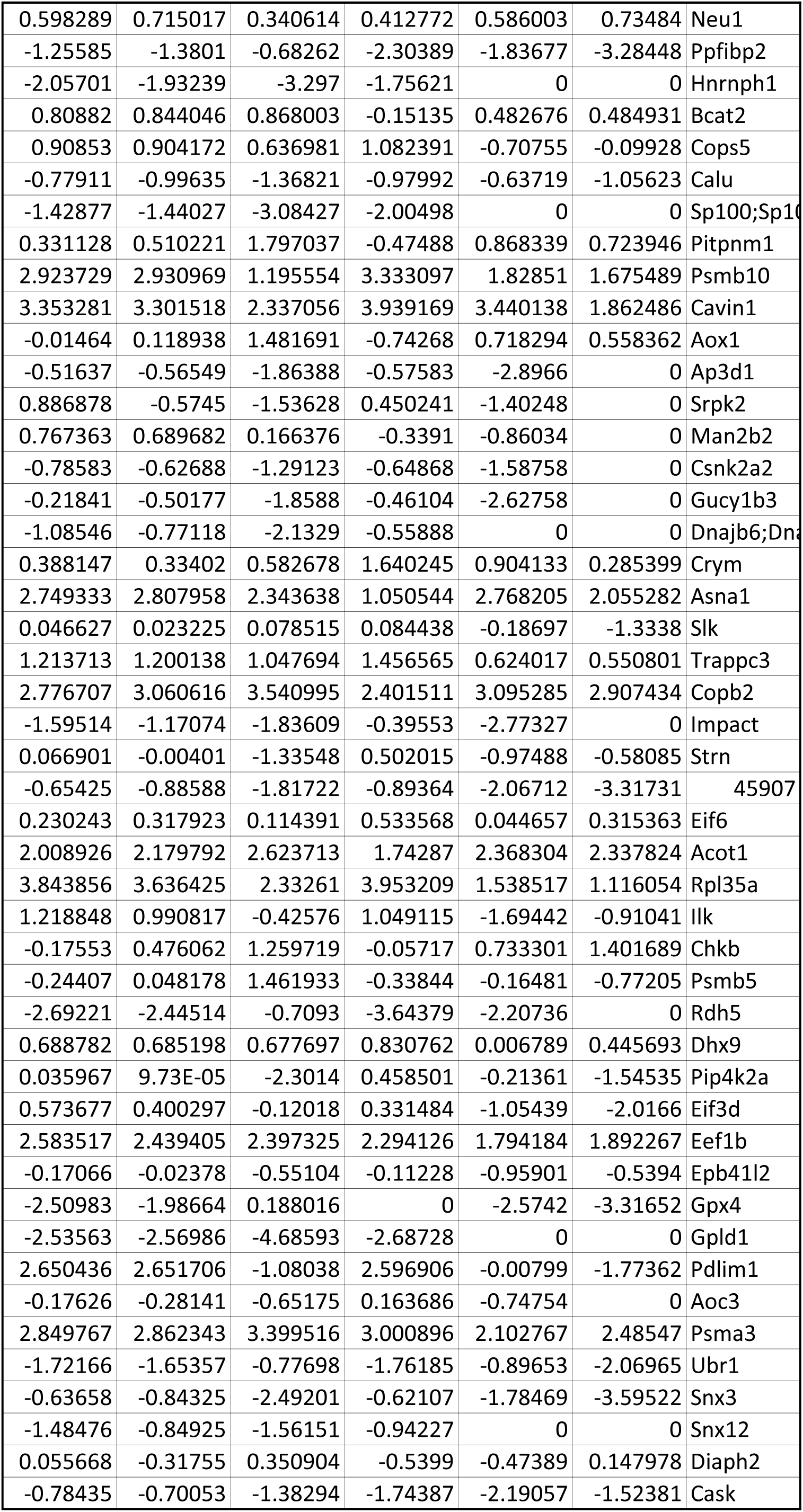
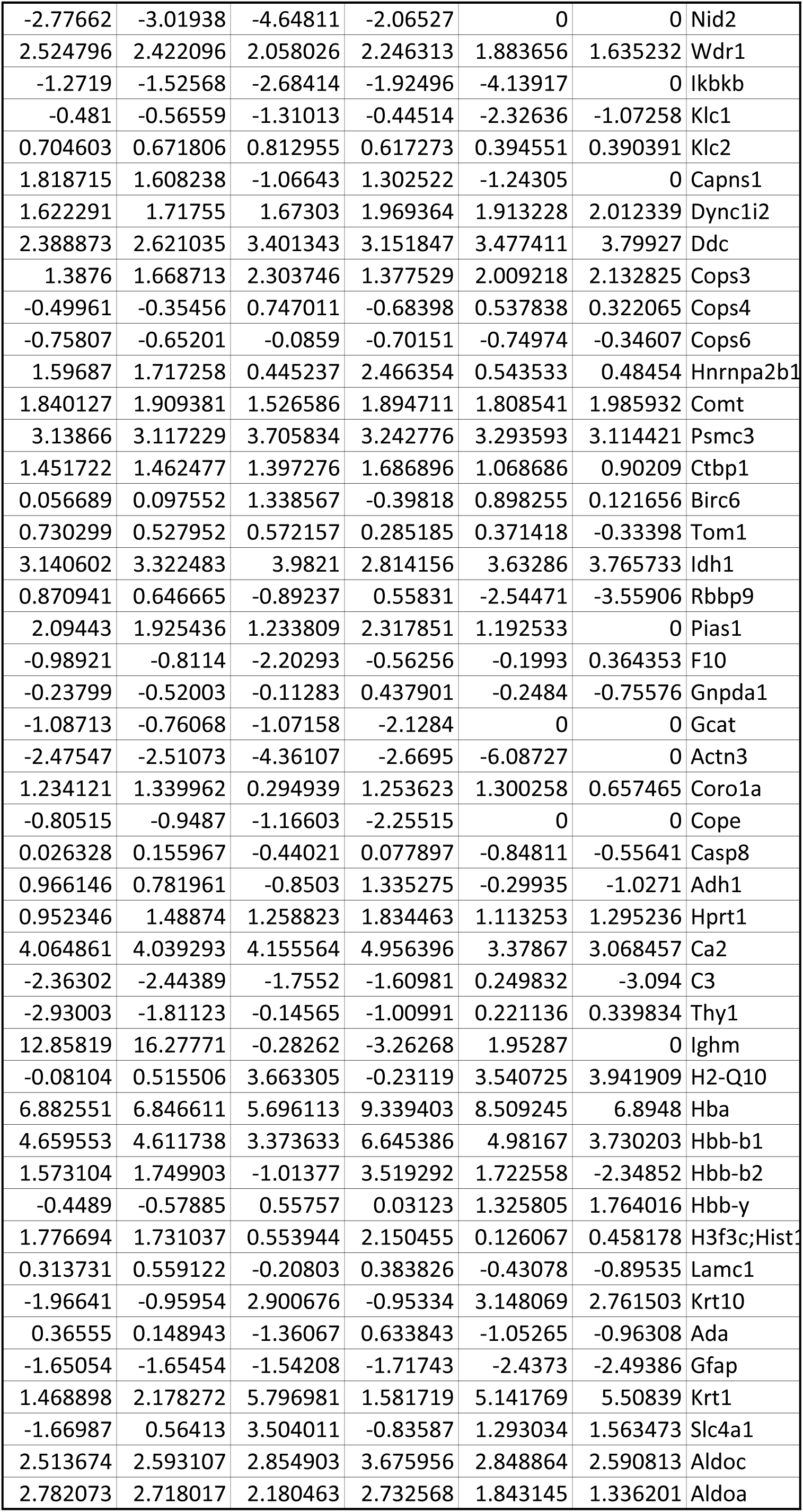
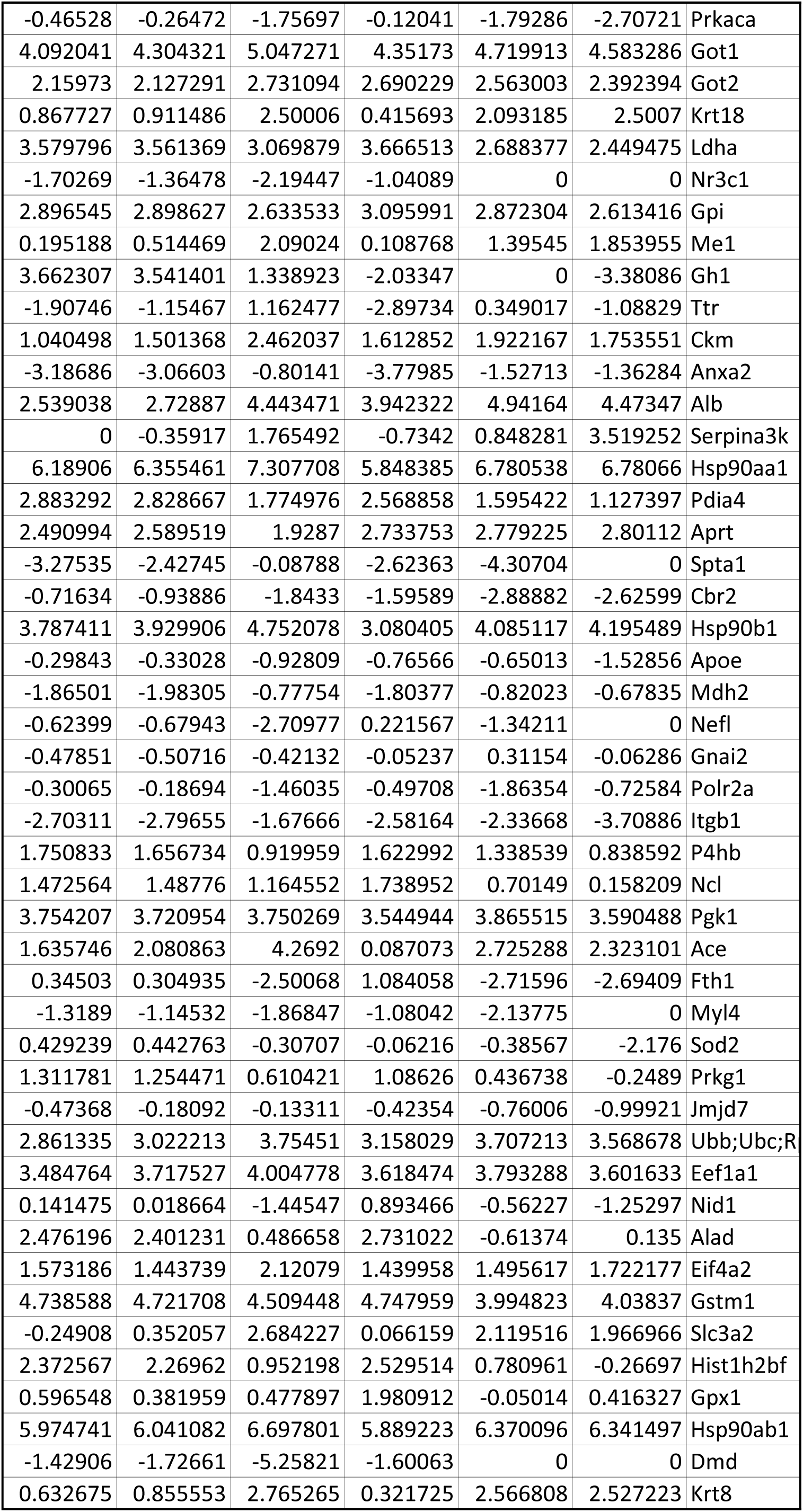
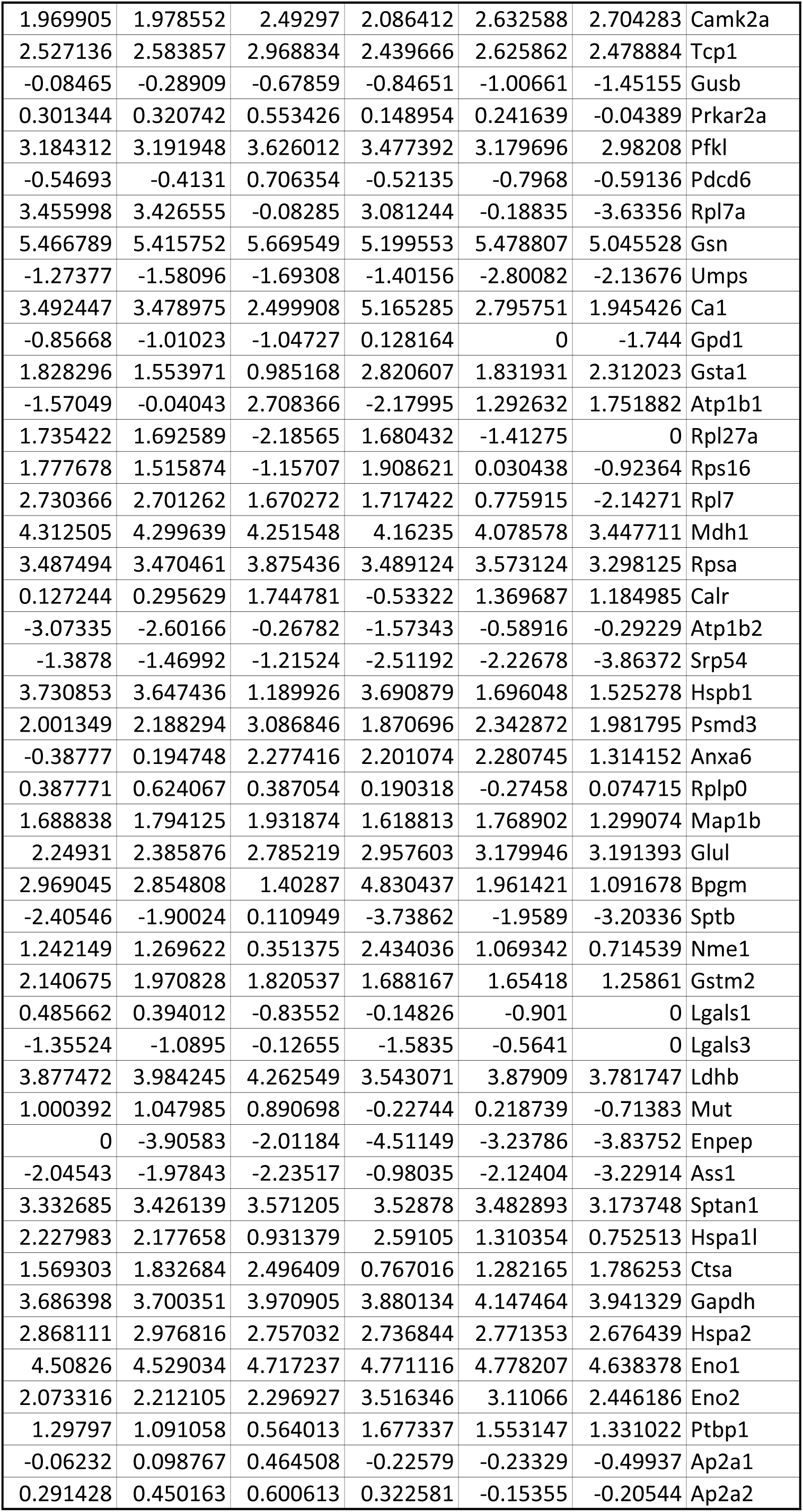
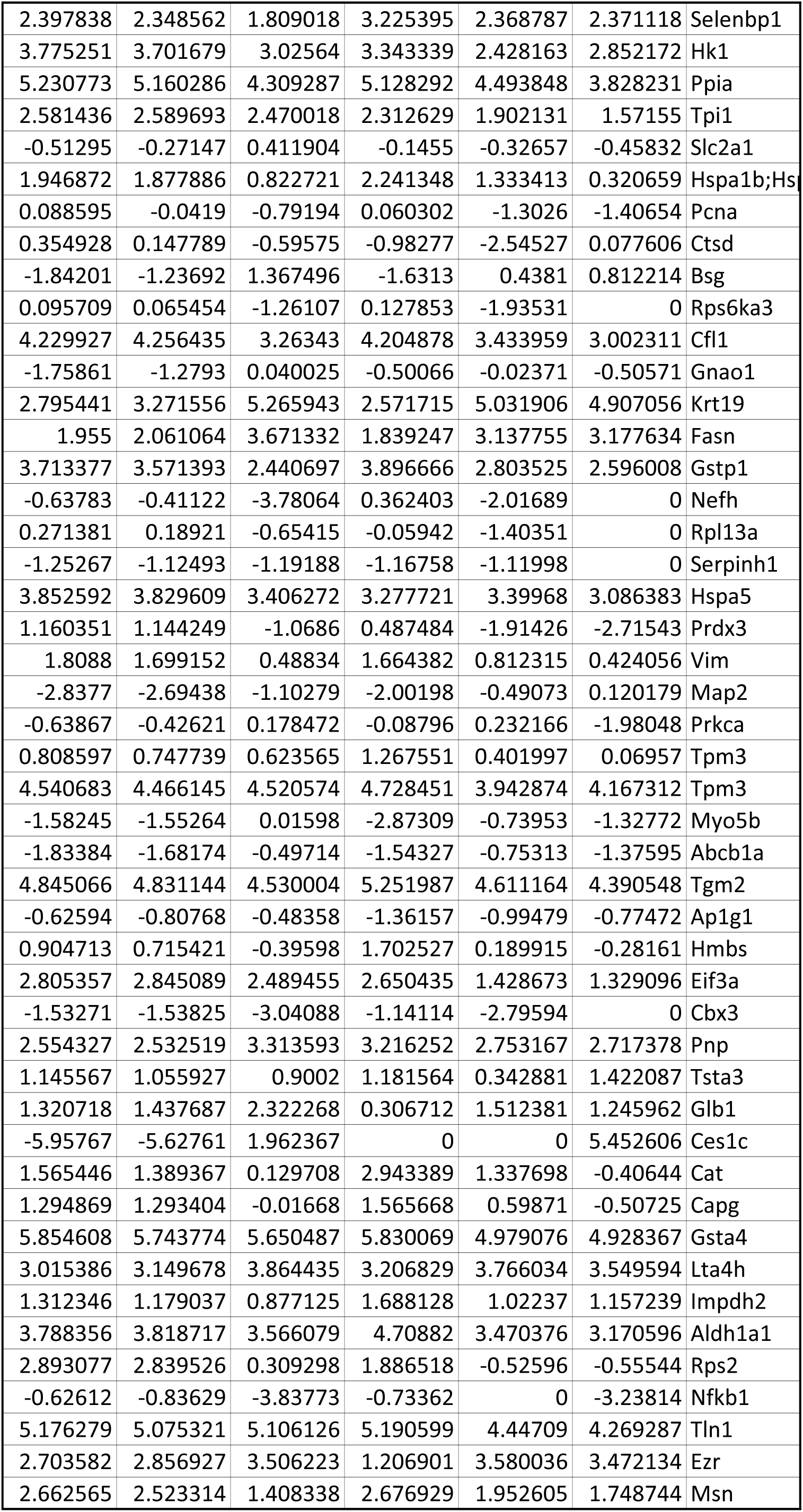
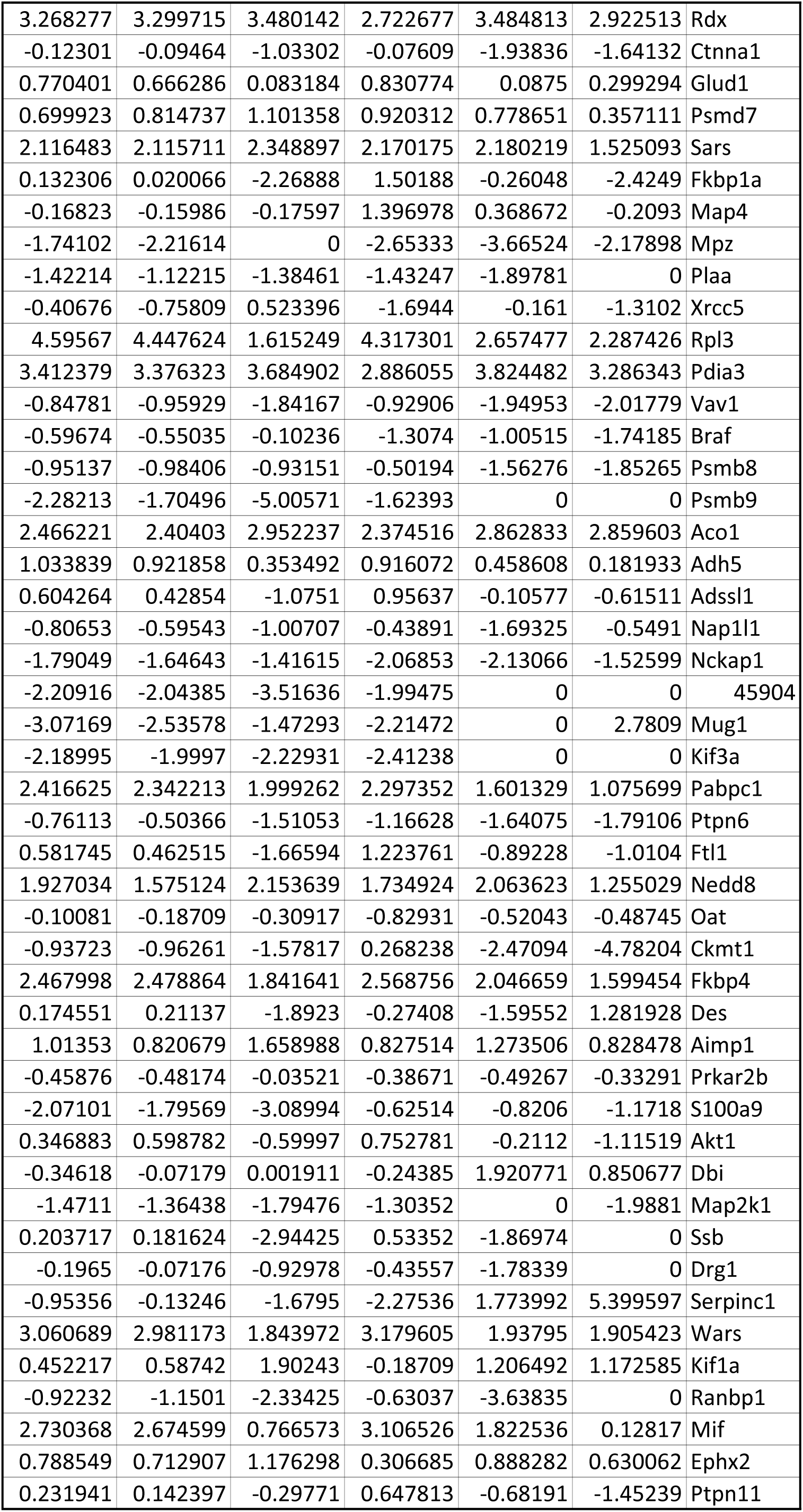
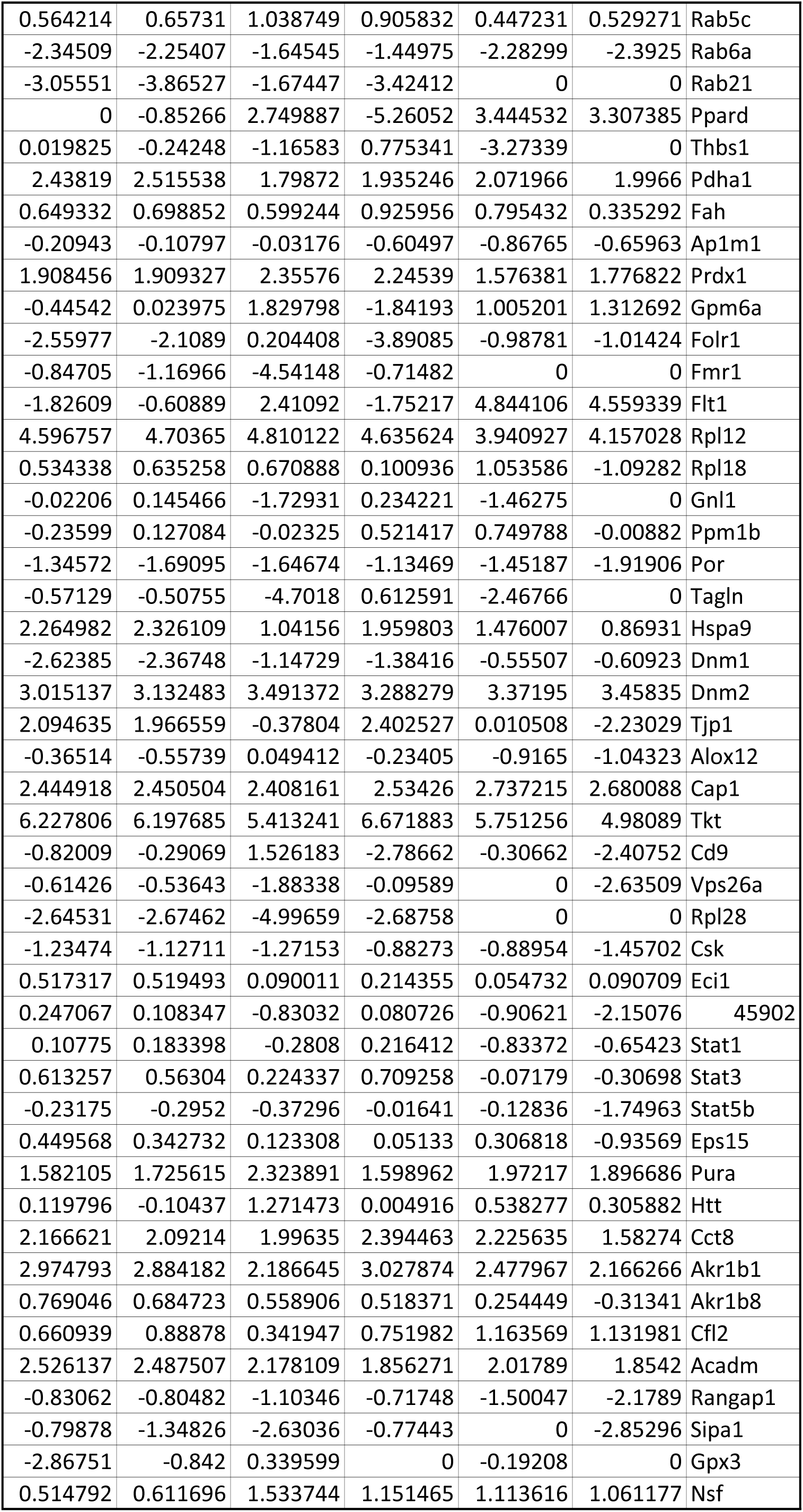
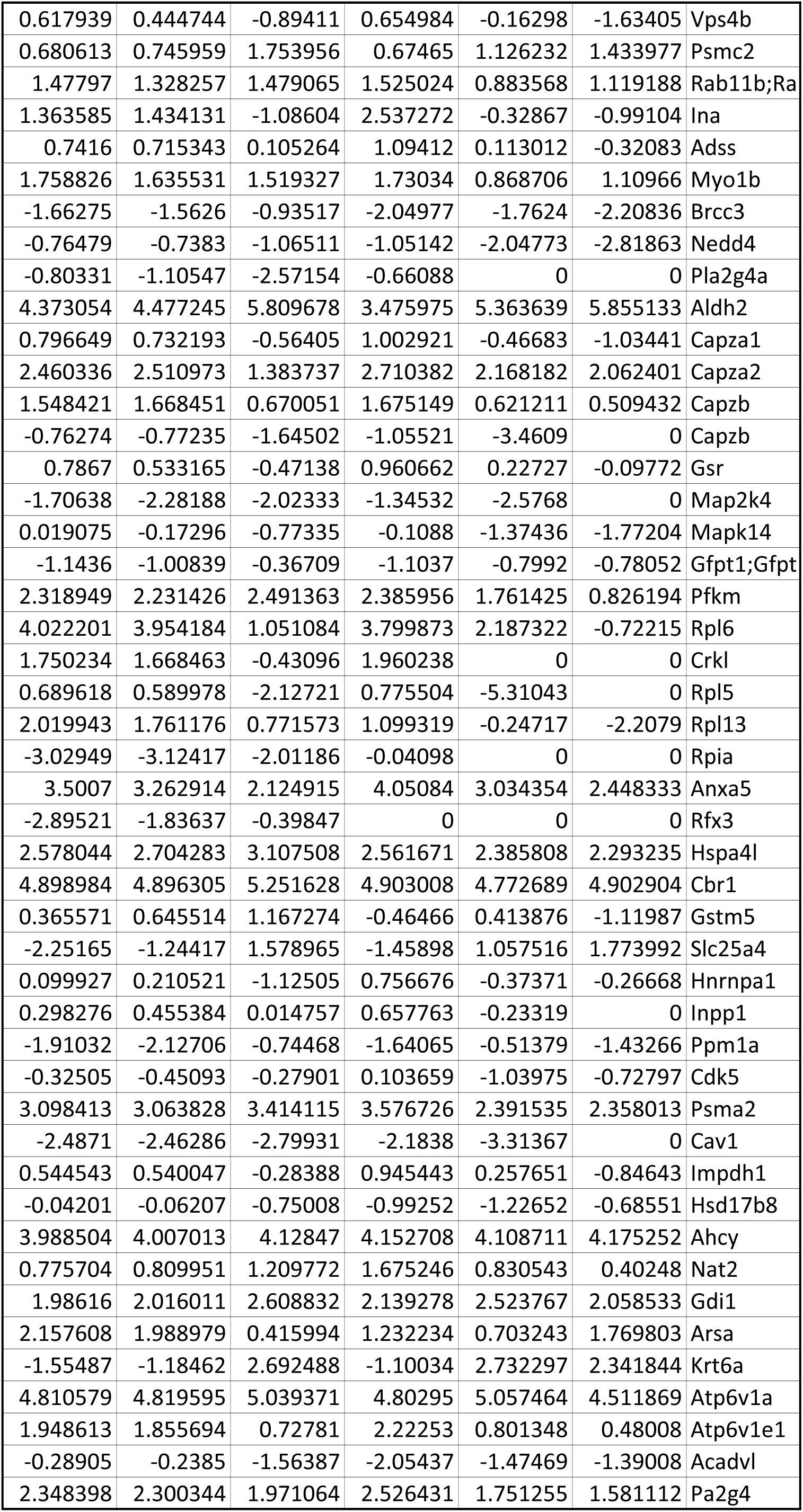
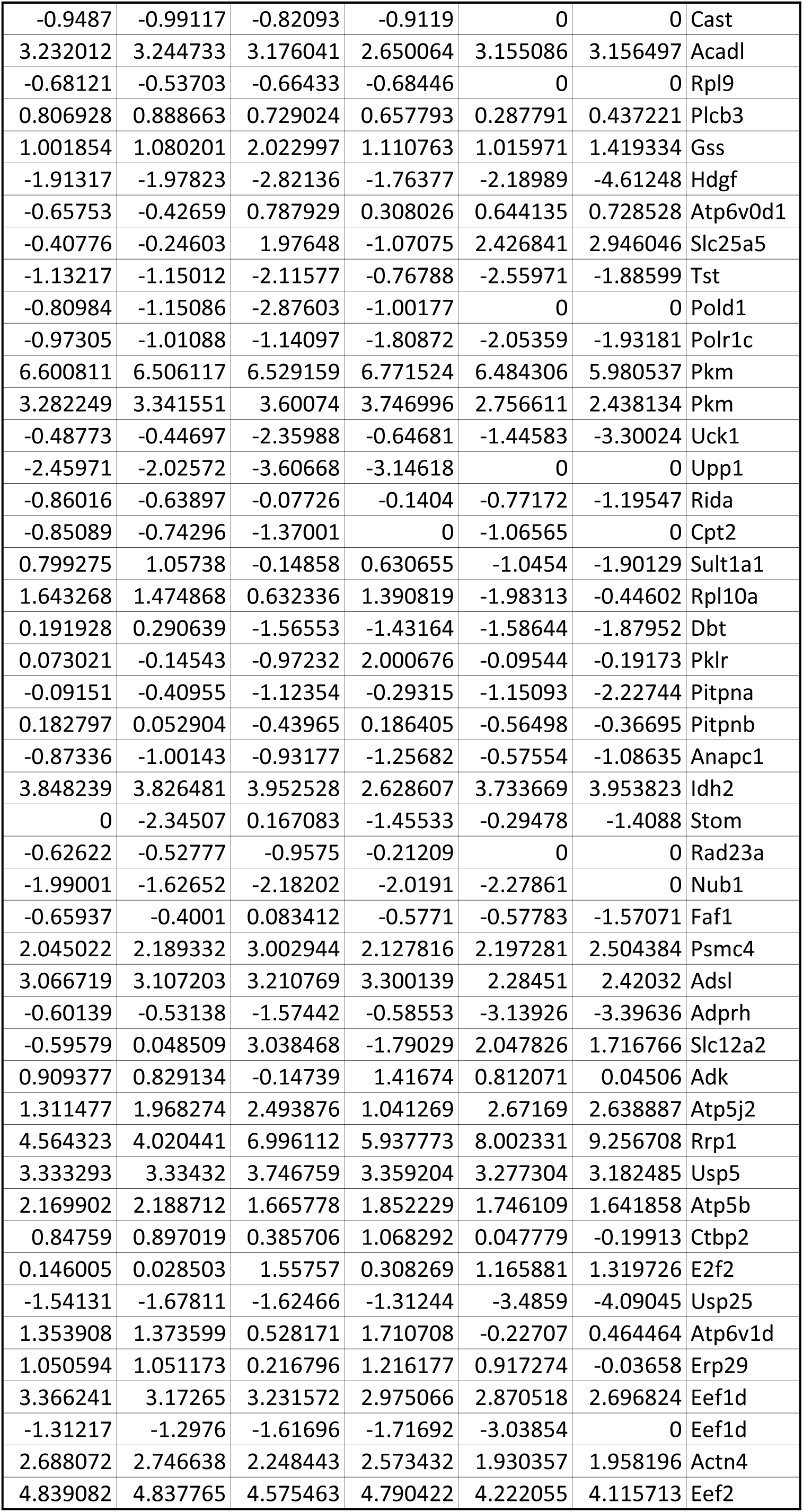
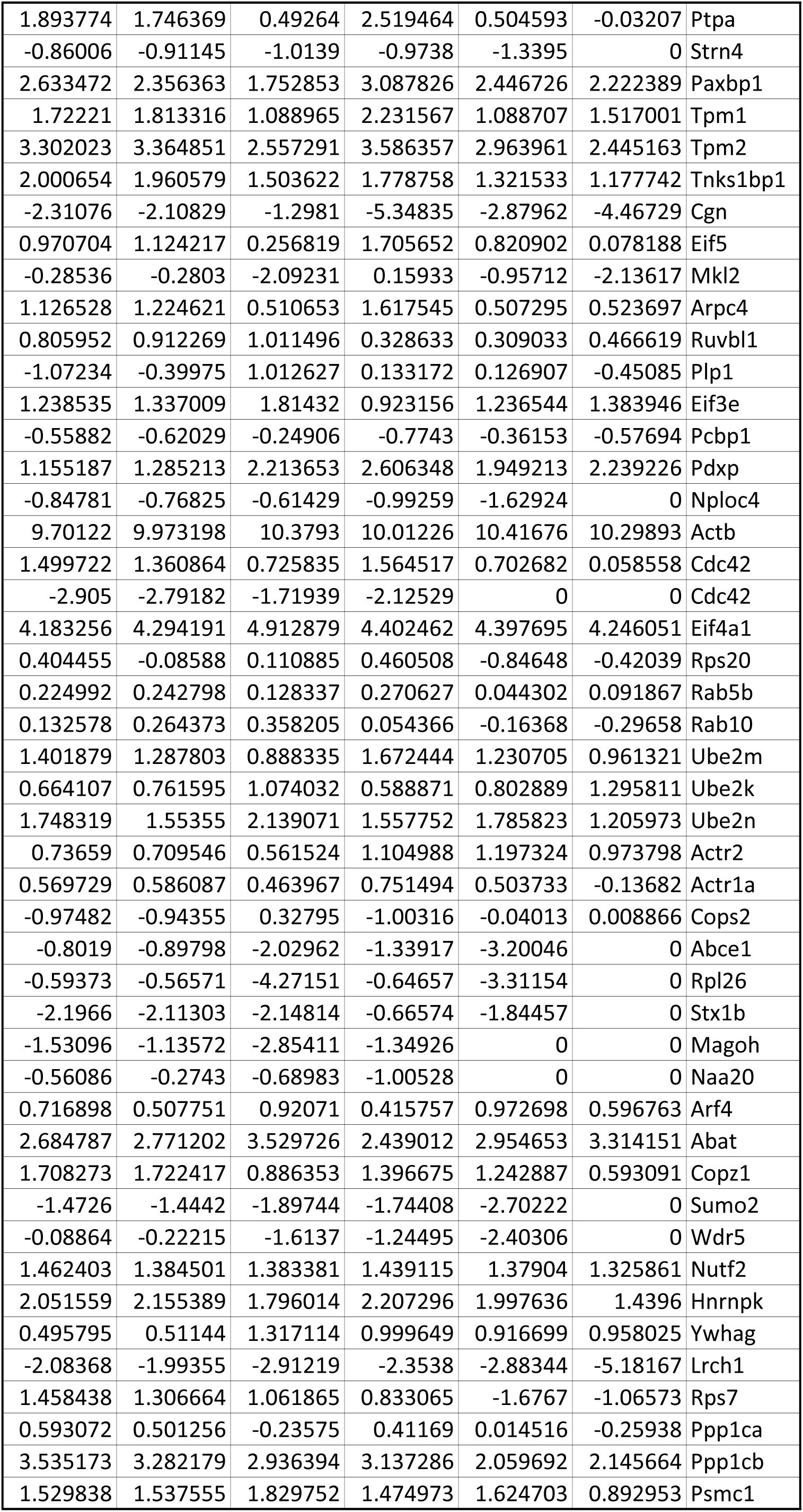
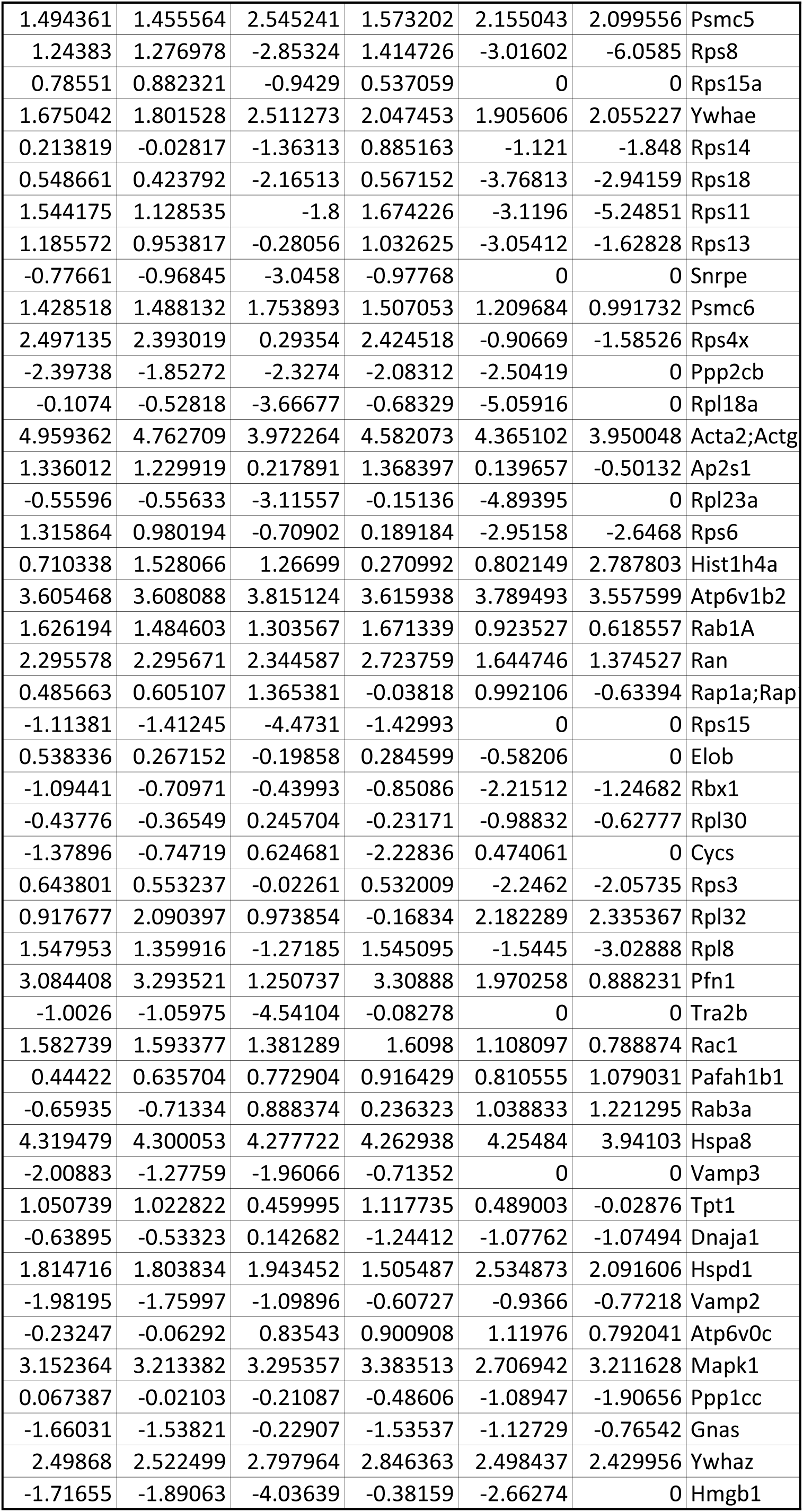
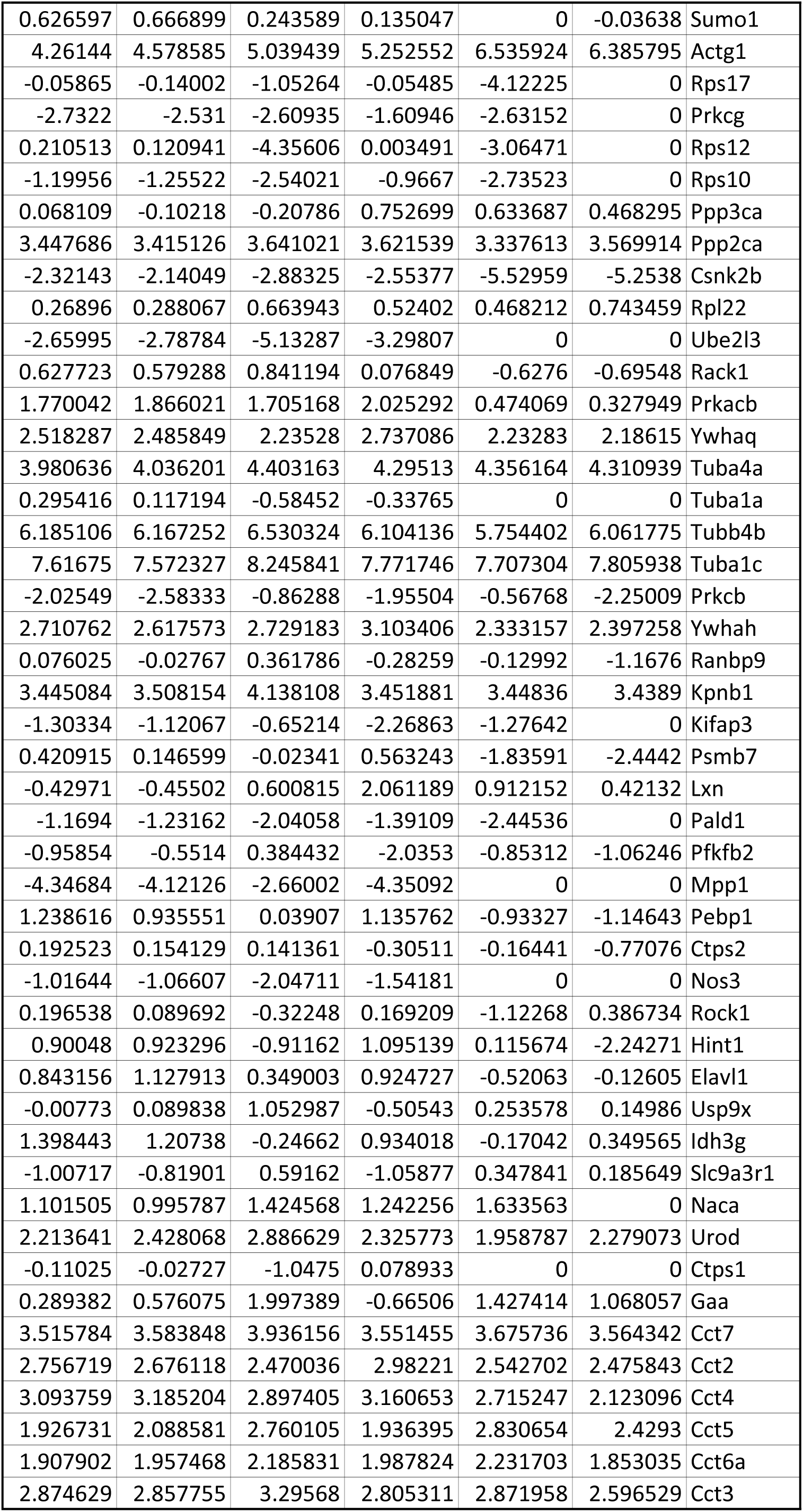
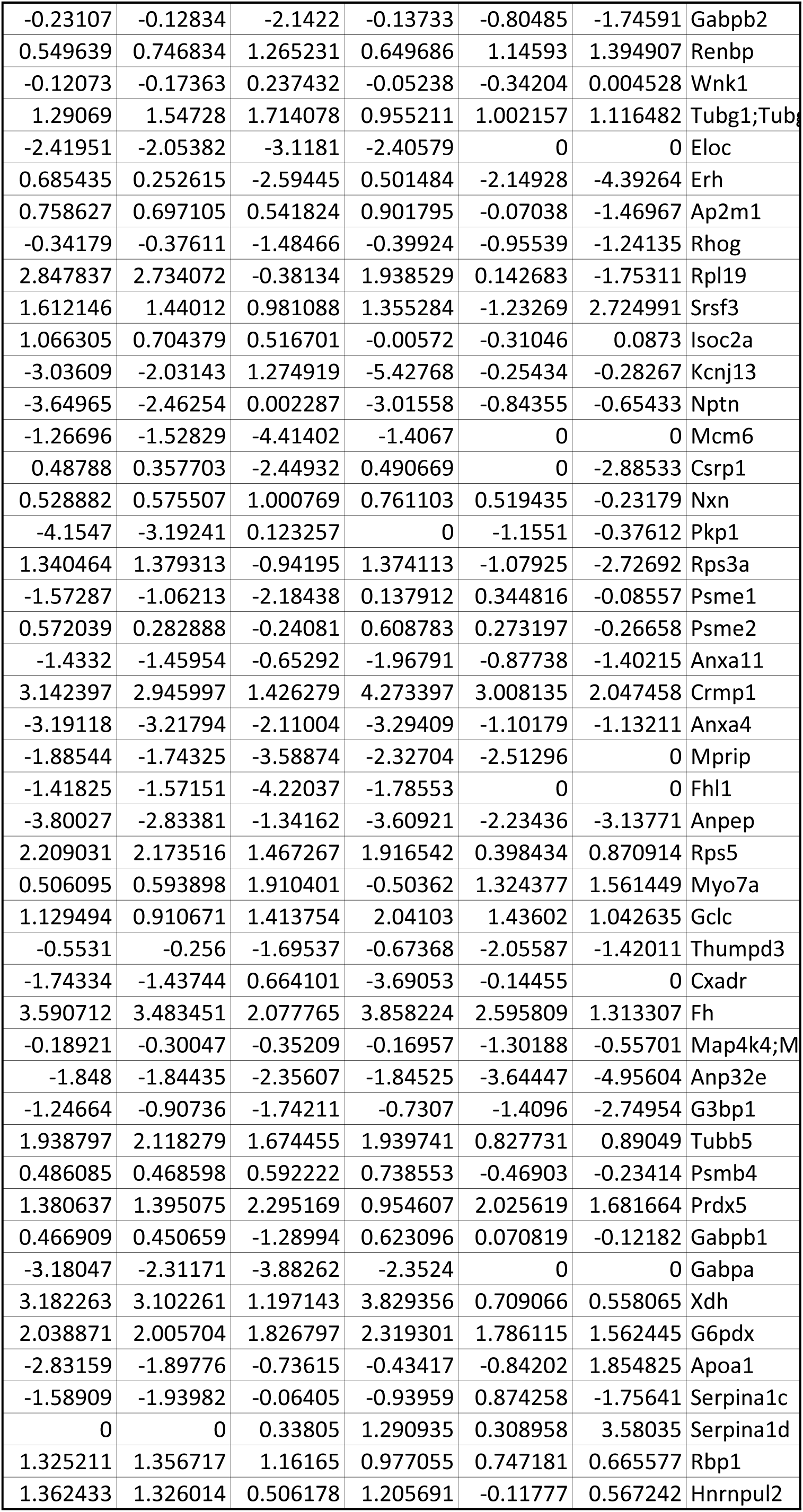
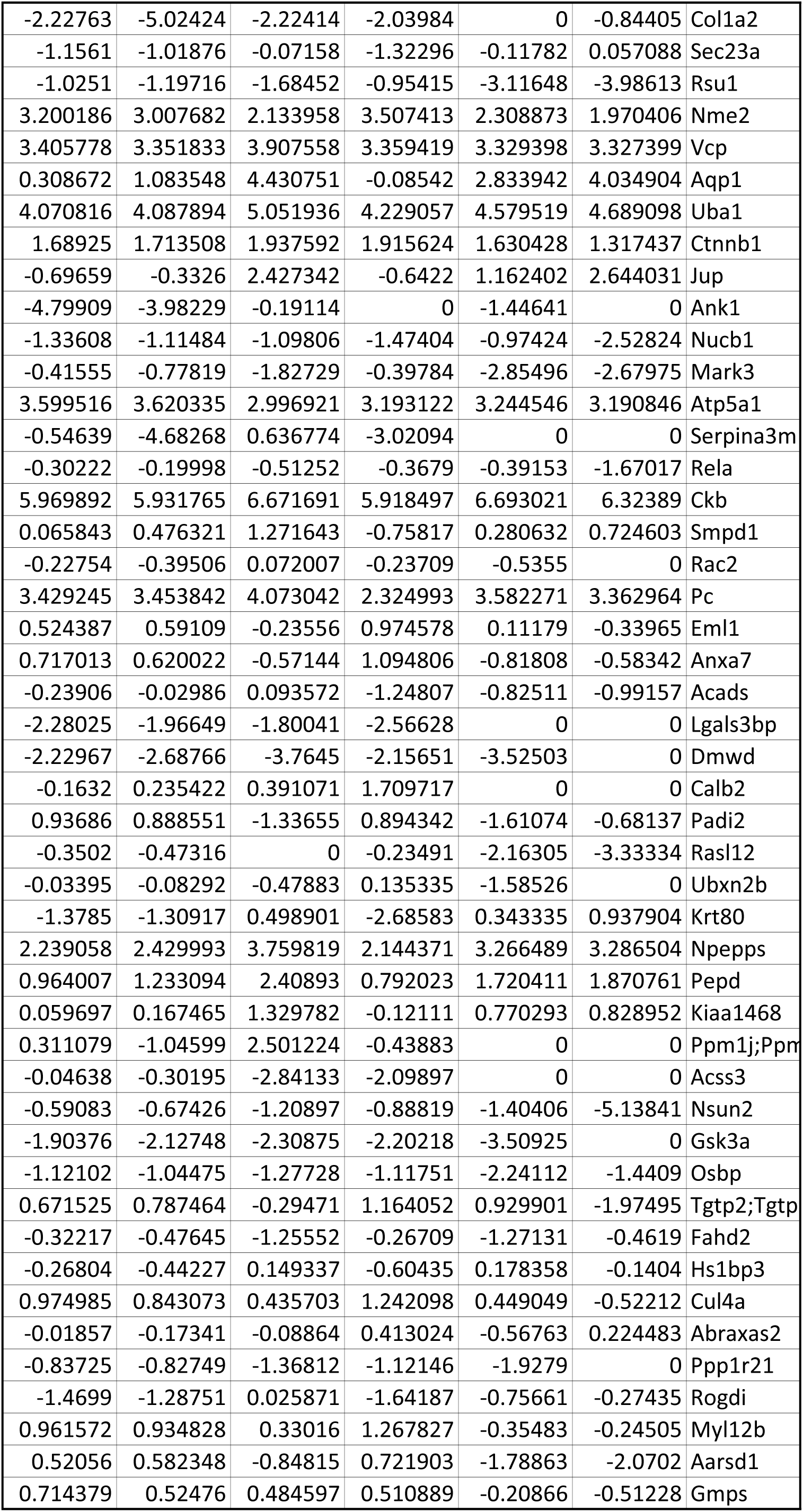
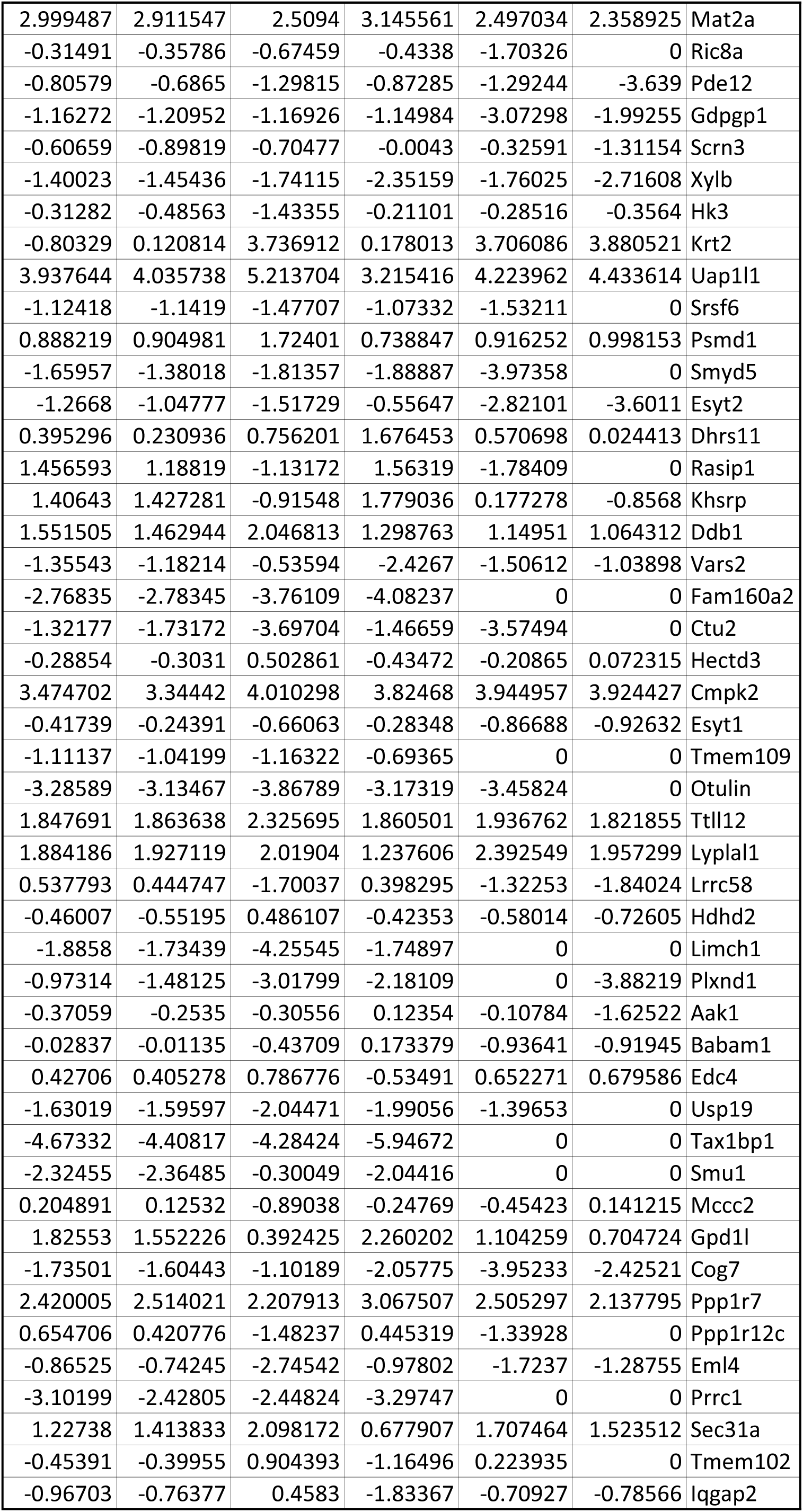
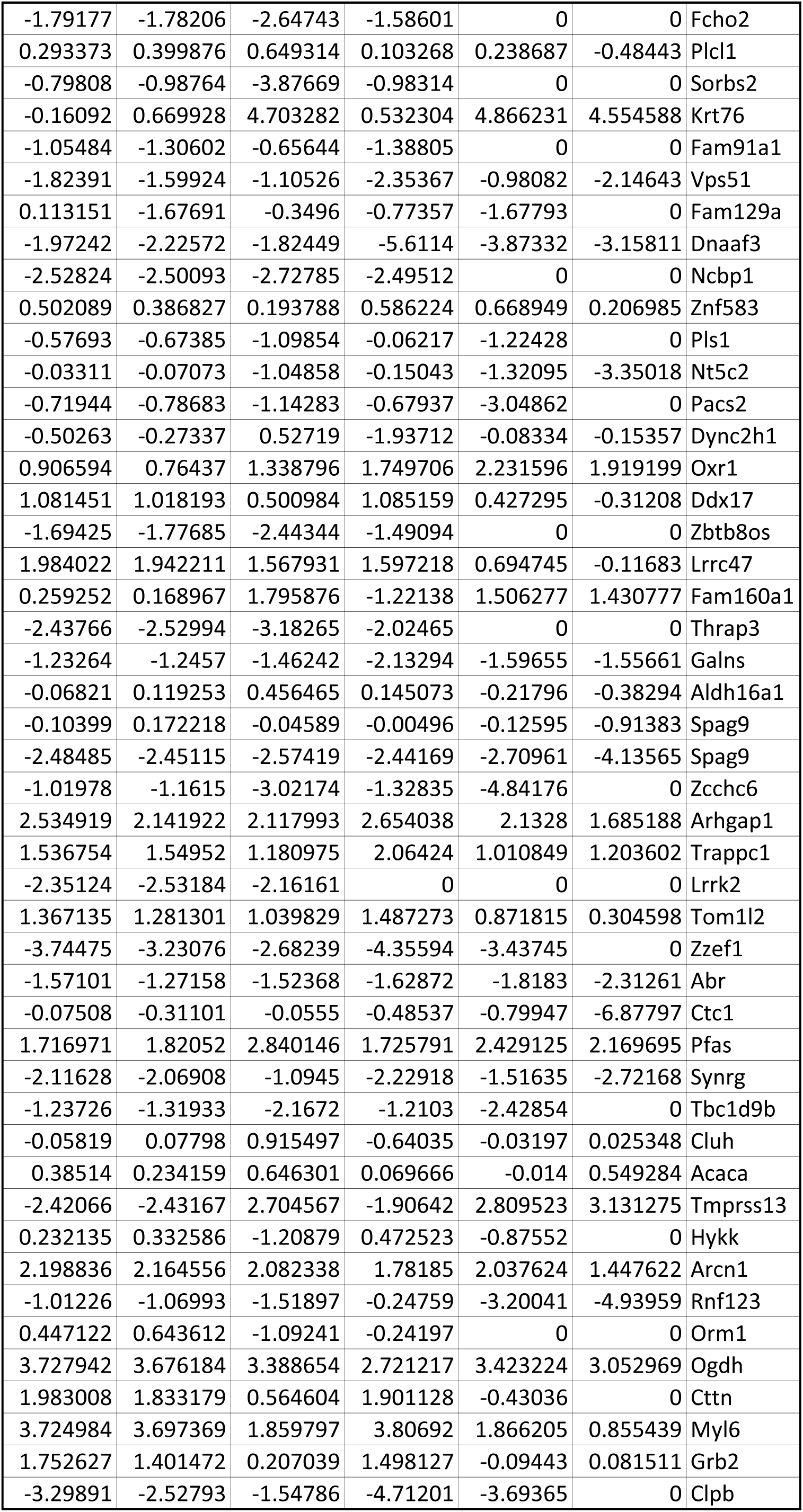
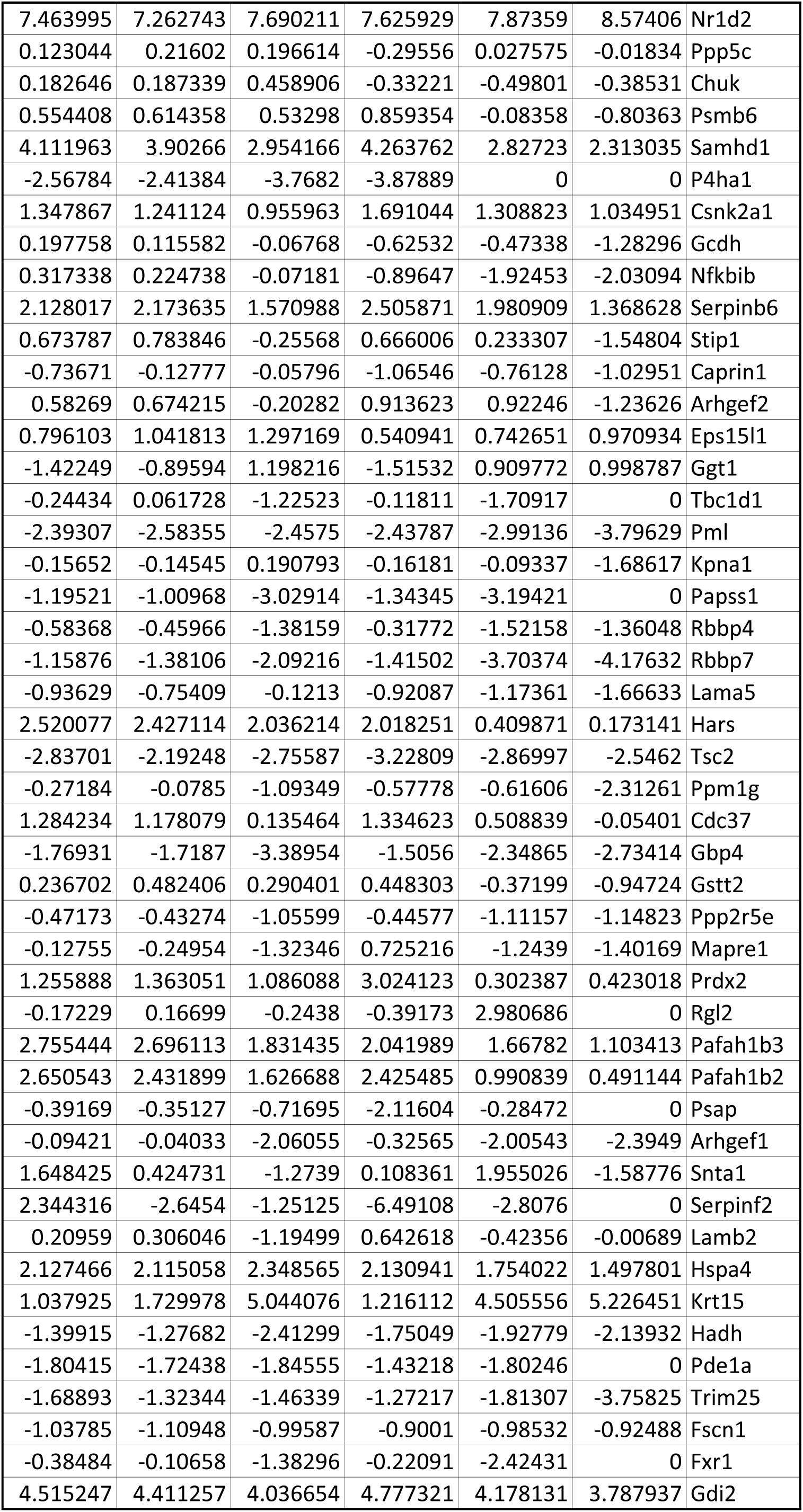
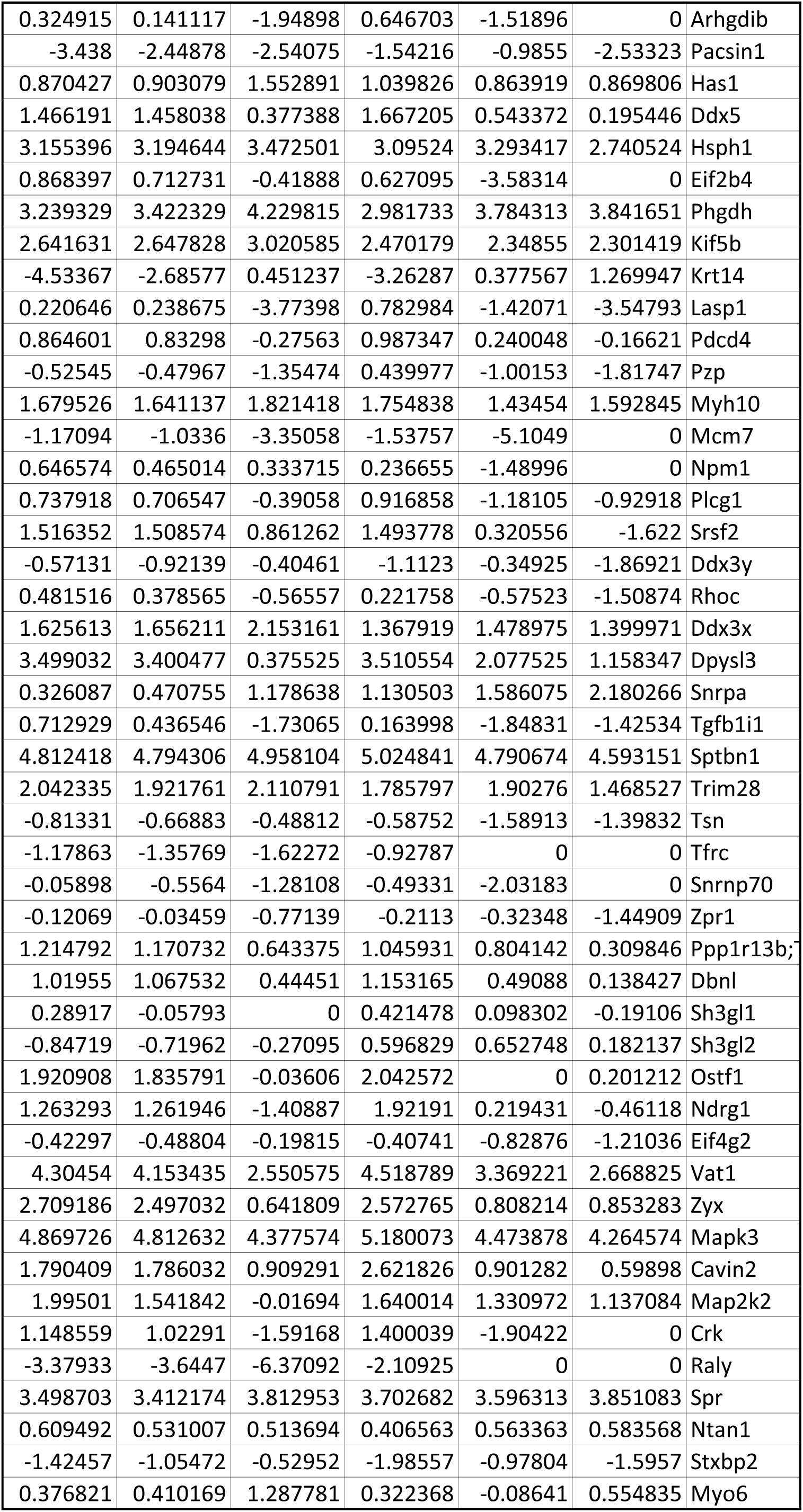
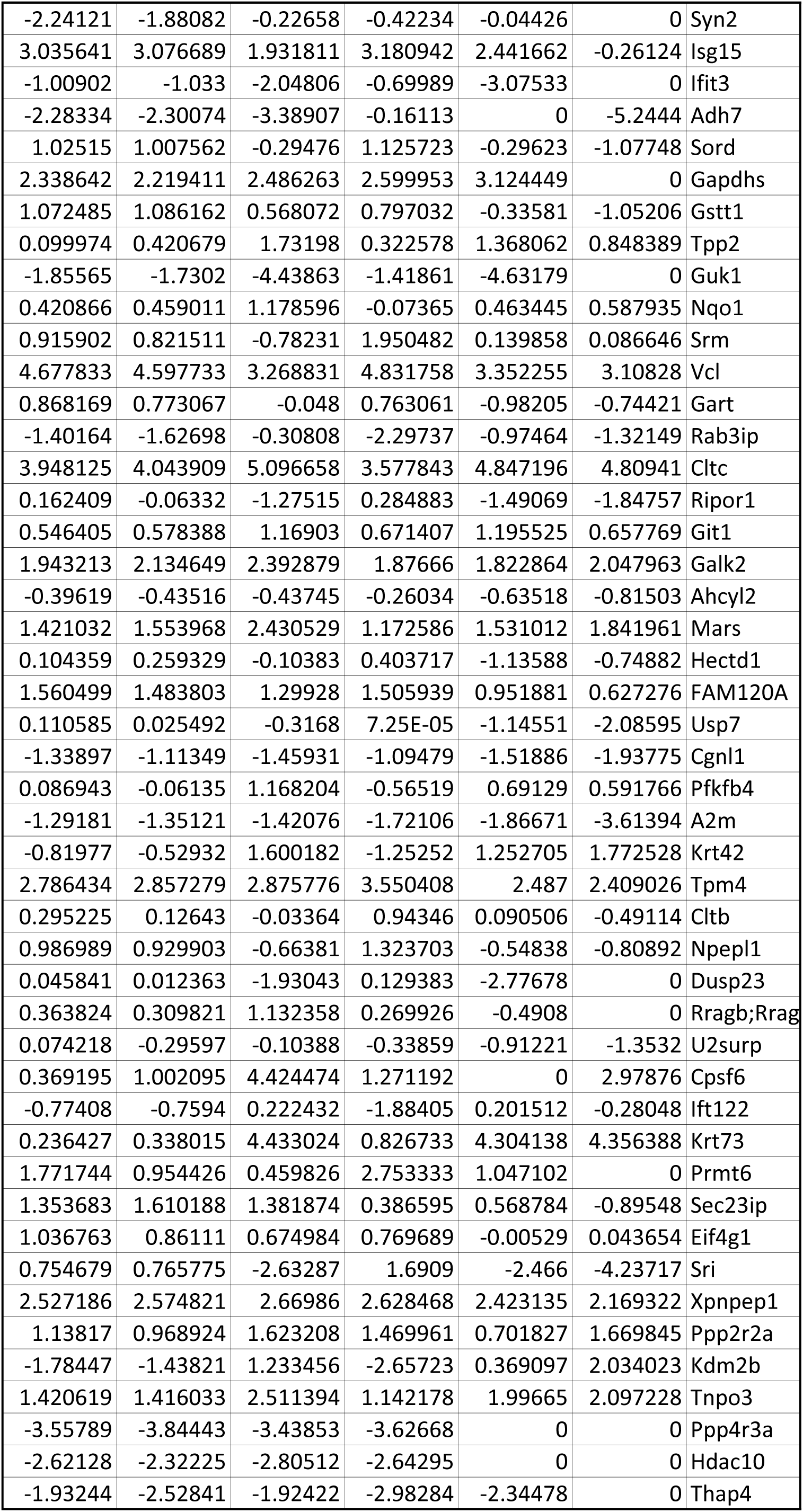
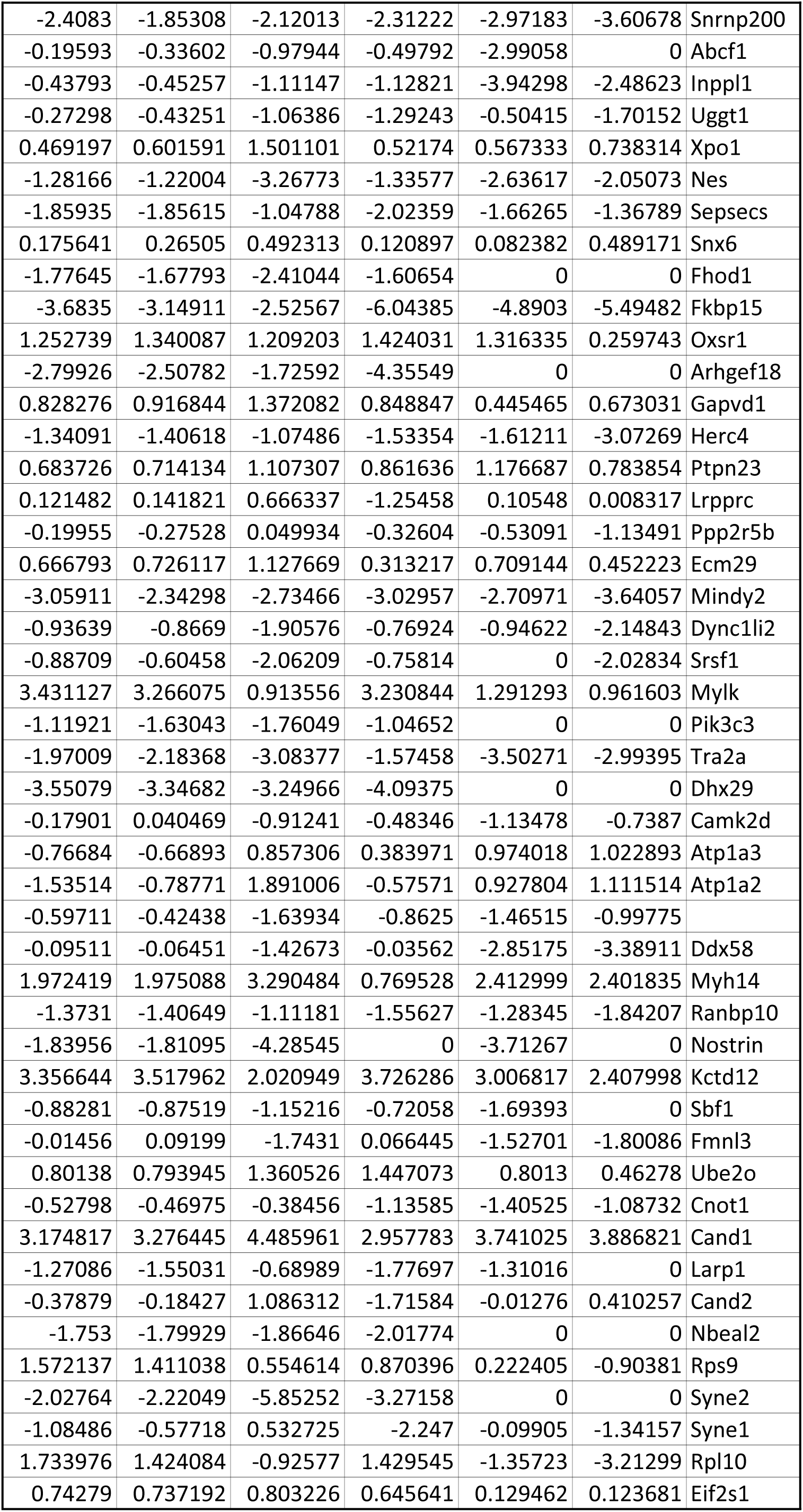
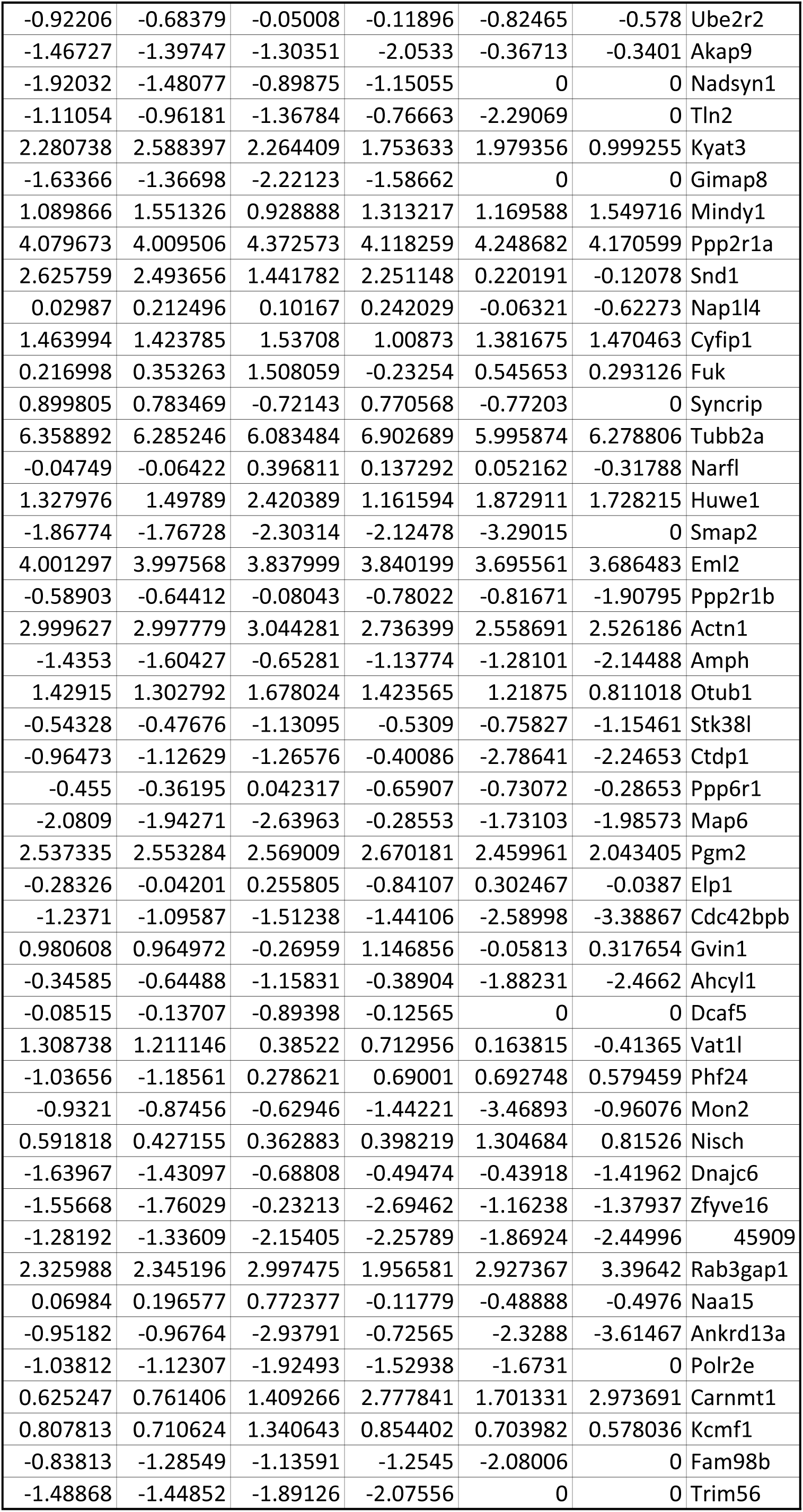
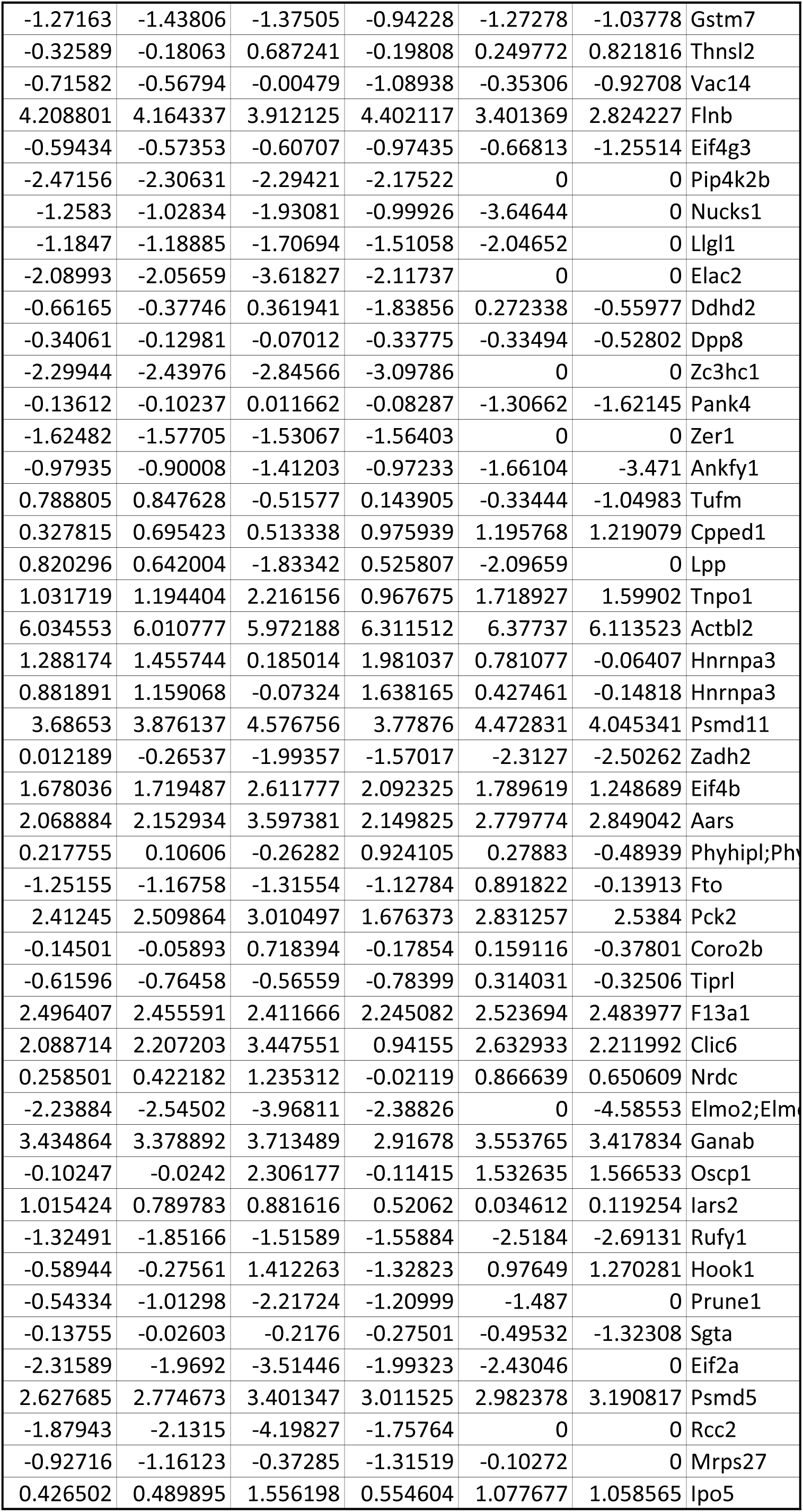
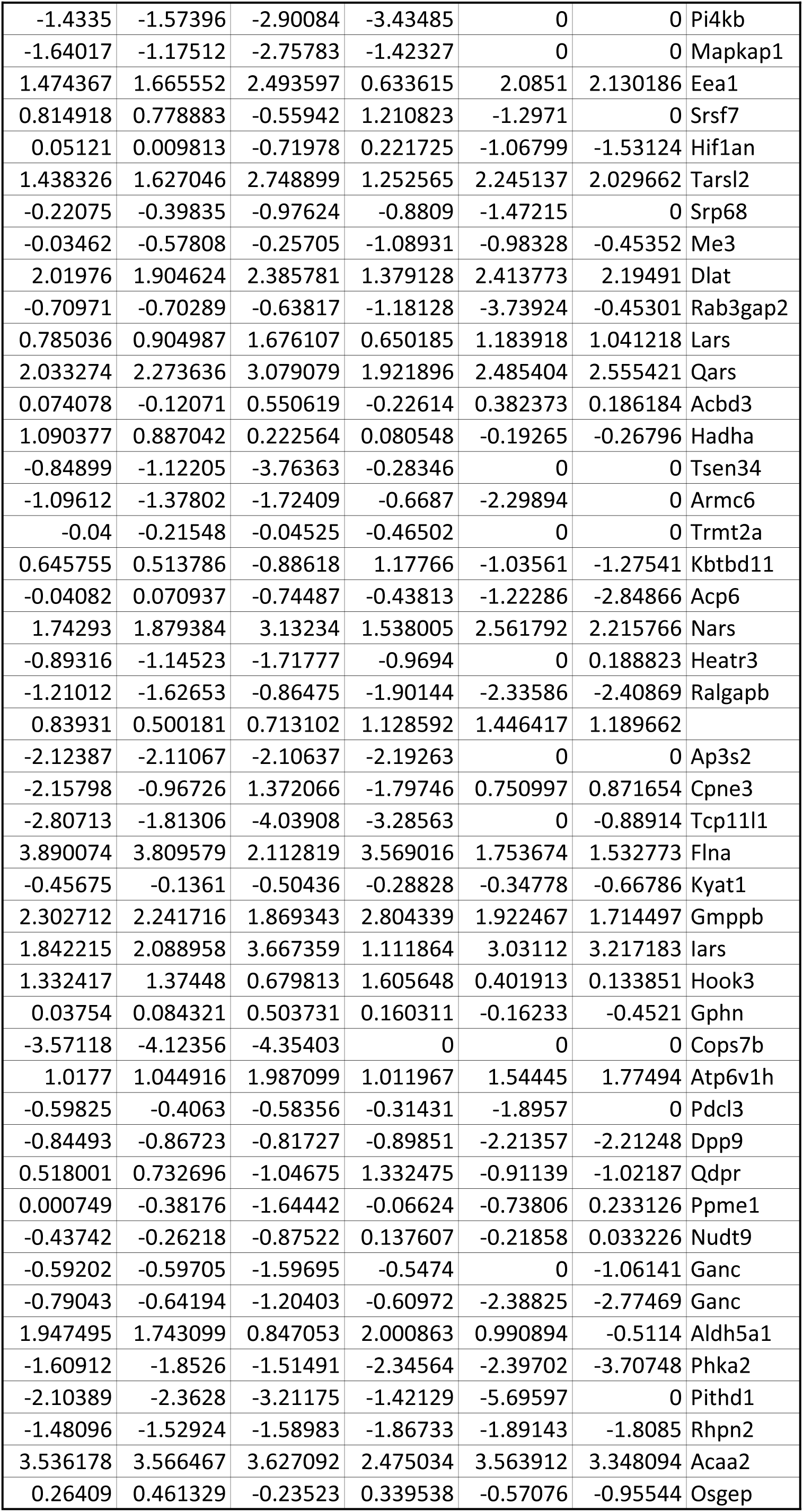
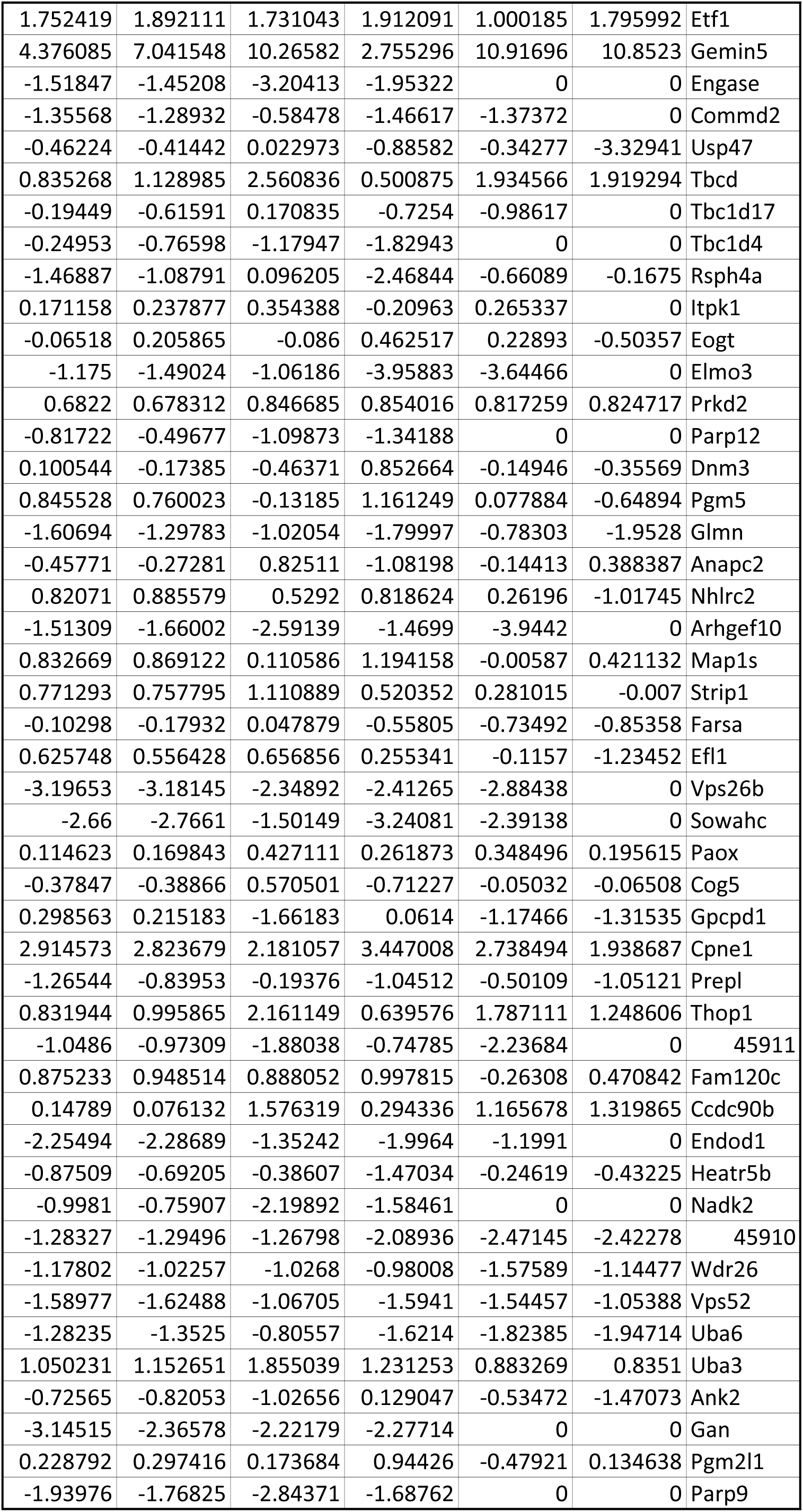
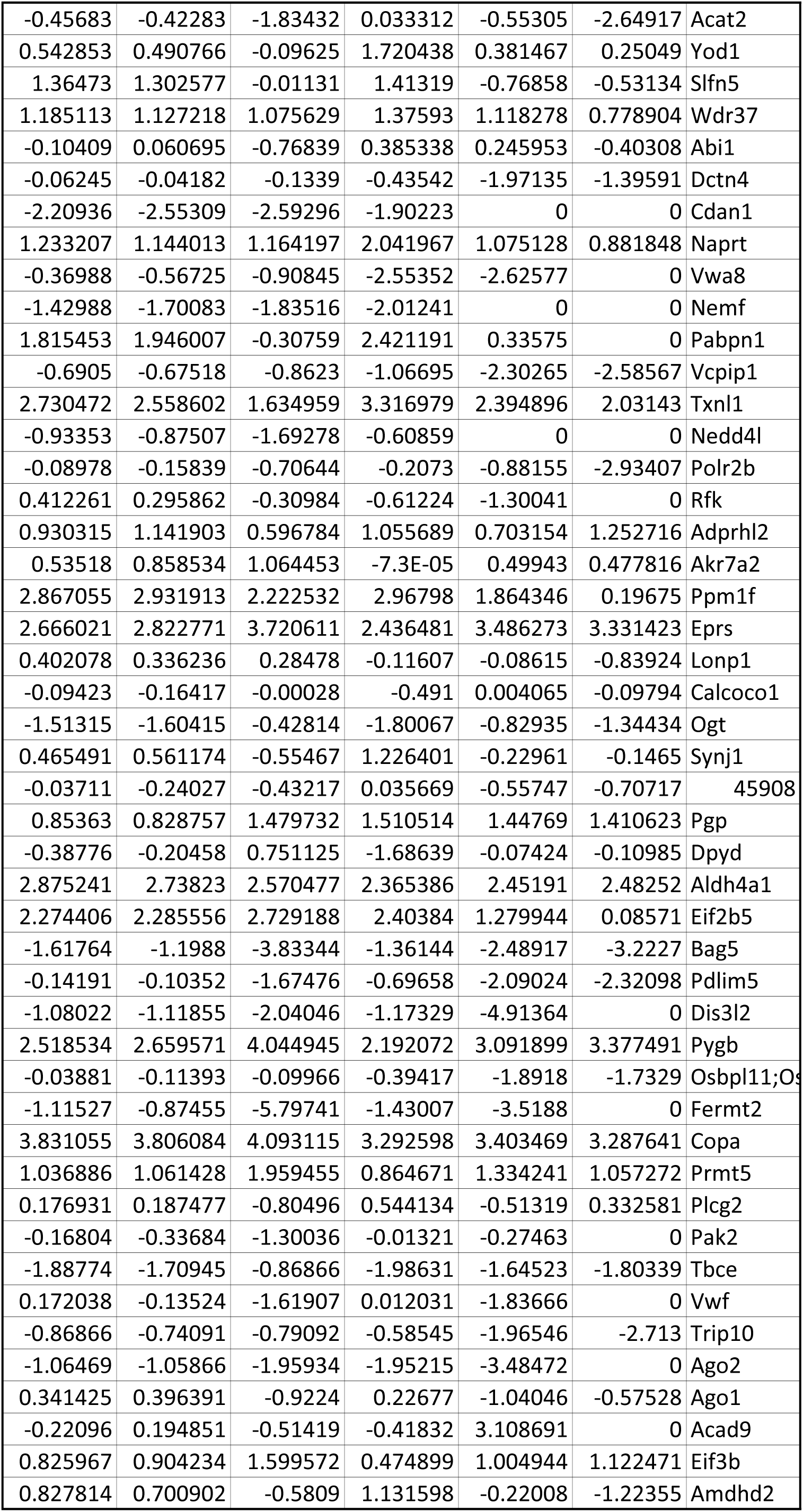
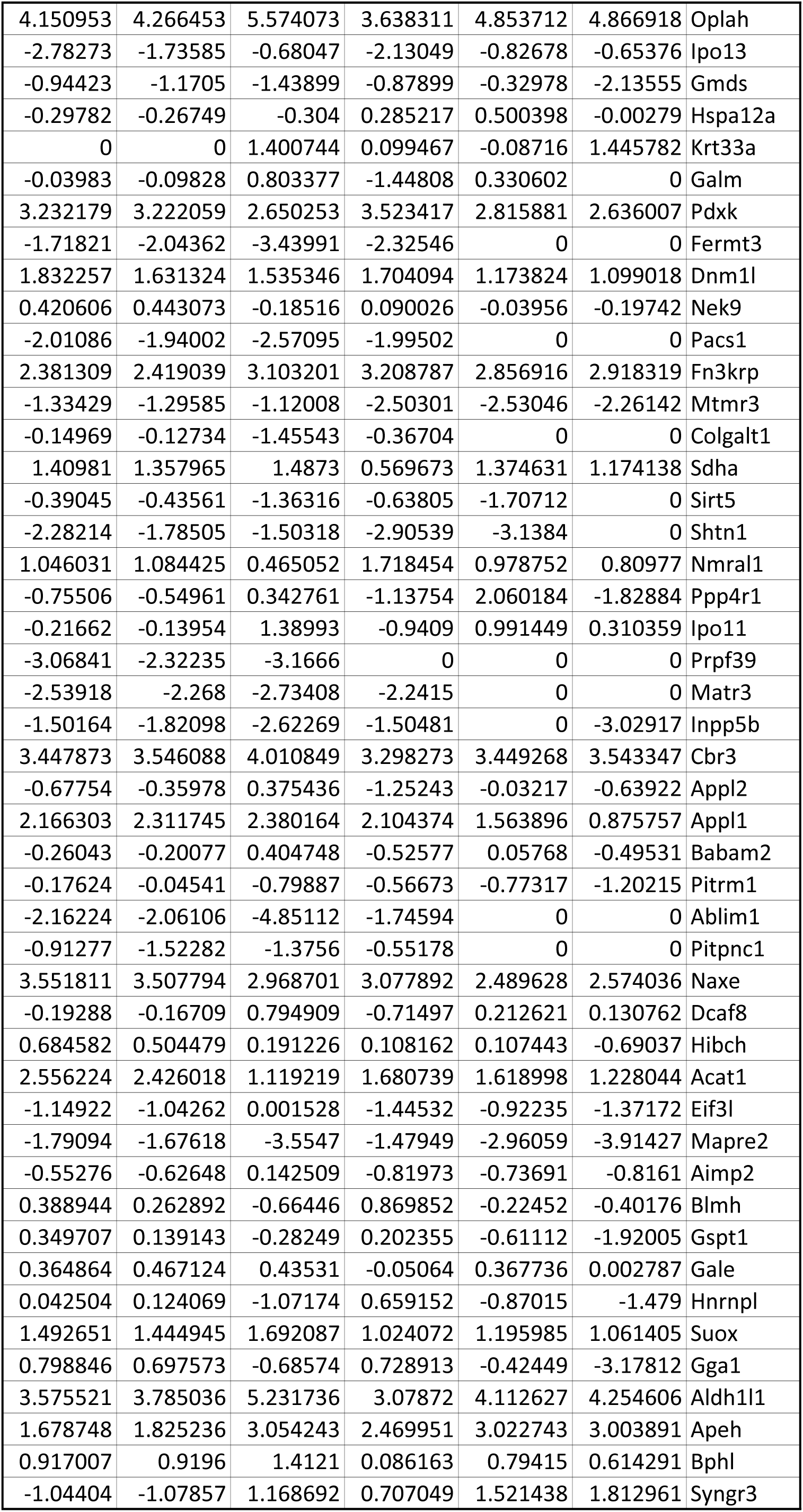
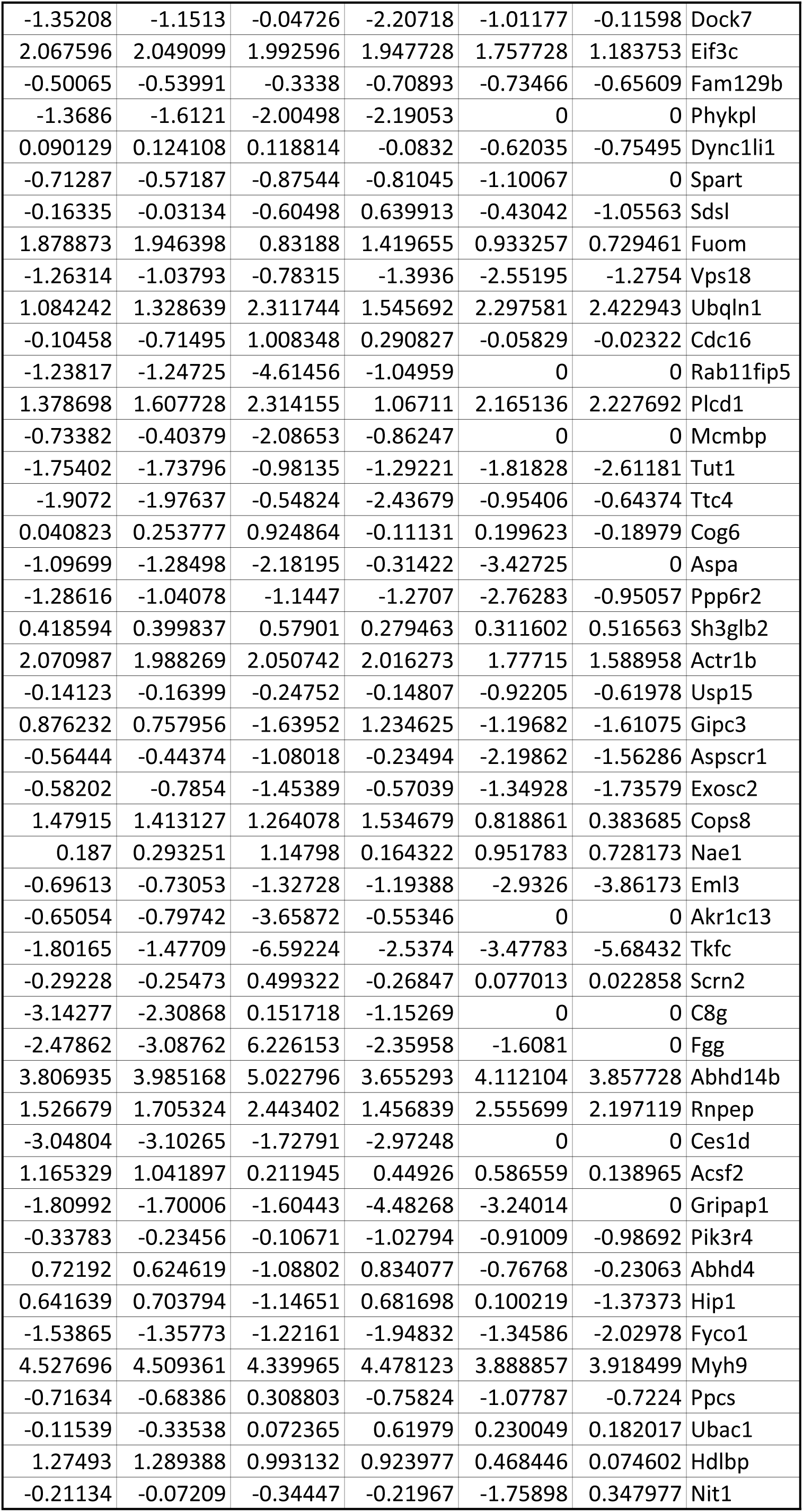
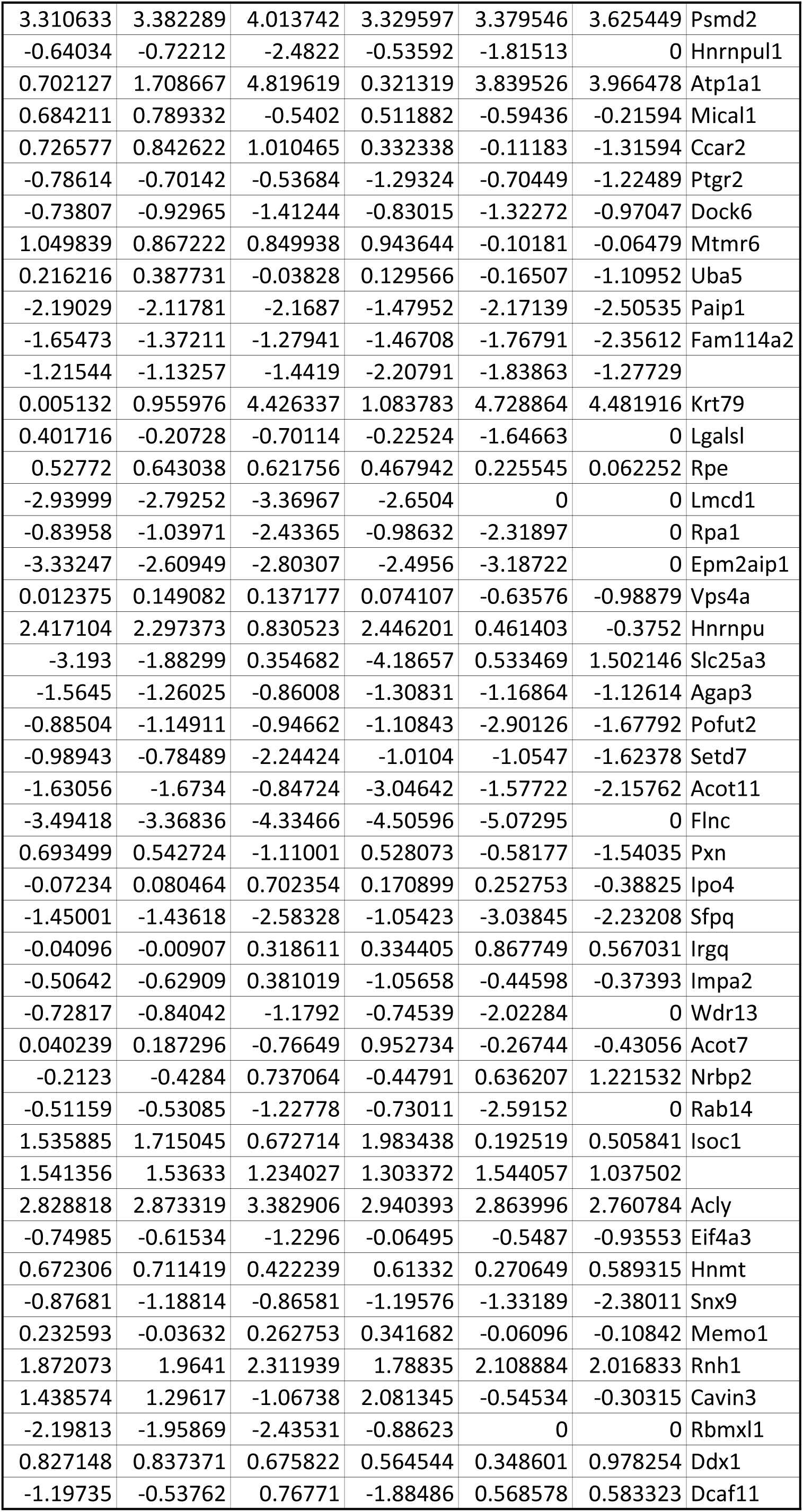
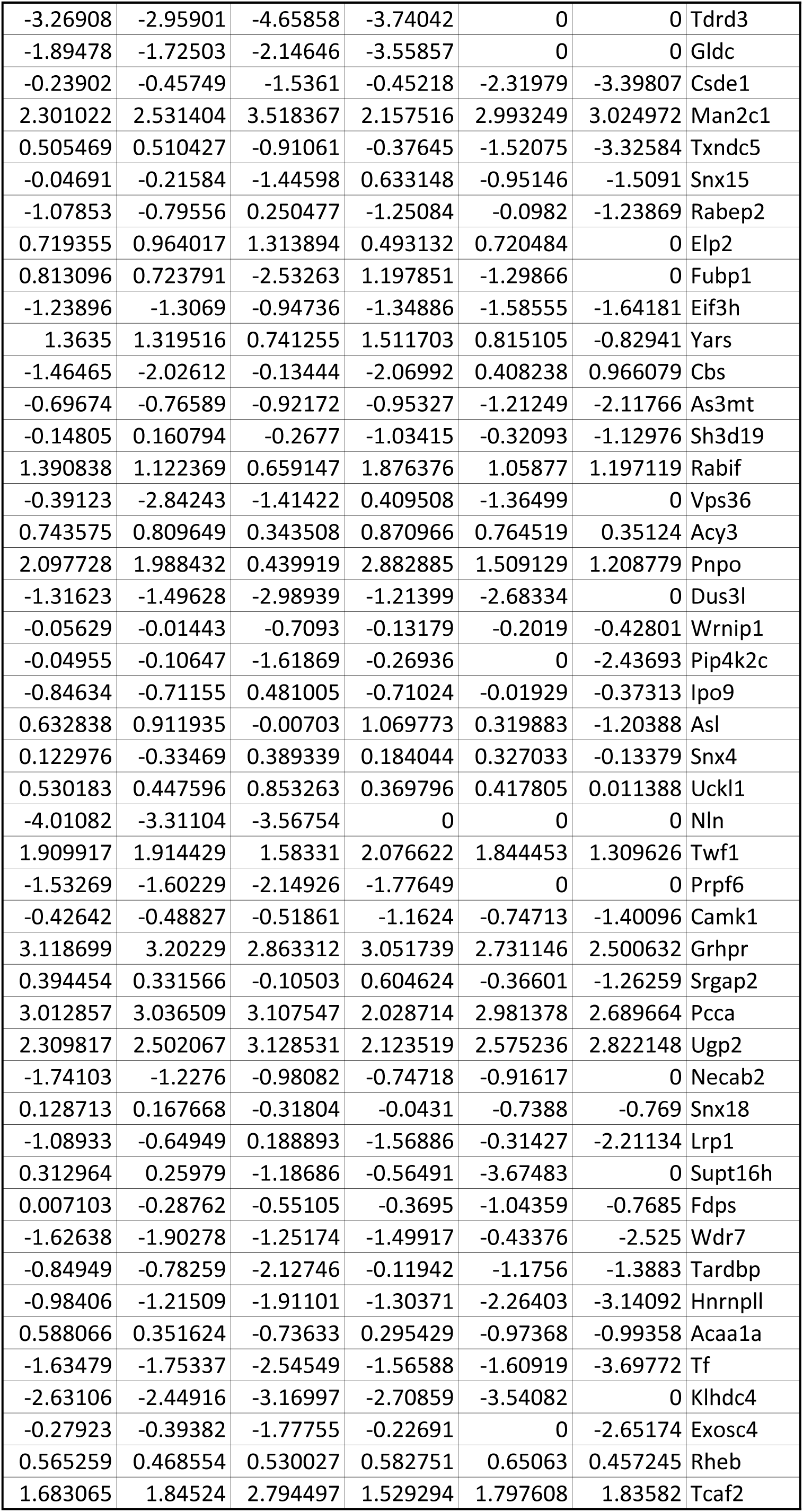
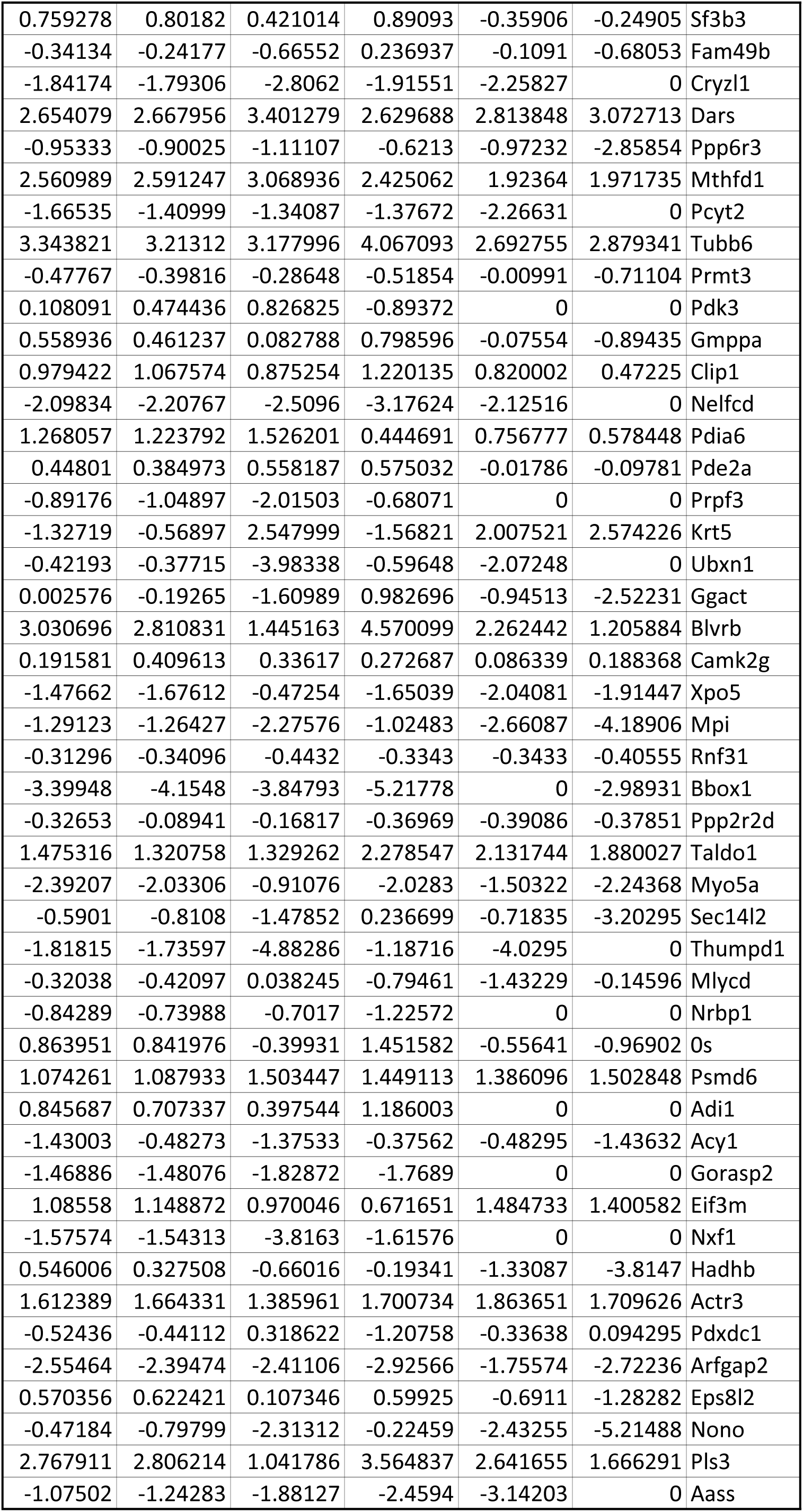
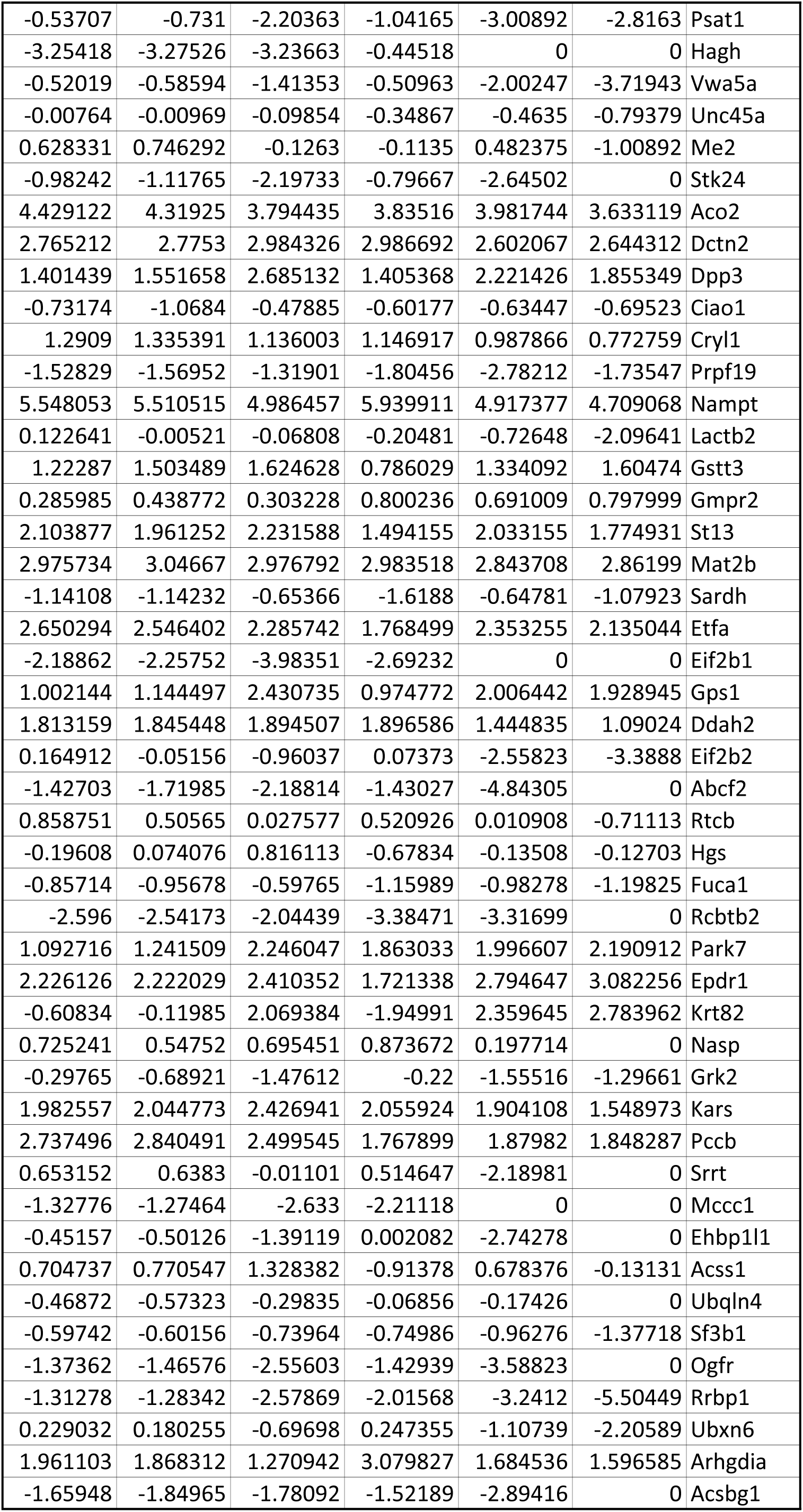
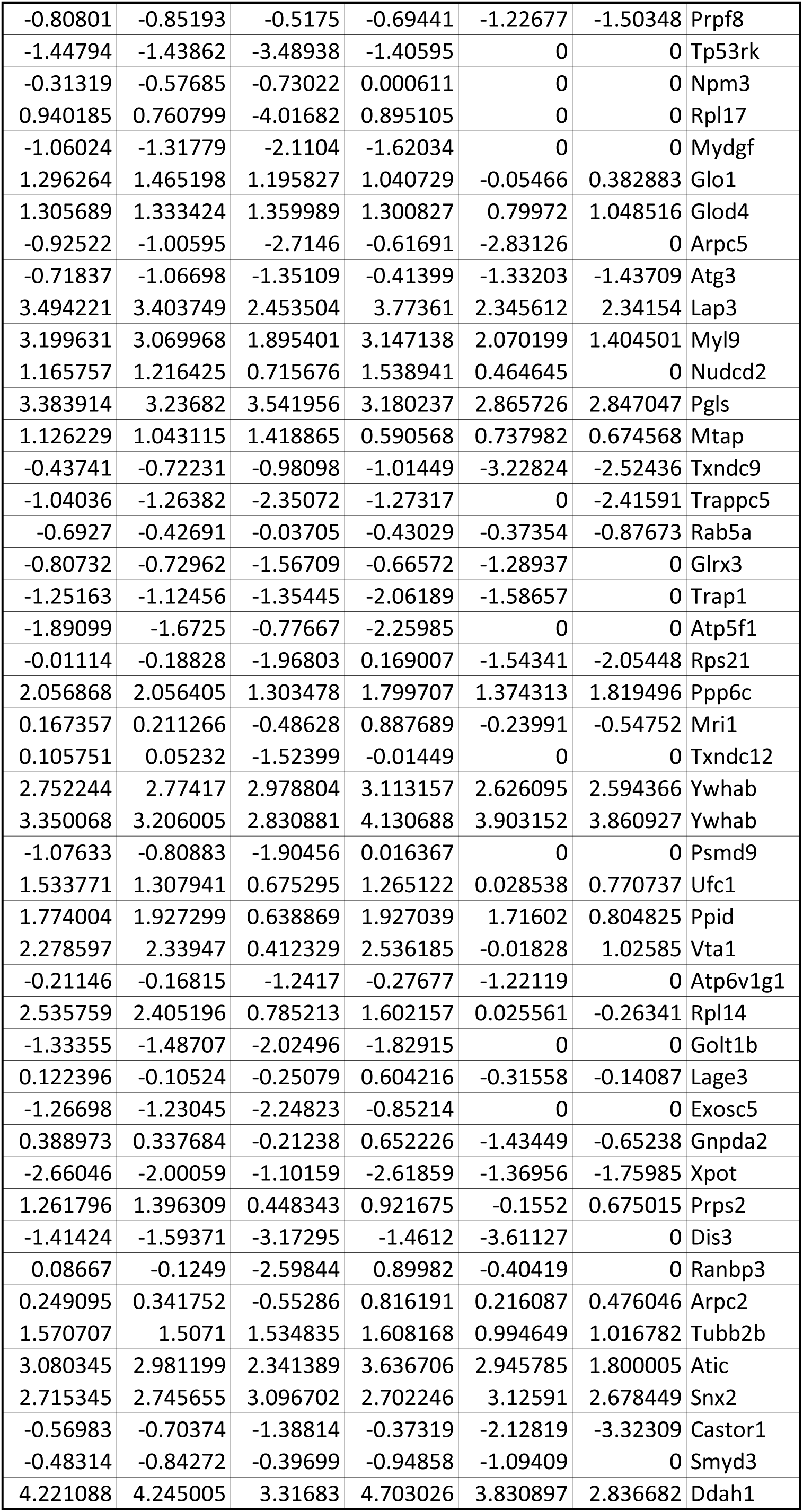
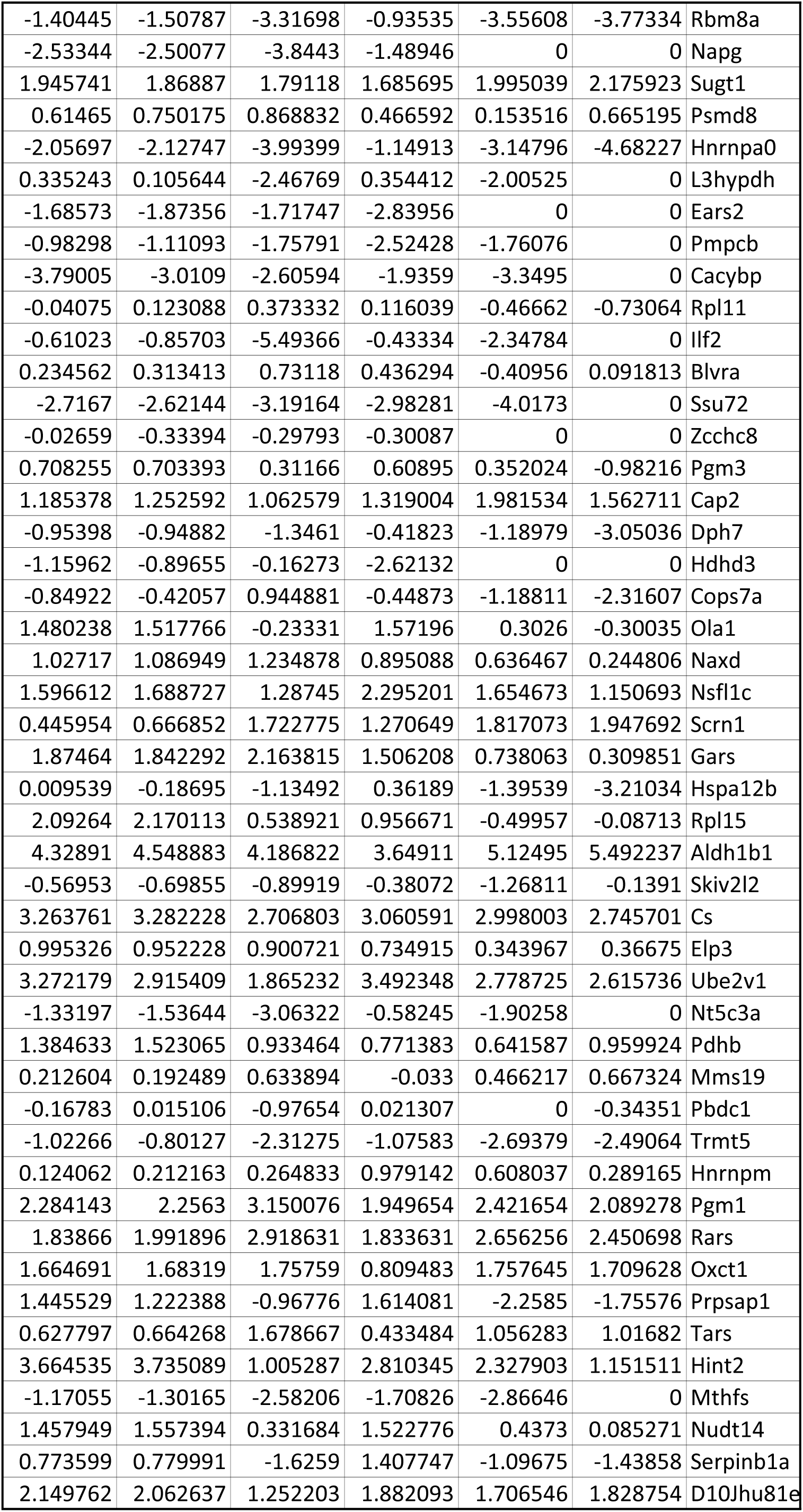
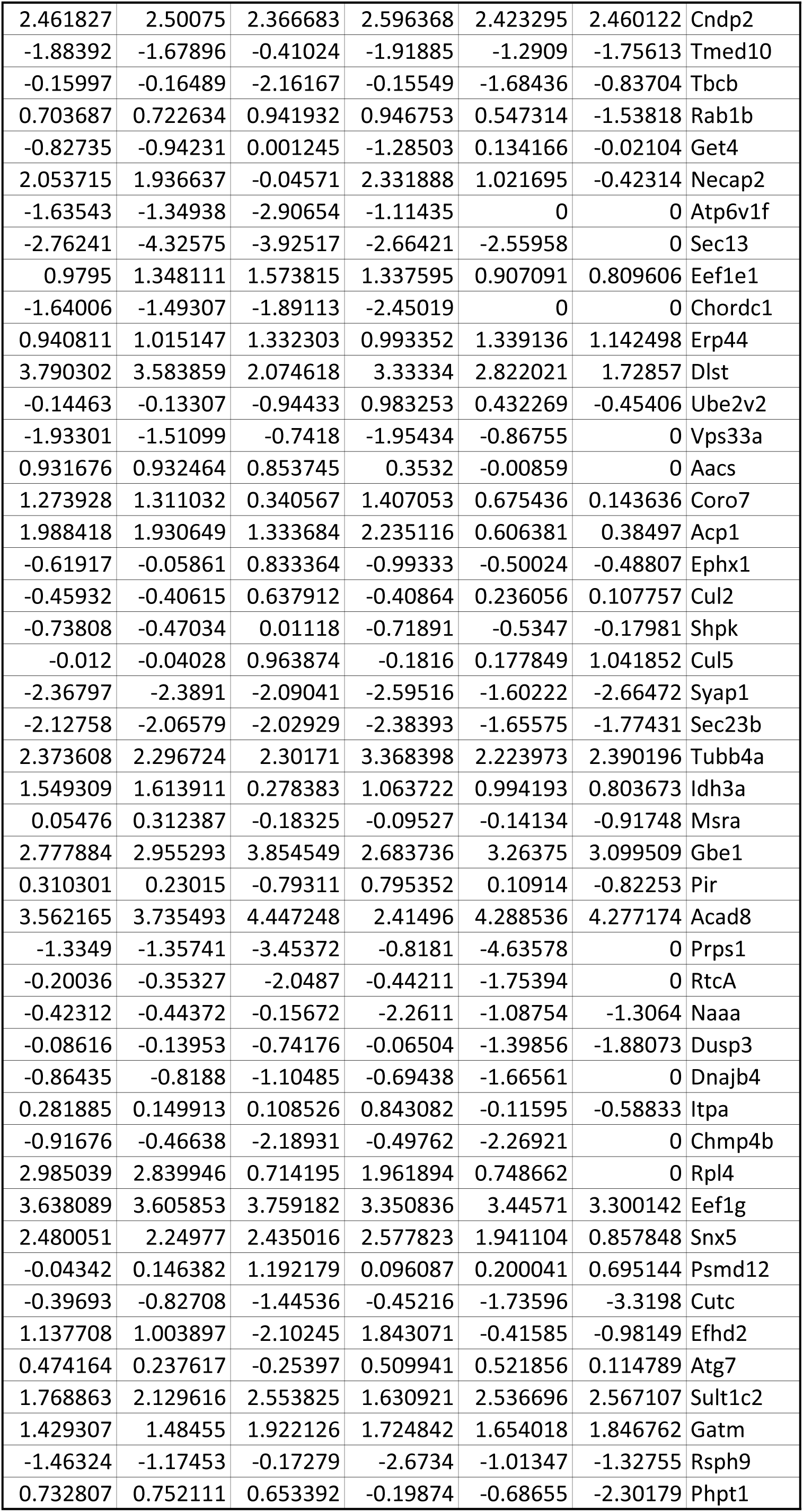
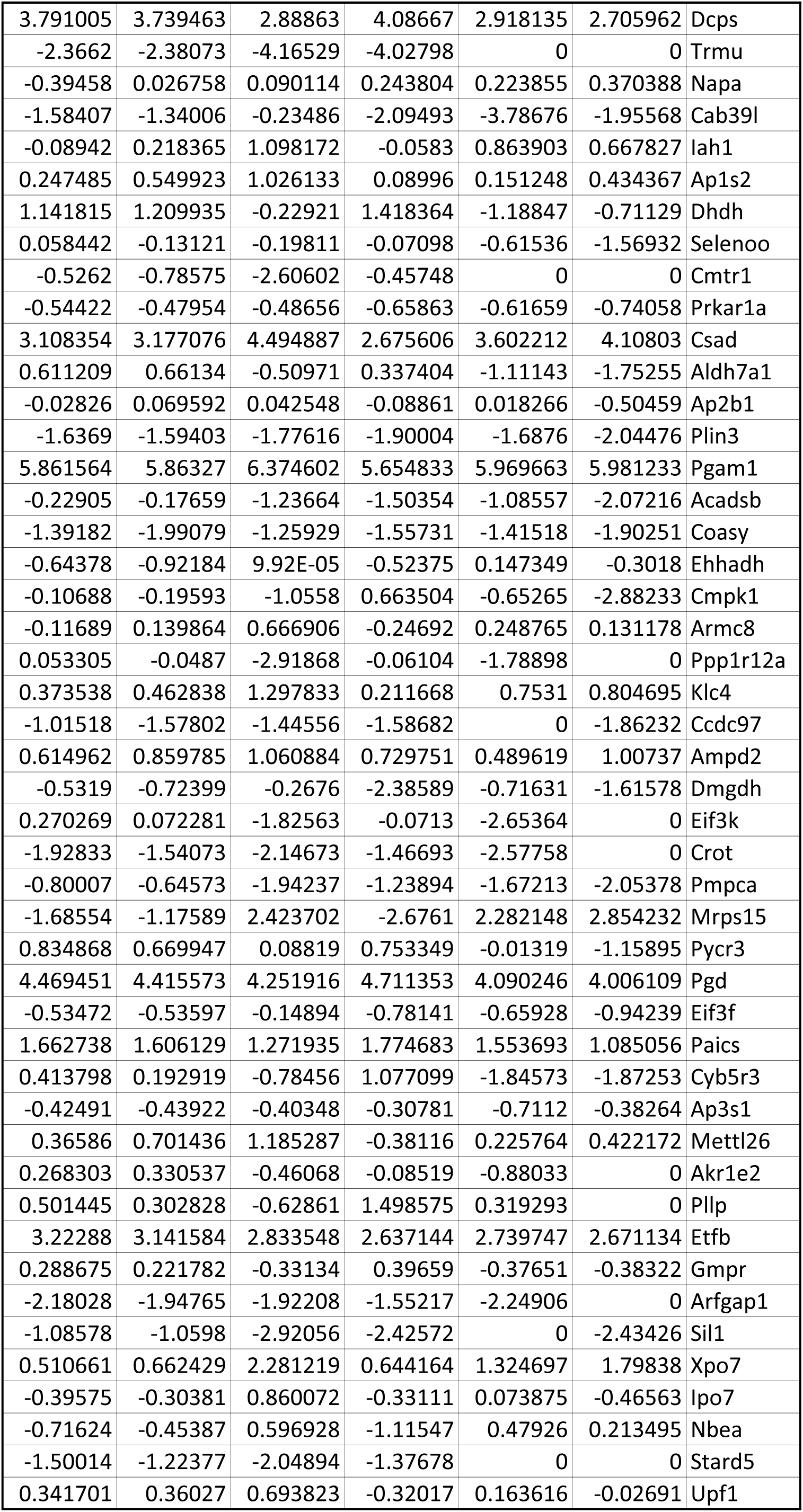
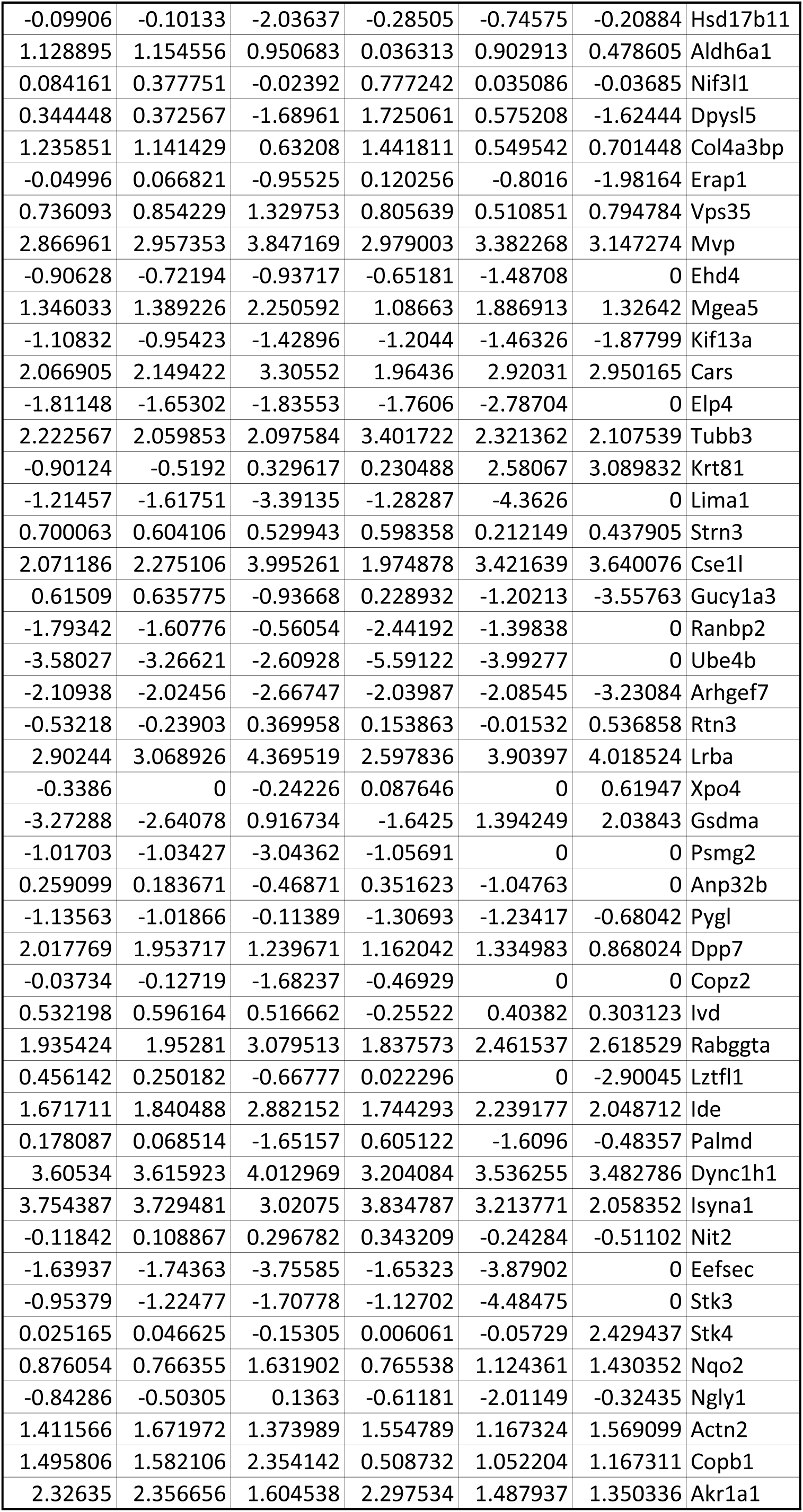
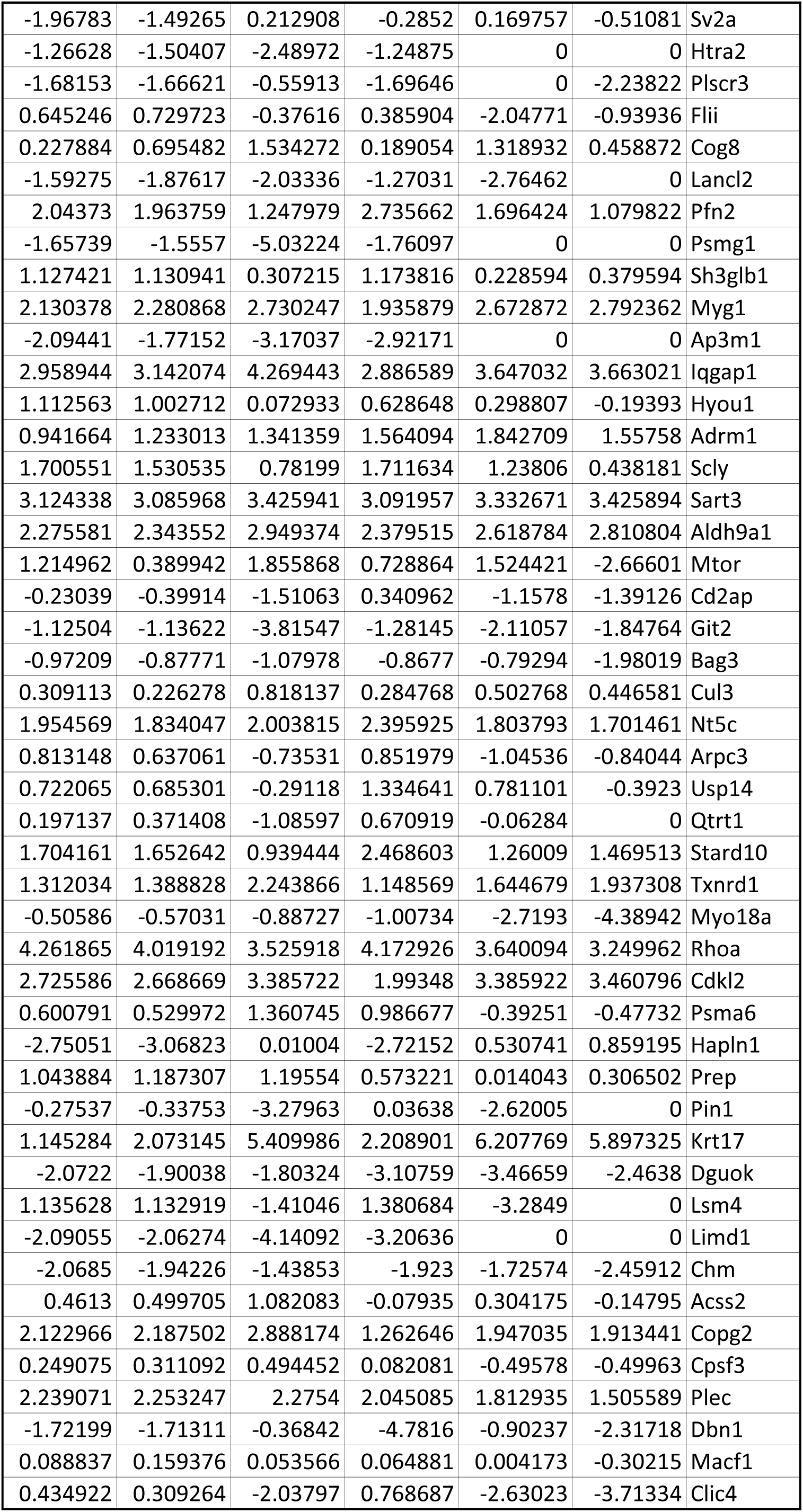
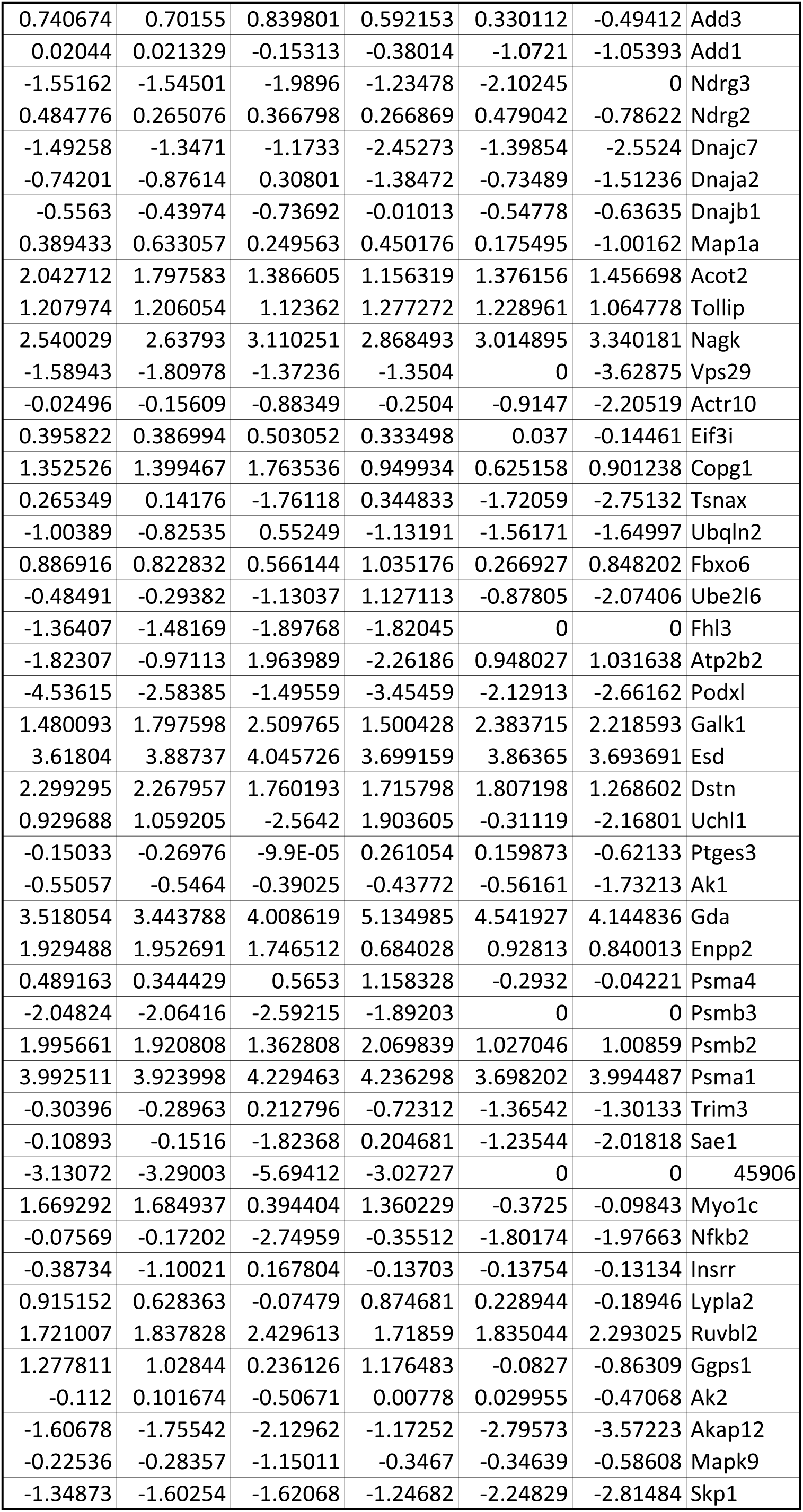
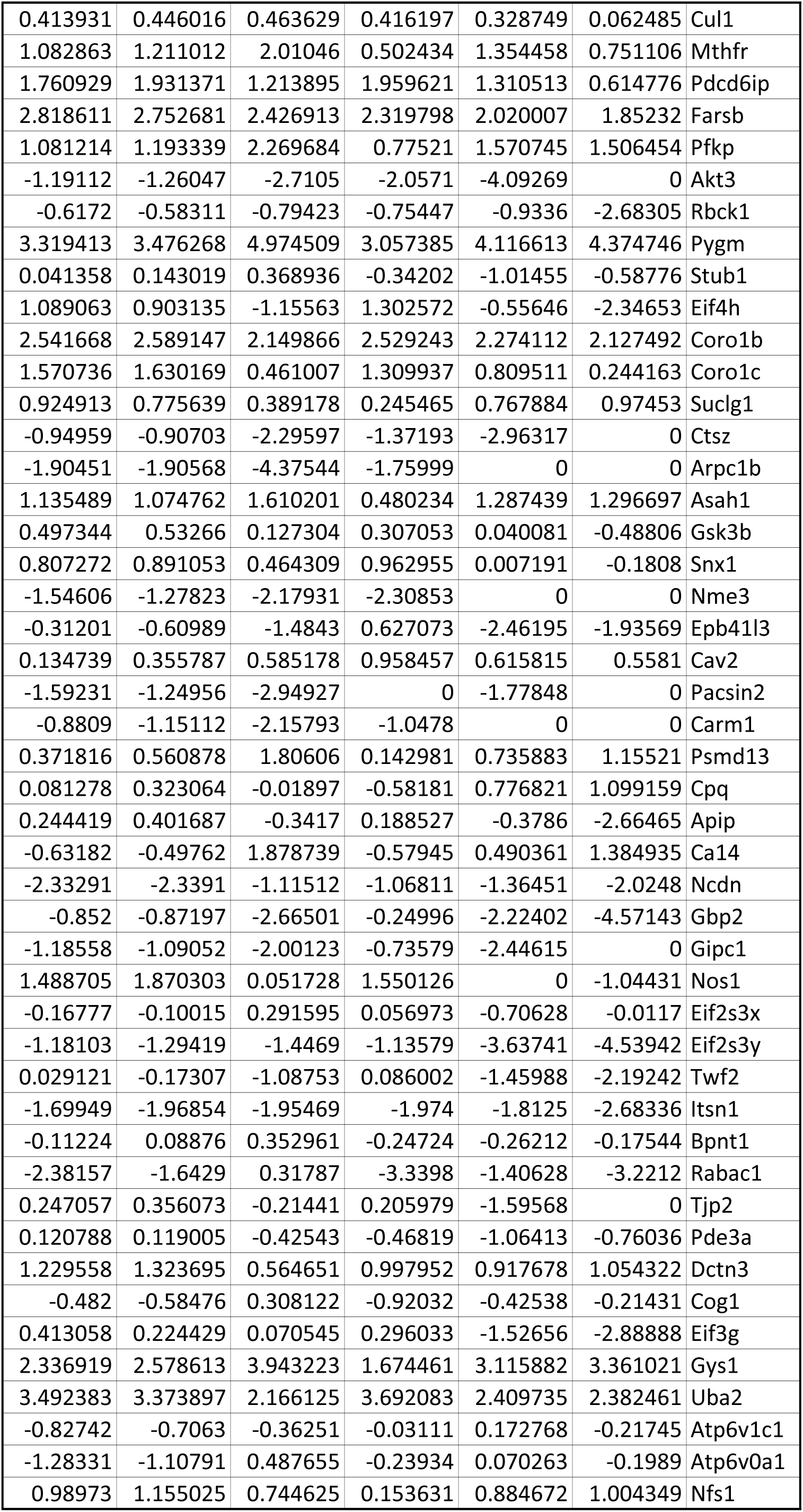
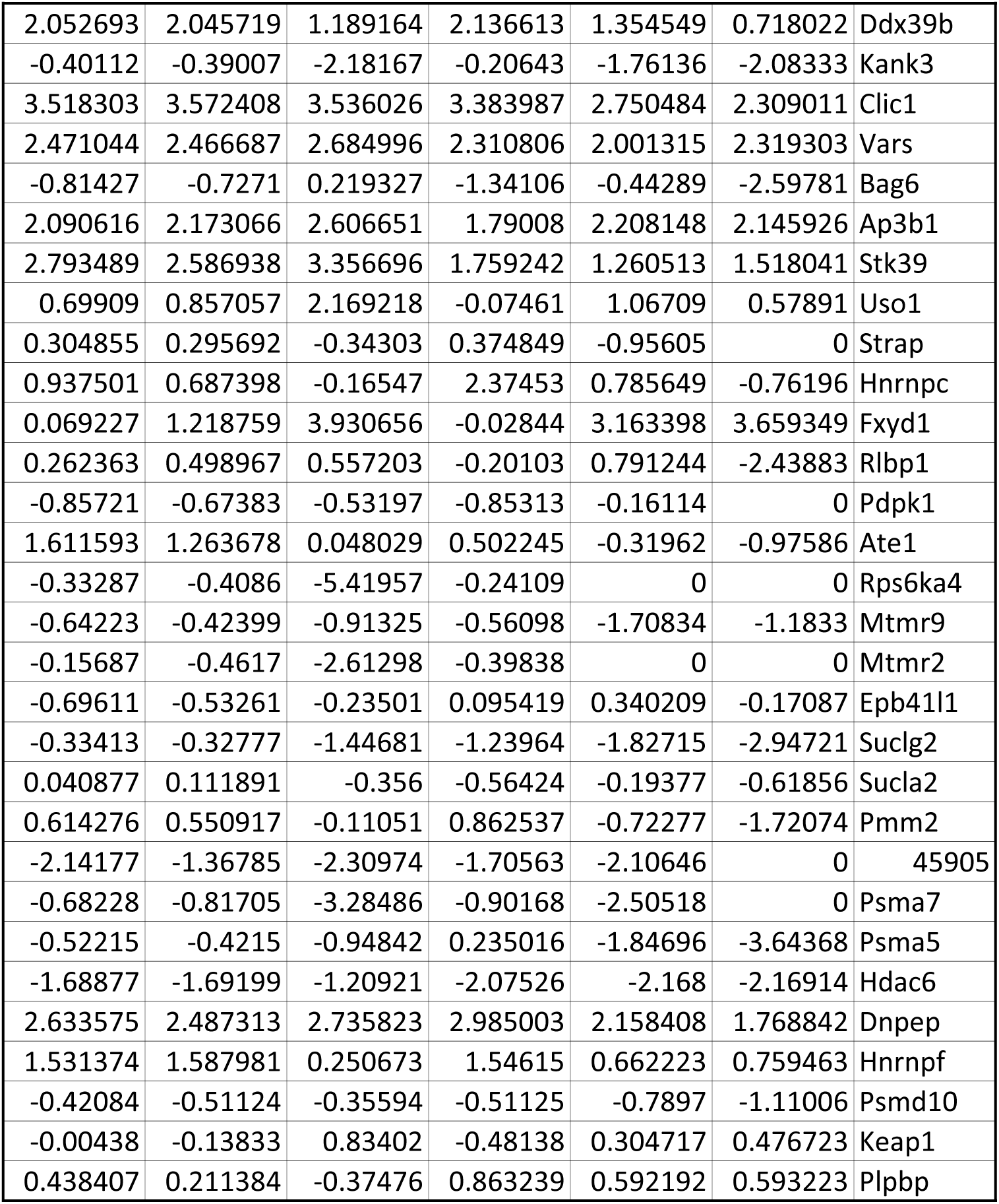
Proteonmics analysis of endosomes from ZT8 and ZT20 mice.

**Extended Data Fig. 1:**
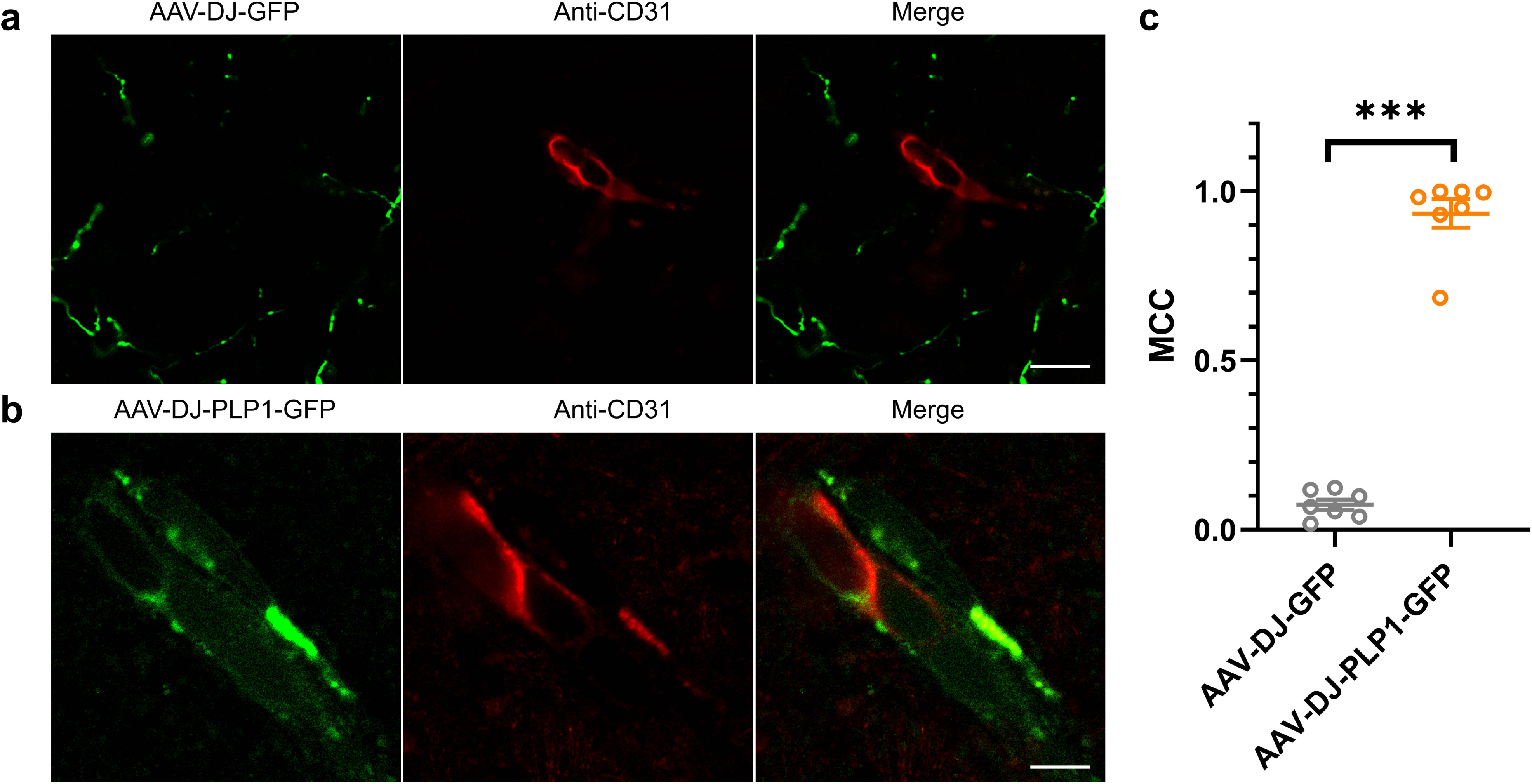
Localization of PLP1-GFP in BECs. a. AAV-DJ-GFP showed robust parenchymal expression but did not co-localize with the endothelial signal. n=7. b. AAV-DJ-PLP1-GFP localized to BECs labeled with CD31. n=7. c. Mander’s colocalization coefficient analysis of GFP or PLP1-GFP proteins with BECs. BECs, brain endothelial cells. Scale, 5 μm. \*\*\**P*<0.001; \*\**P*<0.01; \**P*<0.05.

**Extended Data Fig. 2:**
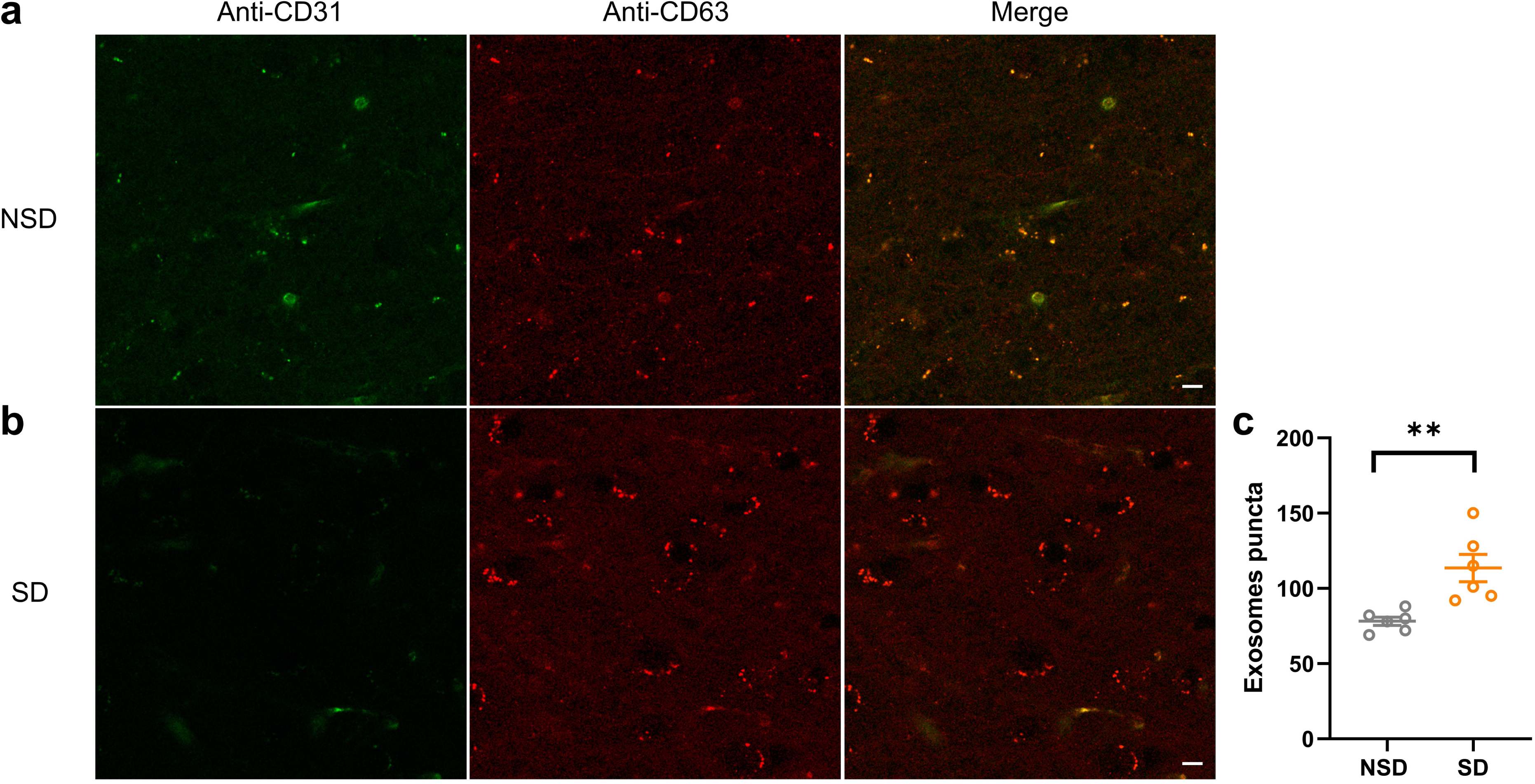
Exosomes are increased in the brain after sleep deprivation. a. Extracellular vesicles in the non-sleep deprived brain slice. CD63 is a marker for exosomes, while CD31 marks the BBB. b. Extracellular vesicles following 6 hours sleep deprivation. C. Puncta counts in A and B. n=6. Scale, 10 μm. \*\*\**P*<0.001; \*\**P*<0.01; \**P*<0.05

**Extended Data Fig. 3:**
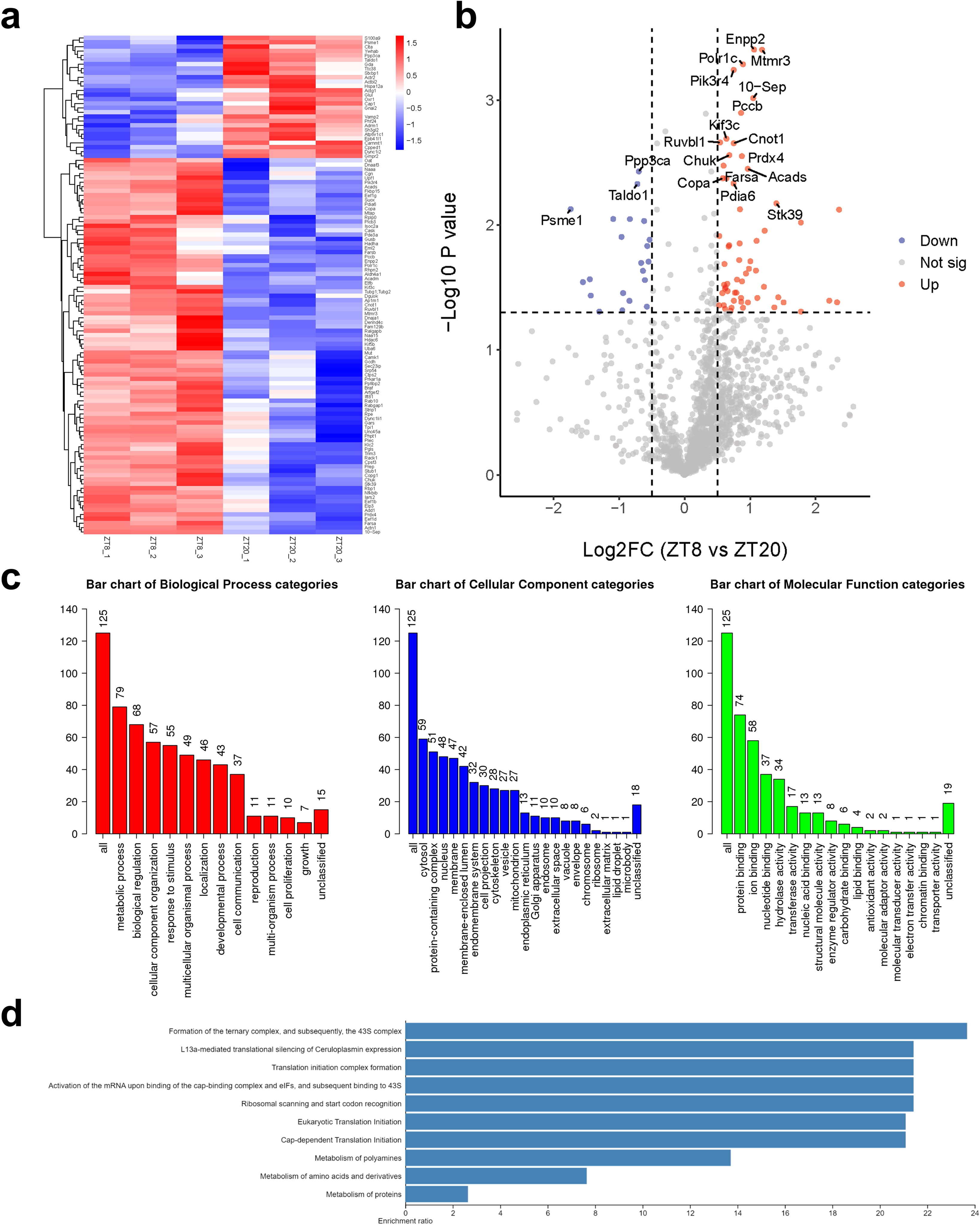
Proteomic analysis of endosomes isolated from brain endothelial cells at sleep (ZT8) and wake (ZT20) times. a. Heat maps of relative protein levels in endosomes isolated from brain endothelial cells at ZT20 and ZT8. n=3 for each group. b. Volcano plot showing proteins expressed differentially between the sleep phase (ZT8) and wake phase (ZT20). Black dots represent the significance thresholds of an applied t-test p-value = 0.05. Details are shown in Table and the Supplementary Information. c. Results of GO analysis produced bar charts of relevant biological processes, cellular components and molecular functions. d. The Reactome pathway analysis for proteins affected significantly in a. \*\*\**P*<0.001; \*\**P*<0.01; \**P*<0.05

**Extended Data Fig. 4.**
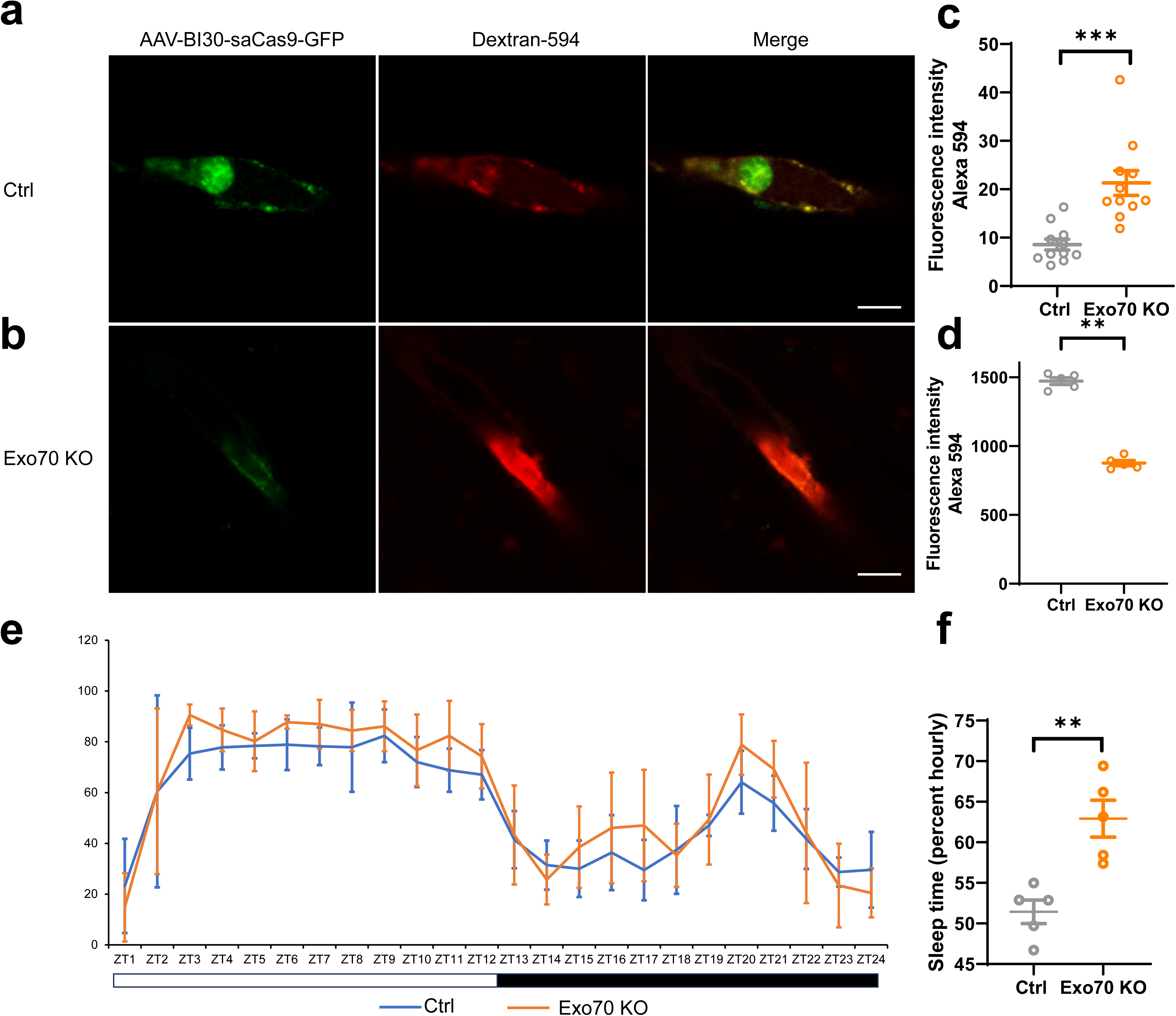
Knockout of Exo70 in the BBB decreases exocytosis and increases sleep. a. Imaging of injected Dextran Alexa Fluor 594 in BECs of control mice. b. Imaging of injected Dextran Alexa Fluor 594 in BECs of Exo70 KO mice. c. Fluorescence intensity of dextran in the brain endothelial cells of experimental and control groups. d. Fluorescence intensity of dextran 594 in the blood of control and Exo70 KO mice. e. Knock out of Exo70 in the BECs increases sleep in mice. f. Quantification of sleep time in control and Exo70 KO mice. Scale, 5 μm. \*\*\**P*<0.001; \*\**P*<0.01; \**P*<0.05

**Extended Data Fig. 5.**
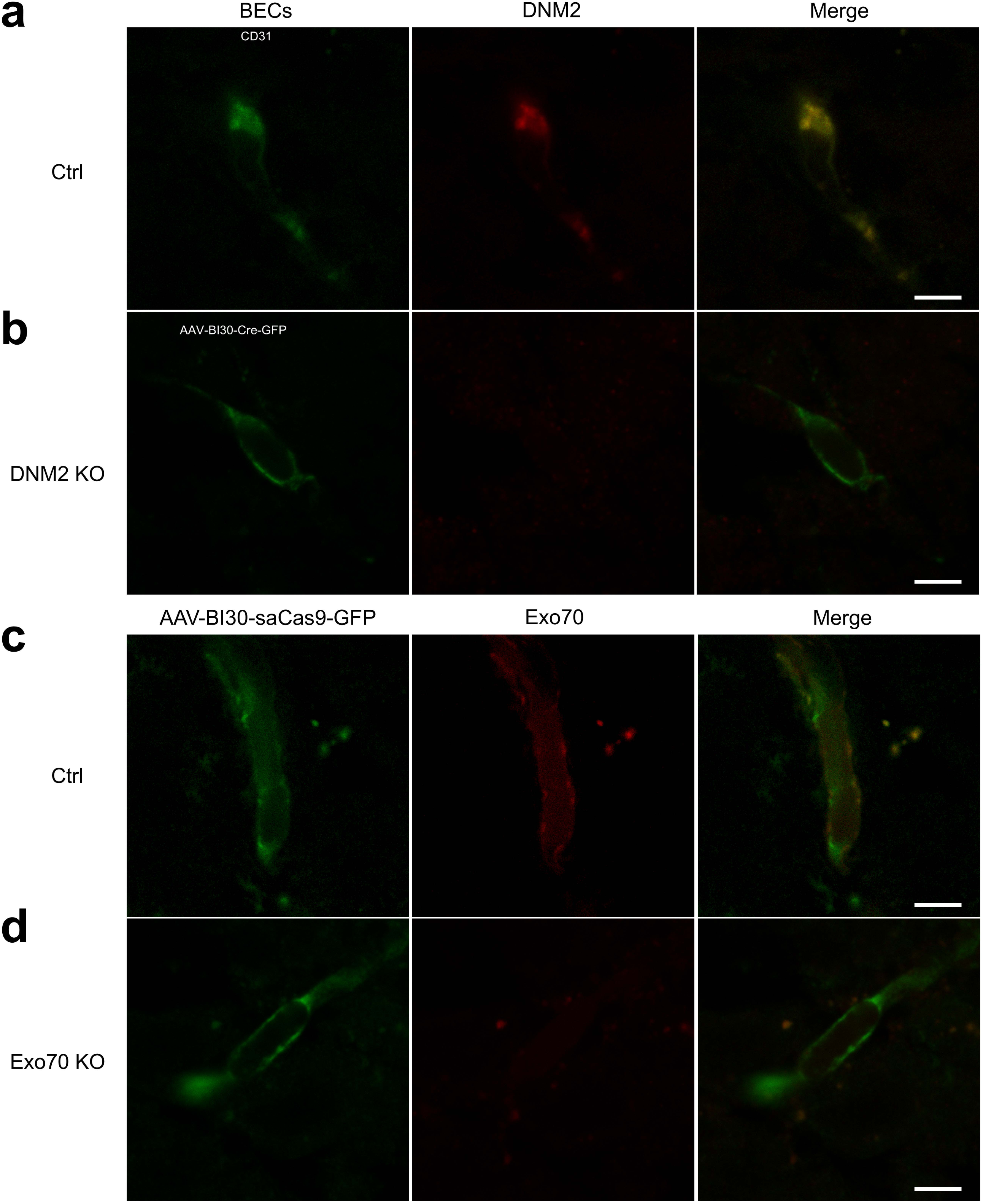
AAV-mediated knockout of DNM2 and Exo70 in brain endothelial cells (BECs). a. Immunostaining of DNM2 in BECs from control DNM2^flox/flox^ mice. b. Immunostaining of DNM2 in BECs from AAV-BI30-Cre-GFP mediated DNM2 knockout mice. c. Imaging of Exo70 in BECs of control mice. d. Imaging of Exo70 in BECs of Exo70 knock out mice. DNM2, Dynamin 2. Scale, 5 μm. \*\*\**P*<0.001; \*\**P*<0.01; \**P*<0.05

